# Combinatorial interpretation of BMP and WNT allows BMP to act as a morphogen in time but not in concentration

**DOI:** 10.1101/2022.11.11.516212

**Authors:** Elena Camacho-Aguilar, Sumin Yoon, Miguel A. Ortiz-Salazar, Aryeh Warmflash

## Abstract

Secreted morphogen signals play a key role in the determination of cell fates during embryonic development. BMP signaling is essential for mammalian gastrulation, as it initiates a cascade of signals that controls the self-organized patterning of the three germ layers. Although morphogen signals are typically thought to induce cell fates in a concentration-dependent manner, development is a highly dynamic process, so it is crucial to understand how time-dependent signaling affects cellular differentiation. Here we show that varying the duration of BMP signaling in human pluripotent stem cells (hPSCs) leads to either cells remaining pluripotent, or differentiating to mesodermal or extraembryonic states, while varying the concentration does not cause efficient mesodermal differentiation at any dose. Thus, there is a morphogen effect in time but not in concentration, and an appropriately timed pulse of BMP induces hPSCs to a mesodermal fate more efficiently than sustained signaling at any concentration. Using live cell imaging of signaling and cell fate reporters together with a simple mathematical model, we show that this effect is due to a combinatorial interpretation of the applied BMP signal and induced endogenous WNT signaling. Our findings have implications for how signaling pathways control the landscape of early human development.

## Introduction

In mammalian development, gastrulation is the first differentiation event of the embryo proper (Arnold and Robertson, 2009; Bardot and Hadjantonakis, 2020), where the pluripotent epiblast differentiates into the three germ layers of the embryo: ectoderm, mesoderm, and endoderm. This process is orchestrated by a cascade of signals initiated by BMP signaling from the extraembryonic tissues, which triggers Wnt and Nodal signaling in the epiblast (Ben-Haim et al., 2006; Brennan et al., 2001; Tortelote et al., 2013). The ligands for all three pathways find their highest expression in the proximal posterior portion of the embryo, where the primitive streak emerges, defining the anterior-posterior axis of the embryo (Conlon et al., 1994; Liu et al., 1999; Winnier et al., 1995). However, how cell differentiation is combinatorially controlled by this cascade of signals at this stage is still not completely understood.

Technical and ethical issues prevent the study of these questions *in vivo*, especially in the human embryo. To overcome this problem, several *in vitro* models of mammalian gastrulation have been developed that can serve as proxies for investigating these stages of development (Camacho-Aguilar and Warmflash, 2020; Fu et al., 2021; Heemskerk, 2020; Shahbazi et al., 2019). In particular, we and others have shown that human pluripotent stem cells (hPSCs) grown in colonies of precise size and shape using micropatterning technology and treated with the upstream gastrulation-inducing signal BMP4 pattern into all three germ layers plus a fourth one transcriptionally similar to extraembryonic tissues which most likely represents the amnion (Chhabra and Warmflash, 2021; Chhabra et al., 2019; Etoc et al., 2016; Minn et al., 2020; Warmflash et al., 2014). While some studies have suggested that these fates may emerge from differential concentrations of BMP generated by diffusion and interactions with the inhibitor Noggin (Tewary et al., 2017), several studies contradict this idea.

First, in colonies of sizes of 10 cells or smaller, BMP acts as a switch controlling a cell state transition from pluripotency to extraembryonic as BMP concentration increases without inducing mesendodermal fates at any dose (Nemashkalo et al., 2017). In fact, primitive streak induction only occurs when the cell density is above a threshold, and it is blocked if either the Wnt or Nodal pathways are inhibited (Chhabra et al., 2019; Nemashkalo et al., 2017). Thus, secondary signals are necessary for BMP to induce differentiation into the three germ layers. However, much remains unknown about the underlying mechanisms.

Here we studied how cell fate decisions are controlled by BMP signaling during human pluripotent stem cell differentiation *in vitro*. We investigated how hPSCs interpret the duration and concentration of the applied BMP4 signal. Our results unveiled an apparent *morphogen effect* in time, where short, intermediate, and long pulses of BMP4 signaling result in cells either remaining pluripotent, or differentiating to mesodermal or extraembryonic states, respectively. In contrast, there was no comparable morphogen effect in concentration: varying the BMP concentration does not cause comparably efficient mesodermal differentiation at any dose. Using live cell imaging of signaling reporters, we discovered that the temporal morphogen effect is controlled by combinatorial interpretation of the exogenously supplied BMP4 and the endogenously induced WNT signaling. Leveraging mathematical modeling, we uncovered the minimal requirements for the logic that controls these cell fate decisions, providing a fate map that explains the mechanism that allows BMP to act as a morphogen in time but not in concentration. Our findings have implications for how signaling pathways control the landscape of early human development and highlight the importance of time in *in vitro* differentiation protocols.

## Results

### BMP signaling produces a morphogen effect in time but not in concentration

As BMP induces extraembryonic fates directly but other fates through intermediates (Ben-Haim et al., 2006; Martyn et al., 2018; Nemashkalo et al., 2017), we speculated that the timescales for these processes may differ, and the duration of BMP treatment may be an important variable. Previous differentiation protocols in hPSCs also indicated that the timing of BMP treatment may influence the outcome (Zhang et al., 2008). We cultured hPSCs for two days, exposing them to varying durations of BMP4 treatment of a fixed concentration, 10 ng/mL, at the outset (Fig. 1A). We observed that cells that were exposed to a BMP pulse of 8 hours or less returned to the pluripotent or undifferentiated state, as marked by high SOX2, OCT4, and NANOG expression (Fig. 1B-D, S1A,B). On the other hand, BMP pulses longer than 32 hours were necessary to differentiate cells uniformly into an extraembryonic state, marked by high CDX2, ISL1, HAND1, and GATA3 expression (Fig. 1B-D, S1C-F). Strikingly, pulses of 16 hours showed a high level of mesoderm or primitive streak differentiation, marked by high BRA expression (Fig. 1B-D). These results unveiled a *morphogen effect* in the duration of BMP signaling, with short, intermediate, and long pulses resulting in pluripotent, mesodermal, and extraembryonic fates, respectively. Importantly, as ~97% of the cells differentiate to an extraembryonic state if BMP is maintained for the entire 2-day protocol, this morphogen effect is not due to heterogeneity in the initial cell culture, but to single cells interpreting the duration of the BMP signal to differentiate to either mesoderm or extraembryonic fates. That is, cells that would have adopted the mesodermal fate if the BMP was withdrawn are converted to an extraembryonic fate with continued signaling.

**Figure 1.**
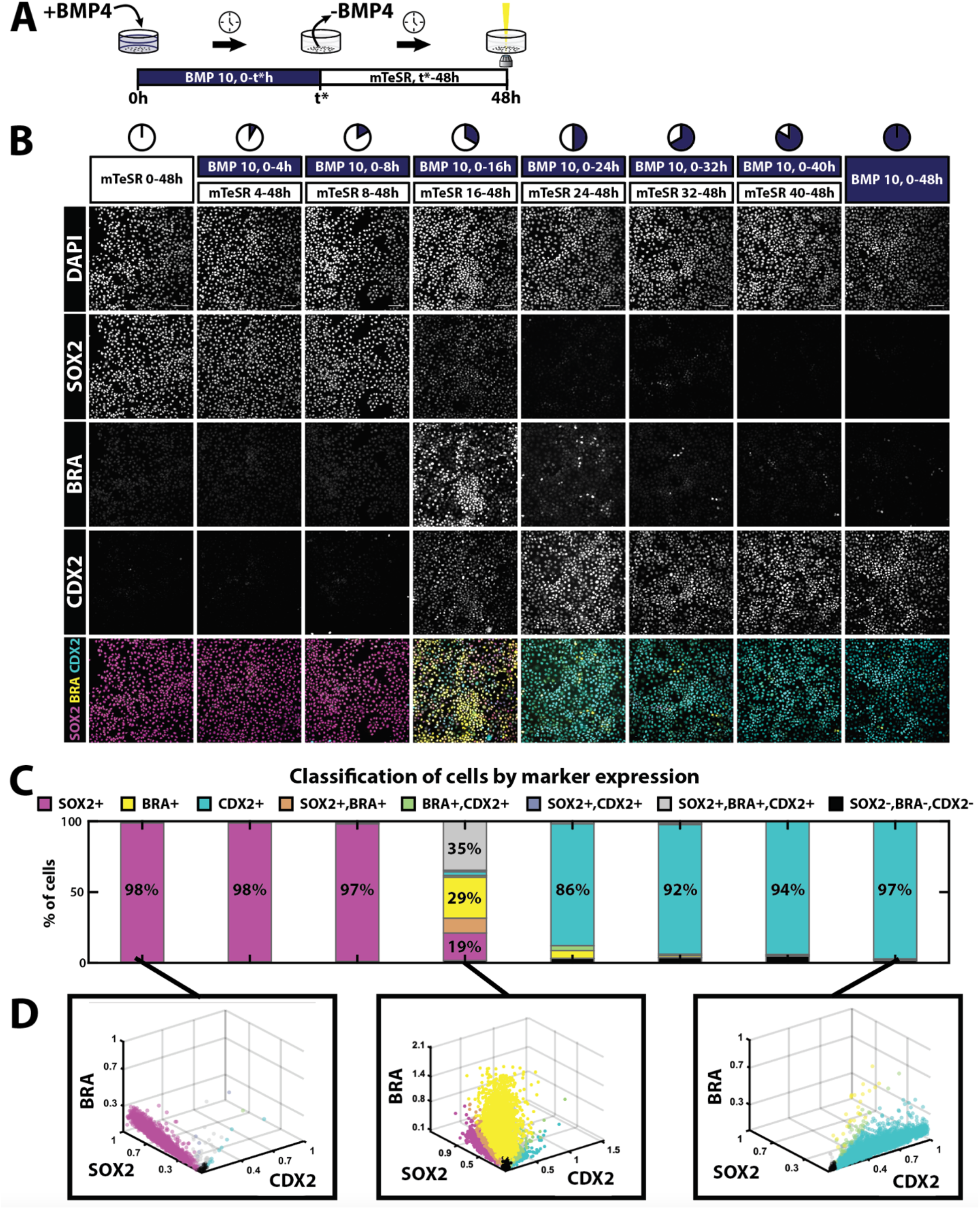
BMP4 signaling produces a morphogen effect in time. (A) Schematic of the induction protocol. hPSCs are treated with a pulse of 10ng/mL BMP4 of varying durations (t*) after which the BMP-containing media is removed and replaced by the pluripotency maintenance media mTeSR1. hPSCs are always fixed and immunostained 48h after the onset of induction. (B) Example images of immunofluorescence for DAPI, SOX2, BRA, and CDX2 after the indicated BMP4 treatments. (C) Quantifications of cell fate proportions for the experimental conditions in (B). (D) From left to right, scatter plots of the quantifications of SOX2, BRA, and CDX2 for the conditions mTeSR 0-48h, BMP 10ng/mL 0-16h followed by mTeSR 16-48h, and BMP 10ng/mL 0-48h, respectively. Each dot corresponds to a single cell, and its color marks the cell fate assigned to the cell, as shown in (C). Scale bars: 100um.

Several studies have suggested that for other pathways, including Sonic Hedgehog, duration and amplitude of signaling may be interchangeable, with shorter treatment with higher doses being equivalent to longer exposure to low doses (Dessaud et al., 2007; Sagner and Briscoe, 2017). If this was the case for BMP signaling in hPSCs, the high proportion of mesodermal cells obtained with a pulse of intermediate duration should be reproduced by treating hPSCs with a constant 2-day pulse of a lower BMP concentration. To test this, we compared the results of treating cells with 10 ng/mL BMP4 pulses of intermediate durations (ranging from 14 to 24 hours) (Fig. 2A,B), to the results obtained by inducing hPSCs with several constant concentrations (ranging from 1 to 4 ng/ml) for 48 hours (Fig. 2C,D). We observed that constant induction by any lower BMP4 concentration could not reproduce the high percentage of mesoderm cells obtained with 10ng/mL BMP4 for intermediate durations (Fig. 2A-D).

**Figure 2.**
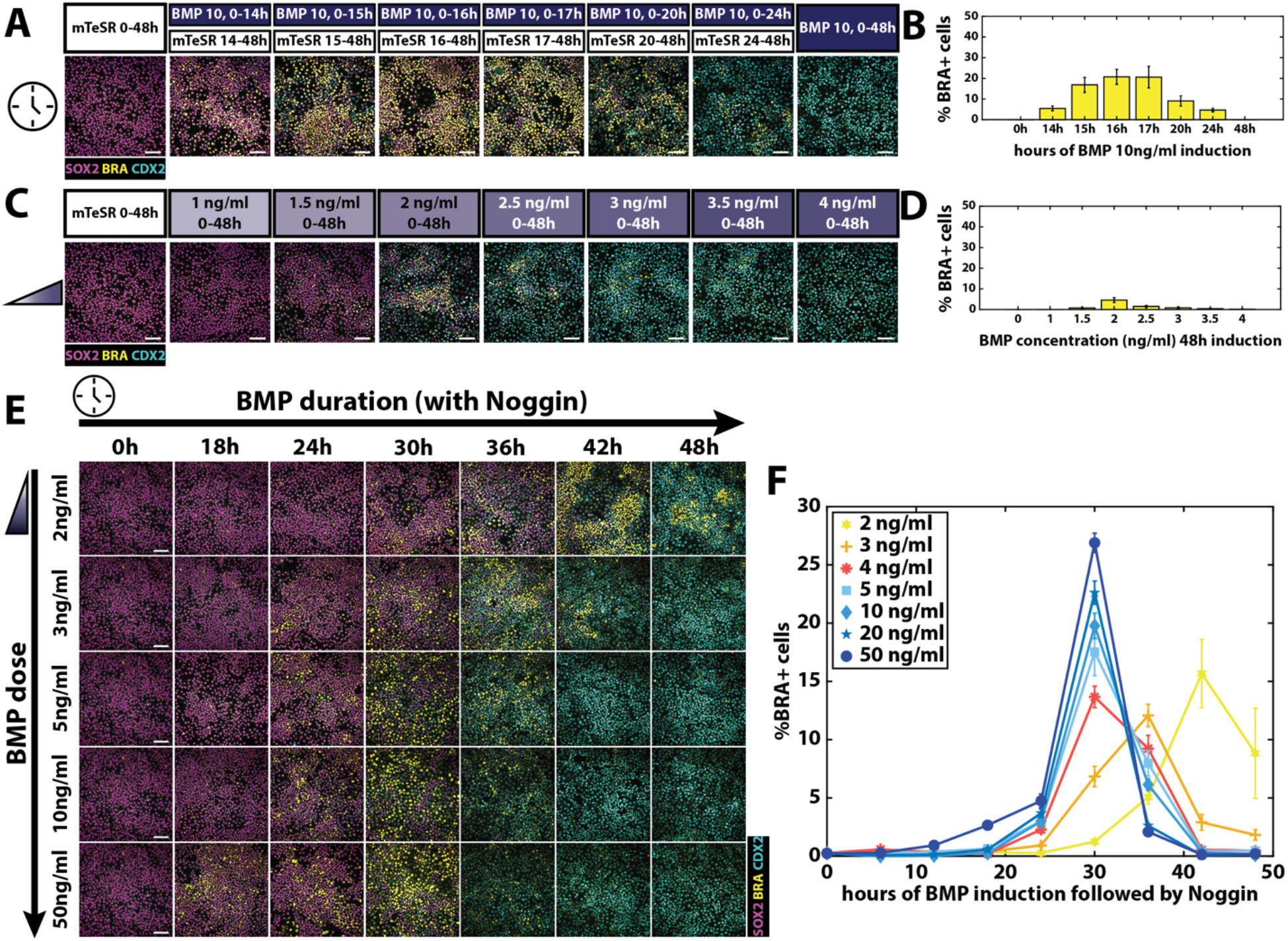
Duration and concentration of BMP4 signaling are not interchangeable. **(A)** Example images of immunofluorescence for SOX2, BRA, and CDX2 in magenta, yellow, and cyan, respectively, after the indicated pulses of 10ng/mL BMP4 treatments followed by mTeSR. **(B)** Quantifications of the proportions of cells classified as BRA+ in the treatments shown in (A). **(C)** Example images of immunofluorescence for SOX2, BRA, and CDX2 in magenta, yellow, and cyan, respectively, after the treatments of the indicated constant concentrations of BMP4. **(D)** Quantifications of the proportions of cells classified as BRA+ in the treatments shown in (C). **(E)** Example images of immunofluorescence for SOX2, BRA, and CDX2 in magenta, yellow, and cyan, respectively, after the 48h treatments with pulses of the indicated times (columns) followed by Noggin, of the indicated concentrations (rows) of BMP. **(F)** Quantifications of the proportions of cells classified as BRA+ in the treatments shown in (E). Scale bars: 100um. Error bars in (B), (D), and (F) show the SEM.

To further investigate the interplay between concentration and time of exposure to control hPSCs fate specification, we induced hPSCs with different pulses (increasing from 0 to 48 hours in 6 hours increments) of varying concentrations of BMP4 (ranging from 2 to 50 ng/mL) and observed the cell fates after 48 hours (Fig. S2A,B). Interestingly, when BMP4-containing media was withdrawn and replaced by fresh media, the data suggested that, for higher concentrations, shorter pulses were needed to induce mesoderm fates. While this result might suggest a tradeoff between time and concentration, an alternative possibility is that it could be due to only a partial shutdown of BMP signaling after higher doses of BMP are removed. This could result both from higher doses being more difficult to completely remove compared to lower ones, or activation of endogenous BMP signals at higher doses.

We therefore repeated the above experiment using a variety of different BMP durations and concentrations with a modified protocol in which BMP-containing media was replaced by Noggin-containing media. Mesoderm induction peaked with a BMP4 pulse of around 30h for nearly all concentrations except for the lowest ones (Fig. 2E,F). The later peak at lower concentrations is due to a slower induction of secondary signals necessary for differentiation, as we show below (see Fig. S3E). Notably, even though the most efficient duration of the BMP4 pulse was common for all concentrations above 3 ng/ml, quantification showed that higher concentrations yielded higher proportions of BRA-positive cells. Taken together, the results suggest that concentration and time of exposure are not interchangeable, and that efficient mesoderm induction after 48 hours of treatment required a pulse of a high BMP4 concentration.

### BMP signal induces WNT in a bistable fashion

In order to understand how signaling dynamics underlie our results on cell fate induction, we took advantage of live cell reporters of signaling activity to measure the signaling response in time under different treatment conditions. In particular, we first tracked the BMP4 response by performing live imaging of hPSCs with a GFP::SMAD4 fusion in the endogenous locus (Nemashkalo et al., 2017), and quantified the strength of the signaling response as the ratio of nuclear and cytoplasmic SMAD4 intensity (Fig. 3A-D, S3A,D). Consistent with our previous work (Heemskerk et al., 2019; Nemashkalo et al., 2017), a sudden increase in BMP4 leads to a rapid translocation of SMAD4 into the nucleus in less than 30 minutes. This response is sustained with a gradual decrease in activity over time for the remaining 48 hours if BMP4 was kept in the media (Fig. 3D, S3A). The addition of Noggin resulted in a rapid decrease in nuclear SMAD4 to baseline (Fig. 3D, S3A). Consistent with our previous observations, removal of BMP4 without Noggin addition resulted in a rapid decrease in nuclear SMAD4 but to a higher level than baseline, indicating that either some exogeneous BMP4 remains in the media or endogenous BMPs remain active once the exogenous signals are removed (Fig. 3D, S3A,D).

**Figure 3.**
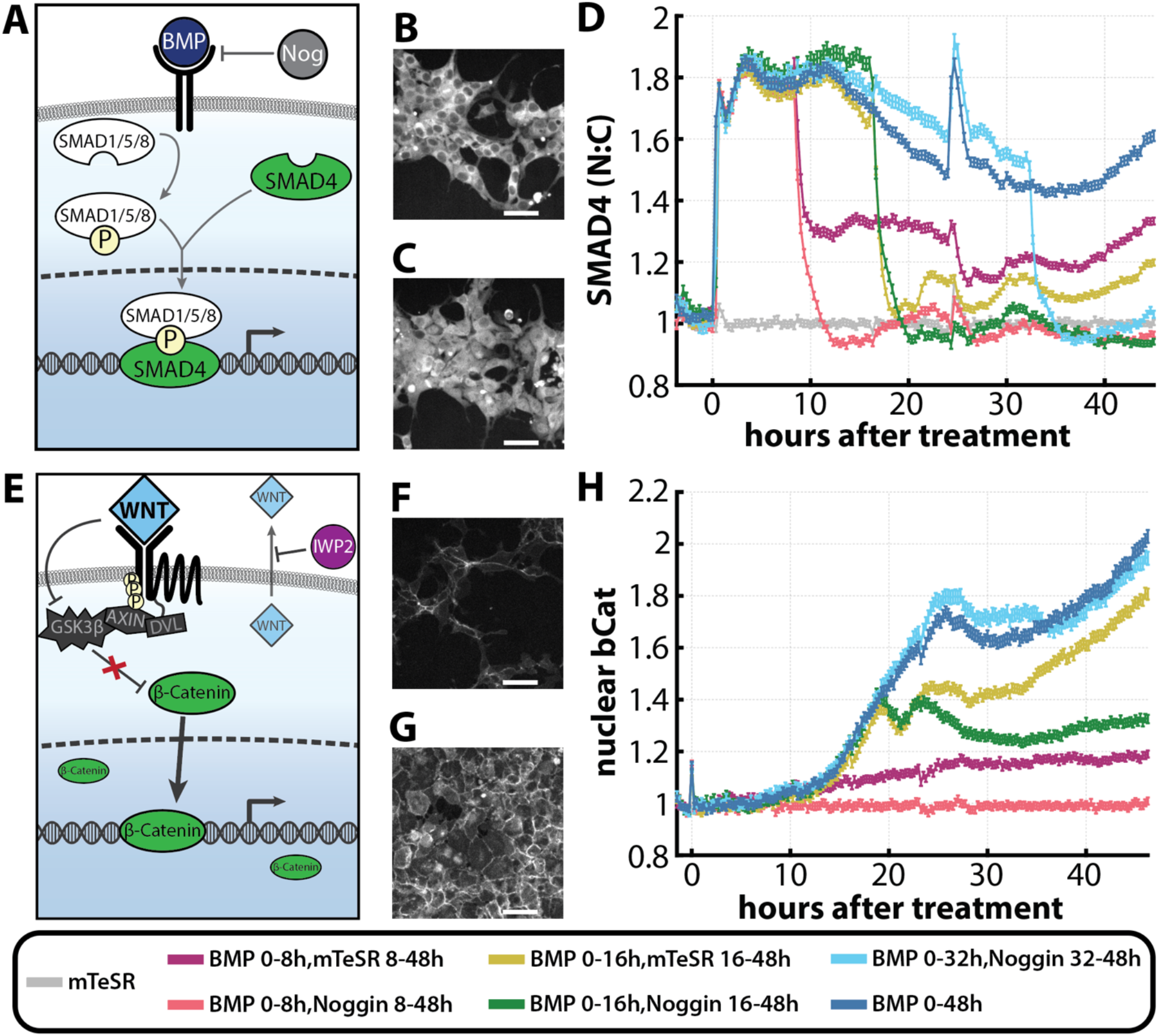
BMP signaling induces a bistable endogenous WNT. **(A)** BMP signal is transduced through SMAD4 translocation to the nucleus. Noggin inhibits BMP signal by preventing it from binding to the receptors. **(B,C)** hPSCs expressing GFP::SMAD4 before BMP4 treatment (B), and 5 hours after treatment with BMP4 (C). **(D)** GFP::SMAD4 average nuclear:cytoplasmic intensity ratio after the treatments with 10ng/ml of BMP indicated at the bottom of the figure. Additional experimental conditions are shown in Fig. S3A. **(E)** Simplified canonical WNT/β-catenin pathway. IWP2 prevents canonical WNT secretion. **(F,G)** hPSCs expressing GFP::β-catenin before BMP treatment (F), and 40 hours after treatment with BMP4 (G). **(H)** GFP::β-catenin average nuclear intensity after the treatments with 10ng/ml of BMP indicated at the bottom of the figure. Data was normalized to the BMP 0-8,Noggin 8-48h condition, as it showed similar dynamics to the mTeSR condition (Fig. S3C). Additional experimental conditions are shown in Fig. S3B-C. Scale bars: 50um.

We then explored WNT/β-catenin response, the next member of the signaling cascade that plays a role in gastrulation and mesoderm (Arnold and Robertson, 2009). We tracked WNT response by performing live imaging of hPSCs with GFP fused to the N terminus of endogenous β-catenin (Massey et al., 2019), and quantified the strength of the signaling response as the mean nuclear β-catenin intensity (Fig. 3E-H). Strikingly, measurement of Wnt dynamics revealed a bistable WNT/β-catenin response (Fig. 3H, S3B-C). If BMP was withdrawn early, β-catenin levels converged to a low level in time. However, if BMP was presented for sufficiently long, WNT activity became self-sustaining and remained stable even after BMP withdrawal. This likely results from positive feedback in which WNT signaling activates its own ligands, as has been shown in other contexts (Wang et al., 2012).

Comparing the signaling trajectories with the resulting fates reveals that only in cases where BMP is repressed after having induced WNT to a high level are cells able to differentiate to mesoderm. This suggests that a combinatorial effect between BMP and BMP-induced, bistable Wnt signals underlies the observed morphogen effect in time. Short BMP durations of high BMP concentration are not enough to activate WNT, and, therefore, cells remain pluripotent. On the other hand, if BMP is maintained for longer than 32 hours, Wnt signaling is induced but is insufficient to override the effects of BMP signaling, and cells become extraembryonic. It is in a window of middle durations, where BMP is maintained for long enough to induce Wnt autoactivation but is then suppressed, that cells are exposed to a low BMP, high Wnt signaling profile for a sufficient window of time, and mesoderm differentiation is obtained.

We also investigated how WNT/β-catenin response depended on cell density, as previous studies have shown that cell density is critical for BMP signal to induce mesodermal fates (Nemashkalo et al., 2017). Consistent with these studies, we observed that if we reduced the number of cells seeded from 40K cells/cm^2^ to 15K cells/cm^2^, pulses of BMP of high concentration could no longer induce mesoderm (Fig. S4A-D). To understand whether the lack of mesoderm correlated with reduced WNT signaling, we compared the WNT response of cells seeded in a density of 40K cells/cm^2^ (high density) versus cells seeded in a density of 15K cells/cm^2^ (low density) by performing live imaging of the β-catenin reporter cells (Fig. S4E). Indeed, we observed that the WNT response under BMP treatment was lower at low densities. While BMP induced a high WNT response in cells seeded at high density, the WNT response of cells seeded at low density under the same BMP treatment was comparable to non-treated cells at high density (Fig. S4E). Moreover, treating cells with IWP2 at the time of BMP treatment to inhibit the secretion of endogenous Wnt ligands suppressed mesoderm induction (Fig. S5). Finally, we investigated how the WNT response depended on BMP concentration and observed that the rate of increase of nuclear β-catenin increased as the BMP dose increased (Fig. S3E). Taken together, these results suggest that BMP induces a bistable endogenous WNT signal in a concentration-dependent manner, which is necessary for mesoderm induction. The mechanism behind this process will be discussed further below.

### A simple mathematical model reproduces the observed WNT bistability

To better understand the observed WNT bistability, we developed a minimal mathematical model that recapitulated the signaling dynamics described above. This model comprises two sub-systems of differential equations that model the BMP and Wnt responses, respectively, and which are connected via BMP4 activation of Wnt, which reflects the transcriptional activation of *WNT3* by BMP signaling (Fig. 4A-B) (Kurek et al., 2015).

**Figure 4.**
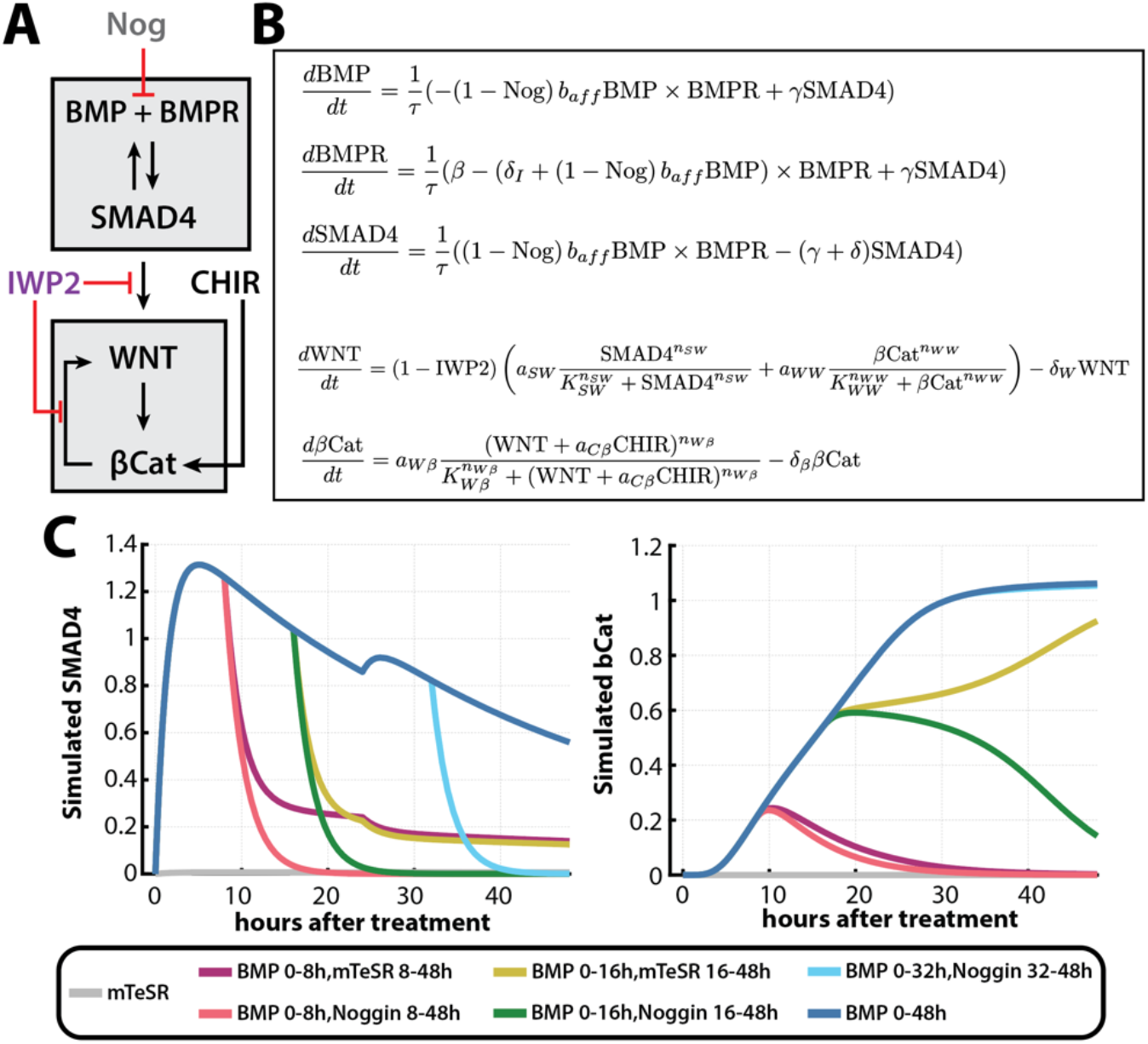
BMP-induced WNT bistability is reproduced by a simple mathematical model. **(A)** Schematic of the network regulating BMP and WNT response. **(B)** System of ODEs that model the network dynamics in (A). **(C)** Simulations of BMP and WNT dynamics under the indicated BMP4 treatments. Media change is also simulated at 24 hours for all conditions. BMP = BMP4 ligand. BMPR = BMP4 receptors. SMAD4 = Nuclear SMAD4. Nog = Noggin. WNT = WNT ligands. βCat = Nuclear β-catenin. IWP2 = IWP2.

We simulated the BMP response using the model proposed in (Heemskerk et al., 2019), where binding of BMP ligands to the receptor complex activates SMAD4 but enhances the degradation of the BMP receptor (Fig. 4A-B). The effect of the BMP inhibitor Noggin is modeled by a direct inhibition of BMP binding to free receptors (Fig. 4A-B). For simplicity, BMP withdrawal without Noggin addition was modeled by multiplying the original BMP concentration by a constant that represented the percentage of BMP that remained in the culture, as observed in Fig. 3 and S3. With this model, features of SMAD4 dynamics observed in the data such as fast SMAD4 upregulation upon BMP4 treatment, rapid SMAD4 downregulation by Noggin, and slow decay were reproduced (Fig. 4C, S6A,B).

The second sub-system models the Wnt response. Wnt expression is activated by SMAD4 and activates nuclear β-catenin. β-catenin, in turn, further activates Wnt signaling. The need to accumulate sufficient Wnt protein before β-catenin becomes active explains the delay in the rise in β-catenin activity observed experimentally (see Fig. 3H). Details of this model can be found in the Supplemental Material.

The model comprised of the two-subsystems could reproduce the observed BMP-induced WNT bistability (Fig. 4C, S6). Short BMP pulses allowed the β-catenin variable to remain in a low steady state, while longer BMP pulses resulted in the β-catenin variable converging to a high steady state where it remained even in the absence of BMP (Fig. 4C). The model also reproduced the signaling dynamics obtained under BMP withdrawal without Noggin and the BMP concentration-dependent increase of WNT signaling (Fig. S6). Interestingly, as in the experimental data, while a 16h pulse of BMP followed by Noggin inhibition resulted in β-catenin returning to a low value, a 16h pulse of BMP followed by withdrawal without Noggin resulted in β-catenin auto-activating to a high level by 48 hours (Fig. S3A-C, S6A,C). Thus, modeling supports the idea that residual BMP in the media explains why a 16h pulse of BMP 10 ng/mL is sufficient to induce mesoderm if the BMP-containing media is replaced by media alone, while longer pulses are needed if Noggin is used.

### Mesoderm induction correlates with a slow loss of pluripotency

Our next goal was to understand the dynamics of the cell fate transitions. We treated hPSCs with pulses of 10 ng/mL BMP of varying durations and examined marker gene expression at 16h, 24h, 32h, and 48h after treatment (Fig. 5A-C, S7). When cells were not exposed to BMP, cells remained pluripotent as marked by high SOX2 expression at every time point observed (Fig. 5A). On the other hand, if hPSCs were exposed to BMP for the whole 48h, a pluripotent-to-extraembryonic transition was observed between 24 and 32 hours, when cells downregulated SOX2 and then upregulated CDX2 (Fig. 5B). Importantly, no BRA expression was observed in this condition at any time point, indicating that cells transition directly to extraembryonic without going through an intermediate mesoderm state. In the experimental conditions where mesoderm differentiation was obtained, such as 16 hours of 10 ng/mL BMP4 followed by mTeSR, a slow SOX2 downregulation was observed, followed by a late BRA upregulation, starting after 32h of induction (Fig. 5C). This late mesoderm induction is consistent with recent studies that propose that SOX2 levels need to be low for Wnt to induce mesoderm differentiation (Blassberg et al., 2022, 2).

**Figure 5.**
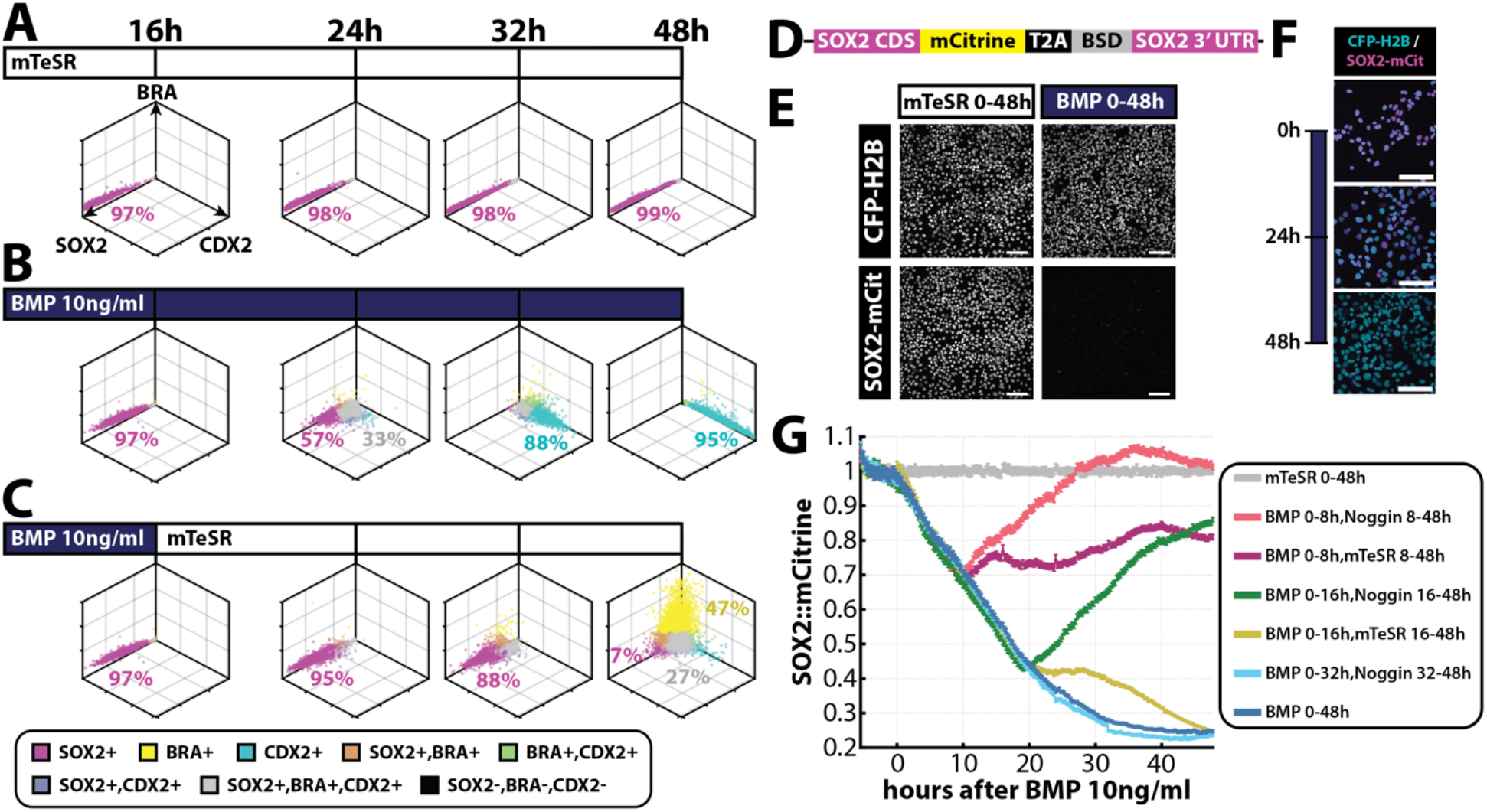
Cell fate transitions under BMP treatment. **(A-C)** Scatter plots of the quantifications of SOX2, BRA and CDX2 under the treatments mTeSR 0-48h (A), BMP 10ng/mL 0-48h (B), and BMP 10ng/mL 0-16h followed by mTeSR 16-48h (C), at 16h, 24h, 32h, and 48h. Each dot corresponds to a single cell, and its color marks the cell fate assigned to the cell as shown in the legend at the bottom. **(D)** Schematic of resulting mRNA transcribed from the labelled SOX2 allele after CRISPR-Cas-mediated SOX2-mCitrine-T2A-BSD knockin. The blasticidin resistance protein (BSD) facilitates selection of labelled cells. T2A is a self-cleaving peptide that enables separation of BSD from SOX2::mCitrine. **(E)** Representative images of hPSCs with mCitrine-labelled SOX2 and the nuclear marker CFP::H2B. (Left) Under no treatment. (Right) SOX2 expression is lost after 48h of 10ng/mL BMP4 treatment. **(F)** Confocal microscopy images of live SOX2::mCitrine hPSCs at 0h, 24h, and 48h after treatment with 10ng/mL BMP4. **(G)** SOX2::mCitrine average nuclear intensity after the indicated treatments. Scale bars: 100um.

In order to observe gene expression dynamics with a higher time resolution, we used CRISPR-Cas9 genome engineering to insert mCitrine at the endogenous locus to form a C-terminal fusion with SOX2 (SOX2::mCitrine) (Fig. 5D, S8, Materials and Methods). To facilitate nuclear identification and analysis, the cells also express CFP::H2B, which was incorporated into the genome using the ePiggyBac transposable element system (Lacoste et al., 2009). Treatment of these cells with BMP4 showed a rapid downregulation of SOX2, with undetectable expression after two days of treatment (Fig. 5E,F, S8A,B), and antibody staining for SOX2 and other pluripotency markers showed that these cells shared similar dynamics with WT hPSCs (Fig. S8).

We imaged SOX2::mCitrine cells under different BMP4 treatments and observed the SOX2 dynamics for two days. Consistent with the results above, SOX2 expression was maintained in cells that were not exposed to BMP4 (Fig. 5G). On the other hand, SOX2 expression rapidly began to decay under BMP4 treatment, with a rate that was initially proportional to BMP4 concentration (Fig. 5G, S8D). If BMP4 was withdrawn from the media early enough, SOX2 expression returned to high levels, while if cells were induced with long pulses of BMP4, they lost SOX2 expression. Interestingly, with a 16-hour pulse of 10ng/mL BMP4, which leads to peak mesoderm differentiation, SOX2 downregulation slowed after 16 hours. A second phase of SOX2 decay led to hPSCs eventually losing SOX2 expression but later than cells induced under longer BMP4 pulses (Fig. 5G). These results show that mesoderm induction correlates with a slow SOX2 decay after BMP withdrawal.

The initial decay of SOX2 expression is strongly correlated with the integral of BMP4 in time (Fig. S9A-E), suggesting that there is a tradeoff between the magnitude and the duration of signaling in the dissolution of the pluripotent state. This correlation no longer holds in later times (Fig. S9E), and the total integral of BMP4 signal does not directly translate to cell fate (Fig. S9F). This supports the hypothesis that BMP initially controls the exit from pluripotency, and the combinatorial interpretation of BMP and endogenous WNT signaling controls the decision between ExE and mesoderm fates.

Taken together, these results suggest that BMP4 concentration initially regulates SOX2 expression, which rapidly starts to decay after BMP4 treatment at a rate that is proportional to SMAD4 activity. If BMP4 is removed early, SOX2 expression recovers to baseline and cells remain pluripotent. Long BMP4 treatments produce a direct pluripotency-to-ExE transition, without going through an intermediate mesoderm state. On the other hand, intermediate BMP4 treatments produce a two-phased SOX2 downregulation, with an initial fast partial decay followed by a slow decay to baseline which correlates with the onset of mesoderm induction.

### A simple network model can recapitulate the observed dynamics

To understand how signal dynamics controlled the observed cell fate transitions, we created a minimal cell fate network (CFN) model, where the nodes of the network correspond to the cell fates observed, i.e. pluripotent, mesodermal, and extra-embryonic, characterized by high SOX2, BRA, or CDX2 expression, respectively. We coupled this CFN model with the signaling model described above (Fig. 6A). For simplicity, we modeled SOX2, BRA, and CDX2 expression in time as a proxy for the corresponding cell fates. To generate mutually exclusive cell states, we included mutually repressive interactions between SOX2, BRA, and CDX2 (Fig. 6A).

**Figure 6.**
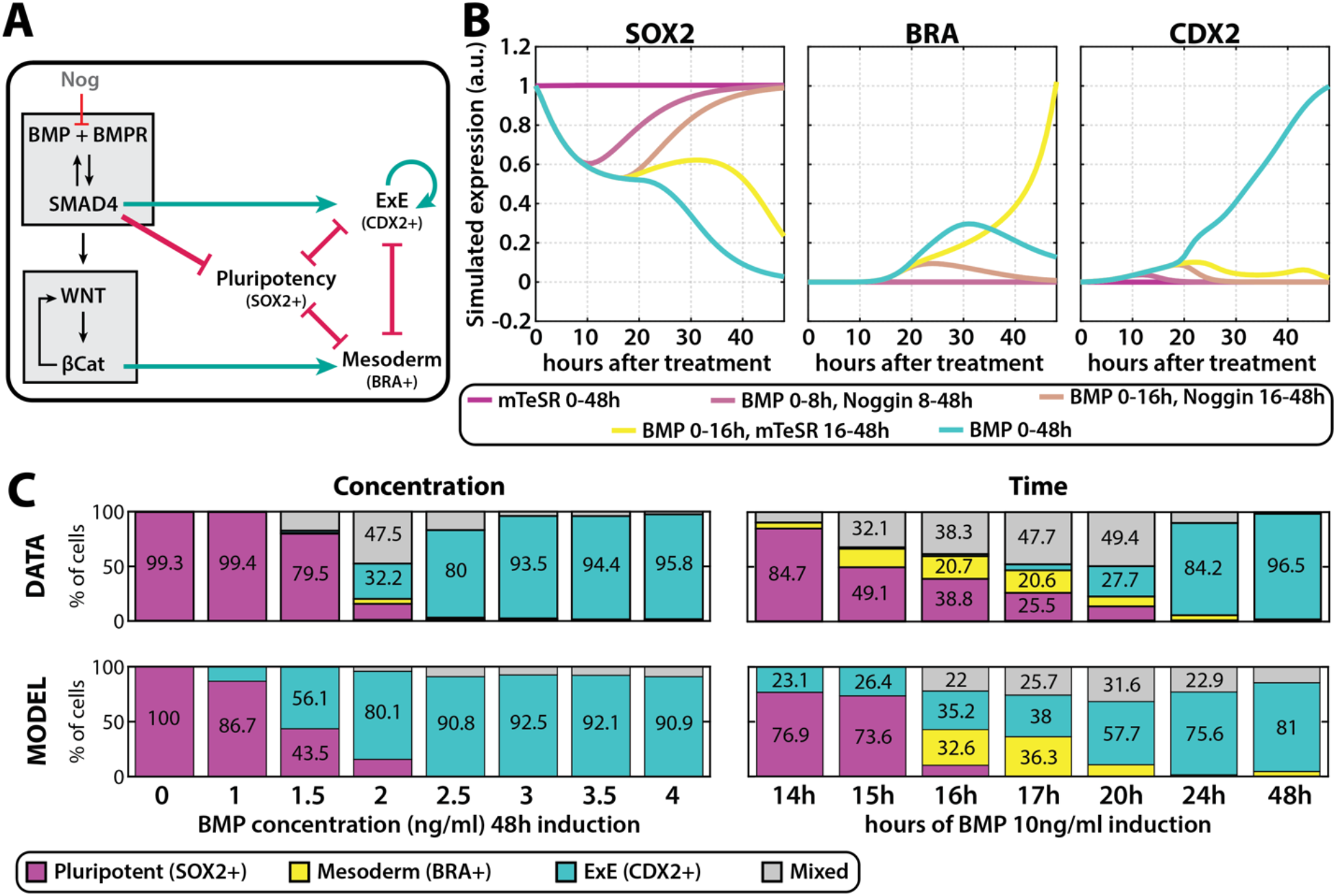
A minimal cell fate network can recapitulate the observed dynamics. **(A)** Schematics of the coupling of the signalling network and the minimal cell fate network. **(B)** Simulated SOX2, BRA and CDX2 dynamics from the model in (A) for the indicated experimental conditions. **(C)** Experimental (top) and simulated (bottom) cell fate proportions under constant low concentrations (left) and pulses of BMP 10ng/mL (right).

Secondly, as previous studies have shown and our data supports (Chhabra and Warmflash, 2021; Li et al., 2013; Minn et al., 2020; Xu et al., 2002; Yang et al., 2021), BMP activation of SMAD complexes leads to transcription of ExE markers, such as CDX2, and therefore we included activation of CDX2 expression by nuclear SMAD4 through a Hill equation (Fig. 6A, Supplemental Information). In this regard, we also observed that CDX2 and ISL1 expression levels, once active, remained high even if BMP was withdrawn from the media, suggesting a possible auto-regulation of ExE markers, which we modeled by an additive auto-activation (Fig. 6A, Supplemental Information).

As WNT signals are necessary for BMP-induced mesoderm formation (Chhabra et al., 2019; Martyn et al., 2018; Nemashkalo et al., 2017), we considered that mesoderm markers are upregulated through β-catenin activation by WNT, and modeled the activation of BRA expression through a Hill function of β-catenin levels (Fig. 6A, Supplemental Information).

Moreover, the SOX2 expression dynamics observed in Fig. 5G suggested a downregulation of SOX2 by BMP, through a SMAD intermediate, as SOX2 expression begins to decay rapidly upon BMP treatment, at which stage neither Wnt signaling, as reflected in nuclear β-catenin, nor mesoderm or ExE markers are yet upregulated (Fig. 3H, 5A-C). In fact, as mentioned above, the initial rate of SOX2 decay showed a strong correlation with SMAD4 levels for different concentrations and durations (Fig. S9). Hence, a direct inhibition of SOX2 expression levels by SMAD4 was included in the CFN model (Fig. 6A, Supplemental Information).

This minimal deterministic CFN model was fitted to simplified expression levels obtained in a subset of experiments by using a Monte Carlo optimization algorithm, the details of which can be found in the Supplemental Information. The model was compatible with the dynamics observed in the data, such as direct pluripotency-to-ExE transition without expressing BRA under constant 2-day BMP4 induction. The model was also able to mimic the slow SOX2 decay under an intermediate pulse of BMP4, which also resulted in a late BRA upregulation (Fig. 6B). Importantly, the model was able to reproduce features and conditions that were not used in the fitting, such as the observed SOX2 dynamics for different BMP4 concentrations, and that a longer pulse of 5ng/ml BMP is necessary for cells to differentiate to the ExE than under 10ng/ml BMP in followed by mTeSR withdrawal (Fig. S10).

Having fitted the deterministic CFN model to the available data, we generated a stochastic version by considering an additive white noise of strength *ξ*, to test whether the model could reproduce the cell fate proportions observed in the data. We fixed all the parameters from the deterministic model and fitted only the single noise parameter and the initial distribution of the SOX2 expression to the cell fate proportions obtained in a subset of experimental data (Supplemental Information). We found that without modification to any of the parameters from the deterministic model, the stochastic model reproduced the observed BMP morphogen effect in time and the low mesoderm induction at any low constant concentration (Fig. 6C). Taken together, the simple cell fate network model recapitulated the dynamics and cell proportions observed in the data including in a large amount of data not used to fit the model parameters.

### Cell fate is a combinatorial response to BMP and WNT

With the obtained model, we then aimed to understand how the signals BMP and WNT controlled cell fate determination. We created a phase diagram, or fate map, that shows, for a given value of SMAD4 and β-catenin, which cell fates are stable, or, in other words, accessible, in the deterministic model under those signaling levels (Fig. 7A). For example, for low SMAD4 and β-catenin levels, only the pluripotent (P; SOX2+) state is stable and, therefore, cells remain pluripotent under low BMP and WNT levels. On the other hand, for high SMAD4 values and low β-catenin levels, the only stable state is the ExE (E; CDX2+) fate, therefore the model suggests that cells would eventually adopt the ExE fate if induced with high BMP while inhibiting WNT signaling. For high SMAD4 and high β-catenin levels, a bistable region is defined, where both the mesodermal (M; BRA+) and ExE fates are accessible, and cells will become one or the other depending on the initial state of the cell.

**Figure 7.**
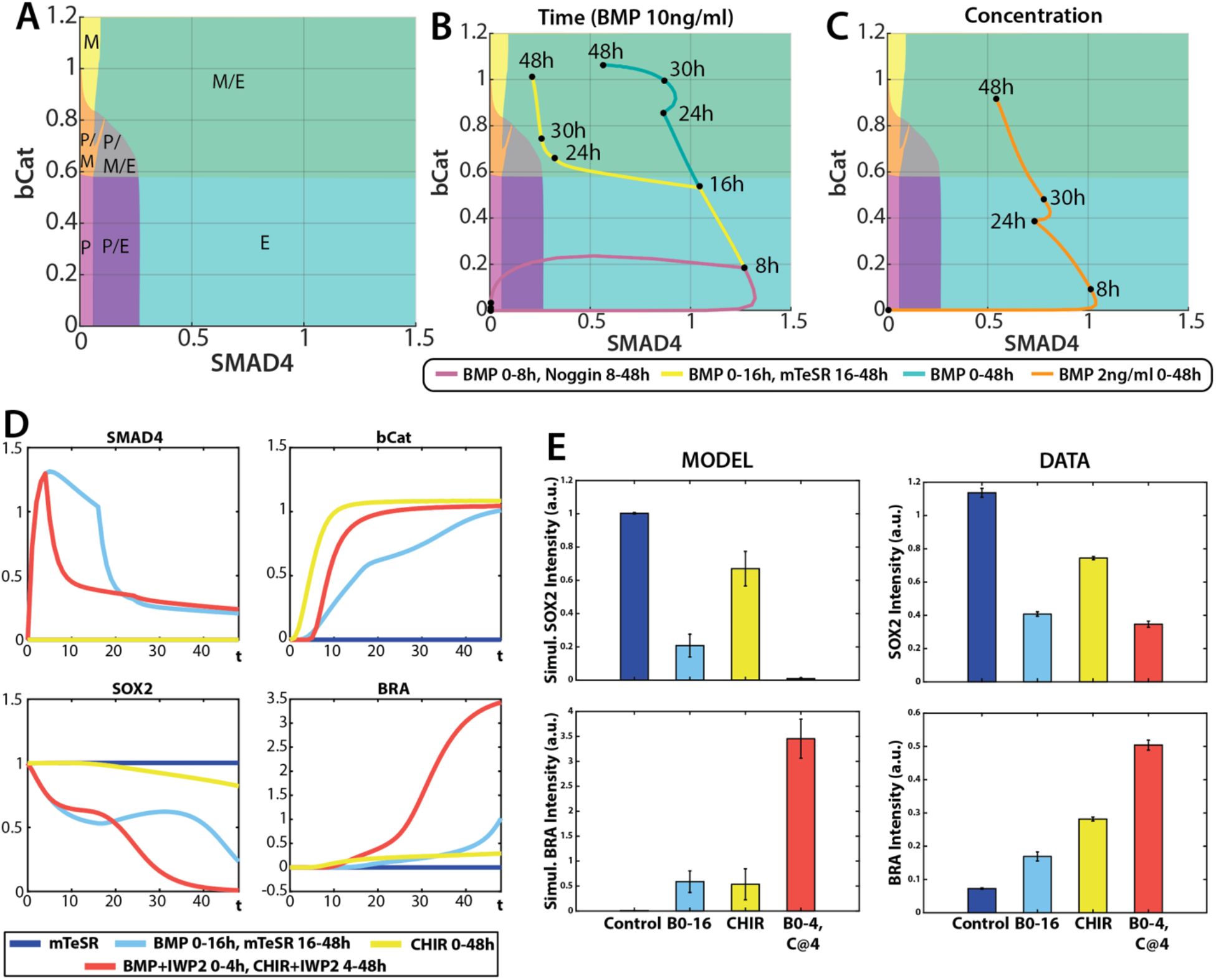
Cell fate is a combinatorial response to BMP and WNT. **(A)** Fate map defined by BMP (SMAD4) and WNT (bCat) levels obtained from the model in Figure 6. P = Pluripotency. M = Mesoderm. E = Extra-embryonic. **(B)** Signaling trajectories on the fate map for the indicated BMP10ng/mL pulses. **(C)** Simulated SMAD4, b-Catenin, SOX2 and BRA dynamics for the conditions indicated on the legend. **(D)** Simulated prediction (right) SOX2 and BRA intensities for the same conditions as in (D), and corresponding experimental confirmation (left).

However, cells are normally not exposed to constant SMAD4 and β-catenin levels, but they experience the signaling dynamics discussed previously, some of which are measured in Fig. 3. Combining these signaling trajectories and the fate map, the model explains the experimental observations by showing that cell fate is a combinatorial response to BMP and WNT. Under short pulses of BMP, endogenous WNT is not activated, so the signaling trajectory moves to a high BMP region for a short period of time after which it goes back to a low BMP, low WNT profile and cells stay pluripotent (Fig. 7B, pink curve, and S11). Under constant high BMP, endogenous WNT is activated, and the signal trajectory ends up in a region of high BMP and high WNT, where both ExE and mesoderm differentiation are possible, however, the trajectory lays in the basin of attraction of the ExE fate and therefore cells adopt this fate (Fig. 7B, blue curve). Under medium durations where WNT becomes self-sustained, even though the signal trajectory initially moves in a high BMP region for some time, decreasing the SOX2 levels in the cell, the withdrawal of the BMP signal situates the signal trajectory in a region of tristability (Fig. 7A-B, gray region), which slows down the decay of pluripotent markers and prevents cells from increasing the expression of ExE markers (Fig. 7B, yellow curve, and S11). This allows cells to become mesoderm once WNT increases and the signaling trajectory moves to the bistable ExE and mesoderm region (Fig. 7B, yellow curve, and S11). On the other hand, constant low concentrations result in a slow WNT upregulation, and therefore the signal trajectory stays in a pluripotency or ExE-promoting region for too long, and are inefficient at promoting mesoderm differentiation (Fig. 7C, S11). Taken together, the model unveils a strategy for efficiently inducing mesoderm with BMP: induction with a high BMP dose that rapidly increases endogenous WNT signaling, after which BMP must be withdrawn for cells to become mesoderm over ExE.

Here we have shown that by pulsing BMP signal, we can obtain a relatively high population of cells that exclusively expresses the mesoderm marker BRA after 2 days. However, this population is heterogeneous and we speculated that this was due to the limitations of the dynamics of endogenous WNT in response to BMP. Inducing hPSCs by CHIR results in a strong, rapid and sustained WNT response (Massey et al., 2019) and yields a very high fraction of mesoderm cells (Fig. S11E). However, this induction is slow, most of the cells still expressing SOX2 after 2 days of induction (Fig. 7E, S11), and taking 3 days to obtain a homogeneous mesodermal population (Fig. S11E). Modeling CHIR through β-catenin activation (Fig. 4) we could reproduce this effect (Fig. 7D,E, yellow condition). We wondered whether we could take advantage of the insights above to speed up mesoderm induction by CHIR. Our goal was to find a 2-day protocol that yielded the highest BRA expression by varying a sequence of inductions with BMP and CHIR. Our model suggested that induction of mesoderm by CHIR was slow because of a strong SOX2 inhibition of the mesoderm state. We reasoned that a pulse of BMP4 signal could be used to destabilize the pluripotency state which, if then followed by CHIR induction, could yield a higher proportion of mesoderm cells by 48 hours. Our model predicted that a 4-hour pulse of BMP was sufficient for this purpose, and if followed by 44 hours of CHIR, would result in a very high BRA expression by day 2 (Fig. 7D,E, red condition). Indeed, experimentally, by day 2 we obtained higher BRA expression and much more complete SOX2 downregulation than with CHIR alone (Fig. 7E, yellow condition). The model explains the combinatorial effect between BMP and Wnt signals in mediating decisions between the pluripotent, mesoderm and ExE fates and offers a platform to rationally design more optimal protocols. Taken together, our study highlights how understanding signaling dynamics can be exploited for developing efficient differentiation protocols.

## Discussion

In this study, we have unveiled a combinatorial mechanism by which BMP and downstream endogenous WNT signaling combinatorially control the cell state transitions observed in mammalian gastrulation and previous *in vitro* studies. We showed a BMP-induced morphogen effect in the duration but not the concentration of signaling, indicating that duration and concentration of BMP signal are not interchangeable in this context. In particular, a specific pulse of high concentration of BMP4 signal is much more efficient at inducing a mesoderm-like state than any constant concentration. Taking advantage of signaling and fate reporter cell lines, we showed that these results depend on an endogenous BMP-induced WNT bistability. A simple minimal cell fate network model explains how the observed cell state transitions are regulated by combinatorial interpretation of these two signals. Induction with a short BMP pulse is not enough to induce endogenous WNT signaling, resulting in hPSCs remaining in the initial pluripotent state. Long pulses of high BMP4 concentration, on the contrary, induce endogenous WNT signals but a high BMP signal overrides the endogenous WNT resulting in extraembryonic differentiation. A pulse of BMP4 of intermediate length that activates WNT autoregulation results in efficient mesoderm differentiation. At lower constant concentrations, WNT activation is slow and cells are therefore pushed towards an ExE fate before experiencing this signal, explaining why low doses of BMP are not interchangeable with a pulse of a high BMP dose. Taken together, our study reveals an underlying logic where, in order to induce efficient mesodermal induction, BMP signaling needs to be sufficiently strong to rapidly induce endogenous WNT signaling upregulation, but sufficiently short for cells to differentiate in a low BMP, high WNT environment.

Our work adds to the growing body of knowledge on how the dynamics of signals are interpreted by downstream regulatory networks. In the case of Shh signaling in the murine neural tube, there is an effective tradeoff between duration and concentration so that shorter durations at higher concentration are sufficient to induce the same fates as longer exposure to lower concentrations (Dessaud et al., 2007). In the case of the three-way decision between pluripotent, primitive streak mesoderm, and extraembryonic amnion, the situation is more complex and relies on the interplay between BMP and Wnt signaling, and cannot be predicted by a parameter of any one of these pathways. Interestingly, we observed that the initial decrease in SOX2 correlated very well with the cumulative integral of BMP signaling as measured by SMAD4 across several different time courses of applied BMP signal (Fig. S9E). This suggests that the initial dissolution of the pluripotent state by BMP signaling may indeed by controlled by the integral of signaling, but that the decision between different potential fates relies on the interpretation of multiple pathways. Indeed, when Wnt signaling is inhibited so that cells can only switch from the pluripotent to the extraembryonic fate, the fraction of cells adopting this fate is well predicted by the integral of the BMP signal (Teague and Heemskerk, submitted).

It is also important to note that measuring the intensity of signaling with a reporter is essential as the external concentration doesn’t translate directly into signaling activity. In the case of BMP signaling, the activation of the SMAD proteins is switch-like and regulated over a narrow range of concentrations, so that it is difficult to regulate the effective activity simply by changing the concentration of ligand in the media (Heemskerk et al., 2019). Further, the loss of the BMP from the media over time causes more prolonged signaling at higher doses (Heemskerk et al., 2019), making it difficult to decouple duration from concentration. Together these features mean that even though the dissolution of pluripotency is responsive to the integral of signaling, in practice, the duration of signal is more easily controlled, and is more likely to be the determining factor in vivo.

It has recently been proposed that BMP4 differentiation rapidly becomes irreversible due to positive feedback through GATA3 (Gunne-braden et al., 2020). In that study, a one-hour pulse of BMP4 was sufficient to induce irreversible differentiation in hPSC colonies, which is in clear contradiction with our results (Fig. 1,2, S1, S2). Irreversible differentiation following short exposure to BMP is contradictory to several studies from our lab and others (Chhabra et al., 2019; Etoc et al., 2016; Jo et al., 2022; Nemashkalo et al., 2017) that showed that BMP signaling is rapidly downregulated by small molecule or extracellular protein inhibitors and that pulses of longer than 10 or 24 hours are needed to induce differentiation in both in micropatterned colonies and regular culture, respectively. Apparently, irreversible differentiation is likely due to incomplete removal of the BMP in the media (Fig. S2C-D), and indeed we found using a live cell reporter for BMP signaling that signaling activity remained substantially above baseline when media containing high doses of BMP was replaced by media alone, however, treatment with Noggin quickly abrogated this continued signaling. Further experiments with washing confirmed that BMP ligands cannot be removed from the media by washing alone (see Fig. S2C-D).

Our results showed that under BMP induction of hPSCs there is a relatively narrow window of durations of BMP exposure which are able to specify mesoderm, and this was related to the dynamics of the induced endogenous WNT signaling. This highlights the need to consider dynamics when developing in vitro differentiation protocols. In particular, while here we investigated the importance of the duration of a single pulse of BMP, more complex time courses of BMP stimulation are possible, and it would be interesting to determine the optimal time courses for achieving different fates. In vivo, while the same cascade of signals controls gastrulation in other mammals, there is a wide range of developmental time scales, and it is unclear by what mechanism the window of BMP duration needed might be matched to the developmental time scale. Interestingly, recent studies have uncovered cell movement as a mechanism that controls the temporal exposure to morphogens (Fulton et al., 2022). Further work could elucidate whether the differences might be due to differential protein stability and cell cycle duration as has been recently proposed in other contexts (Matsuda et al., 2020; Rayon et al., 2020).

We show that there is a form of memory in the WNT signaling so that transient exposure to BMP can be sufficient to induce WNT in a sustained fashion. This is reminiscent of the interplay between WNT and NODAL signaling in which cells remember prior exposure to WNT, which alters their subsequent response to NODAL (Yoney et al., 2018; Yoney et al., 2022). The mechanisms are different as in the case described here BMP leads to ongoing WNT signaling, likely through sustained expression of WNT ligands, while WNT affects the interpretation of NODAL by inducing EOMES without a subsequent requirement for the continuation of WNT activity.

A popular recent approach based on the Waddington landscape metaphor has been shown to be a powerful yet simple way to study cell fate transitions in different contexts (Camacho-Aguilar et al., 2021; Coomer et al., 2022; Corson and Siggia, 2012; Corson and Siggia, 2017; Huang, 2012; Mojtahedi et al., 2016; Sáez et al., 2022; Valcourt et al., 2021). In these models, the differentiation of a cell is depicted as a ball rolling down a landscape of hills and valleys, that represent the different cell types the stem cell can differentiate into. In particular, we and others have shown that using dynamical systems theory one can enumerate the nature of the possible bifurcations, and parametrize these through the signals to build the landscape model, which then can be quantitatively fitted to the data (Camacho-Aguilar et al., 2021; Sáez et al., 2022). These studies, however, lacked signaling dynamics data and the underlying bifurcations had to be guessed and then validated through fitting. Also, these models have not been compared yet to any more mechanistic models. Here using a cell fate network model which, although simple, contains some mechanistic knowledge, we have deciphered the underlying fate map without any prior knowledge about the candidate bifurcations. Interestingly, our three mutually repulsive state network contains the bifurcations present in the elliptic-umbilic catastrophe, as had been proposed before (Rand et al., 2021). To our knowledge, this is the first experimental system that confirms this idea. We believe that our results provide a foundation to quantitatively compare mechanistic and landscape models using quantitative data on signaling dynamics.

Much work has been done to decipher the dynamics of signals that control the patterning of micropatterned hPSC colonies treated with BMP4 (Chhabra et al., 2019; Etoc et al., 2016; Heemskerk et al., 2019; Warmflash et al., 2014). These studies have unveiled that, initially, there is a homogeneous response to BMP4 across the whole colony, which is restricted to the edge via receptor localization and accumulation of Noggin at the colony center between 10 and 20 hours after treatment (Etoc et al., 2016; Heemskerk et al., 2019). Subsequently, around 30h, waves of WNT and Nodal signaling start near the edge of the colony and move inwards, spatially correlating with a ring of BRA-positive mesodermal cells (Chhabra et al., 2019; Heemskerk et al., 2019). Although our results have been obtained in a culture with lower cell density where self-organized patterning does not occur, they are consistent with the observations of a pulse of BMP throughout the colony which induces endogenous WNT signal. The cells that adopt a mesodermal fate are those that are displaced from the edge and, therefore, only experience transient BMP signaling followed by upregulation of Wnt signaling to high levels. Thus, the model developed here could be a good starting point for building a spatial model to understand how patterns arise from the interplay of dynamic signaling and combinatorial interpretation.

## Materials and Methods

### Cell lines

The cell lines used were ESI017 (NIHhPSC-11-0093), ESI017 GFP::β-Catenin RFP::H2B (Massey et al., 2019), RUES2 GFP::Smad4 RFP::H2B (Nemashkalo et al., 2017), and ESI017 SOX2::mCitrine CFP::H2B (this study). ESI017 cells were obtained directly from ESIBIO while RUES2 were a gift of Ali Brivanlou (Rockefeller University). The identity of these cells as pluripotent cells was confirmed via triple staining for pluripotency markers OCT4, SOX2, and NANOG. All cells were routinely tested for mycoplasma contamination and found negative.

### Cell Culture, Treatments, and Differentiation

All cell lines were maintained in pluripotency maintenance culture as described in (Nemashkalo et al., 2017). ESI017 SOX2::mCitrine CFP::H2B were maintained in mTeSR Plus medium (STEMCELL Technologies; 100-0276) and Blasticidin (5μg/ml; A.G. Scientific; B-1247-SOL) for selection, which was removed before experiments.

Ibidi μ-Slide 8 Well plates (Ibidi; 80826) were used for experiments, which were coated with Matrigel (5μl/ml; Corning; 354277) diluted in DMEM/F12 (VWR;45000-344). For all experiments, cells were seeded into mTeSR1 medium (STEMCELL Technologies; 85857) containing rock inhibitor Y27672 (10μM; STEMCELL Technologies; 05875) at a density of 4 x 10^4^/cm^2^ (except when noted otherwise). Treatment started 21 hours after seeding, and media was always changed every 24 hours, and when performing the indicated specific treatments.

The following recombinant proteins and small molecules were used: BMP4 (R&D Systems; 314BP050), Noggin (500ng/ml; R&D Systems; 6057-NG-100), IWP2 (Stemgent; 040034; 3μM), CHIR99021 (24μM; MedChem Express; HY-10182).

For micropatterning experiments, these were done in a 96-well plate (CYTOO) of 700 μm circular micropatterns. Coating and seeding were done as previously described by (Warmflash et al., 2014). Briefly, wells were coated with Human Recombinant Laminin 511 (Fisher Scientific) in a 1:20 dilution in PBS with calcium and magnesium for 3 hours at 37 °C. hESCs were then seeded as single cells and maintained in mTeSR medium. After overnight incubation, cells were treated with 50 ng/ml BMP4 (R&D systems) for 48 hours and with 1 hour pulse of 50 ng/ml BMP4 (R&D systems), as previously described in (Gunne-braden et al., 2020), with or without 100 ng/ml of Noggin (Fisher Scientific).

The following recombinant proteins and small molecules were used: BMP4 (R&D Systems; 314BP050), Noggin (250ng/ml; R&D Systems; 6057-NG-100), IWP2 (Stemgent; 040034; 3μM), CHIR99021 (24μM; MedChem Express; HY-10182).

### Plasmids and Generation of SOX2-mCitrine-Labeled hPSC Cell Line

We used CRISPR-Cas9 technology for gene editing, where the ESI017 hPSC line was used as the parental line. We used previously published constructs to fuse mCitrine directly with SOX2 at the C-terminus of the SOX2 coding sequence (Martyn et al., 2018). The SOX2 homology donor consists of a 1-kb homology arm, an mCitrine::T2A::blasticidin cassette, and a 1-kb right homology arm. Cas9 expression plasmid, homology donor DNA (Plasmid AW-P46), and guide RNA (Plasmid AW-P45; GTGCCCGGCACGGCCATTAA) were nucleofected in hPSCs using the P3 Primary Cell 4D-Nucleofector X Kit (Lonza; V4Xp-3012), and positive transformants were selected with blasticidin (10μg/ml; A.G. Scientific; B-1247-SOL) and CloneR (STEMCELL Technologies; 05889) for two days, after which cells were passaged and single clones were handpicked and amplified. Sanger sequencing was performed to screen and confirm a successful clone (Primer sequences are listed in Table 1). After establishment, the stable line was checked for pluripotency markers, i.e. OCT4, SOX2 and NANOG expression, as well as BMP differentiation both in regular culture and micropatterning, and was found indistinguishable from WT ESI017 hPSCs.

**Table 1.**
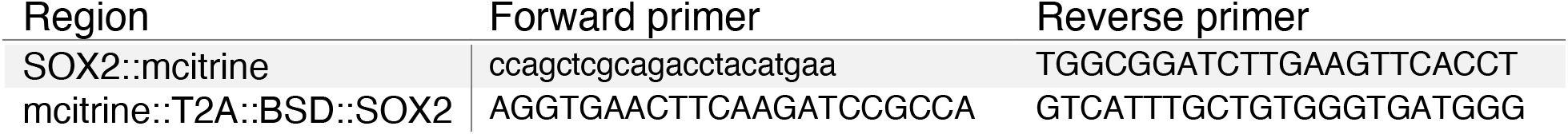

An ePiggyBac (ePB) master vector based on the pBSSK backbone (Lacoste et al., 2009), harboring transposon-specific inverted terminal repeat sequences (ITR) was modified to deliver a nuclear marker CFP::H2B (Plasmid AW-P68). ePB master vector and helper (Plasmid AW-27) were nucleofected into the established SOX2::mCitrine cell line using the P3 Primary Cell 4D-Nucleofector X Kit (Lonza; V4Xp-3012). G-418 (40ng/mL; ThermoFisher; 10131035) started two days after nucleofection and lasted for at least 7 days.

### Immunofluorescence Antibody Staining

Cells were fixed with 4% PFA and stained as described in (Nemashkalo et al., 2017). Antibodies and dilutions used are listed in Table 2.

**Table 2.** Imaging and Analysis.

| Protein | Species | Dilution | Vendor | Catalog no. |
| --- | --- | --- | --- | --- |
| Sox2 | Rabbit | 1:200 | Cell Signalling Technologies | 23064s |
| Brachyury | Goat | 1:300 | R&D Systems | AF2085 |
| Brachyury | Rabbit | 1:400 | R&D Systems | MAB20851 |
| Cdx2 | Mouse | 1:100 | Biogenex | MU392A-5UC |
| Isl1 | Mouse | 1:50 | DSHB | 39.4D-5 |
| Oct4 | Mouse | 1:400 | BD Biosciences | 611203 |
| Nanog | Mouse | 1:400 | BD Biosciences | 560482 |
| Nanog | Goat | 1:200 | R&D Systems | AF1997 |
| Hand1 | Goat | 1:200 | R&D Systems | AF3168 |
| Gata3 | Rabbit | 1:100 | ThermoFisher | PA1-101 |
| Tbx6 | Goat | 1:200 | R&D Systems | AF4744 |

Imaging acquisition was done on an Olympus/Andor spinning disk confocal microscope with a 20x, 0.75NA air objective.

For live imaging experiments, reporter cell lines were maintained in antibiotic selection for the H2B fluorescent marker for at least three days and stopping 2 days before seeding, at the latest, to maximize the number of fluorescent cells in the culture. Time-lapse imaging intervals were 15 minutes, and Z-stacks were acquired in three planes spaced 2.5μM apart. During imaging, temperature (37 °C), humidity (~50%), and CO2 (5%) were controlled, and media change was performed without moving the plate from the stage. 8 positions of each condition were selected for imaging.

Image analysis was performed using Ilastik (Berg et al., 2019; Sommer et al., 2011) for initial segmentation and custom-written MATLAB code available at https://github.com/warmflashlab/Camacho-Aguilar2022 BMPWNT for further analysis. Smad4 dynamics was quantified as the nuclear to cytoplasmic Smad4 ratio, (Heemskerk et al., 2019). Nuclear protein expression was measured by mean nuclear intensity and normalized by mean nuclear DAPI to correct for intensity variations due to optics.

For the analysis of the micropatterning experiment, stitching of the colonies was performed in Fiji using the algorithm in (Preibisch et al., 2009). Segmentation and mean intensity quantification were done on Ilastik (Berg et al., 2019; Sommer et al., 2011) and custom software written in MATLAB (MathWorks), previously described in (Warmflash et al., 2014).

## Cell Fate Classification

(see Appendix)

## Mathematical Models

(see Appendix)

## Acknowledgements

We thank Idse Heemskerk, Seth Teague, Francisco-Jesús Castro-Jiménez, Nathan Lord, and Eric Siggia for helpful discussions on the project. We thank the Brivanlou lab for sharing the plasmids used to create the SOX2::mCitrine cell line. Sapna Chhabra for performing the experiment in Fig S8C and for helpful feedback on the project. Joseph Massey for sharing helpful pieces of code for analyzing live-cell imaging data. Cecilia Guerra and Lizhong Liu for technical assistance with experiments; and all the members of the Warmflash lab for helpful feedback. The work was supported by grants to AW from the National Science Foundation (MCB-1553228 and MCB-2135296) and the Simons Foundation (511079).

