## Supplementary figures and text for "Combinatorial interpretation of BMP and WNT allows BMP to act as a morphogen in time but not in concentration"

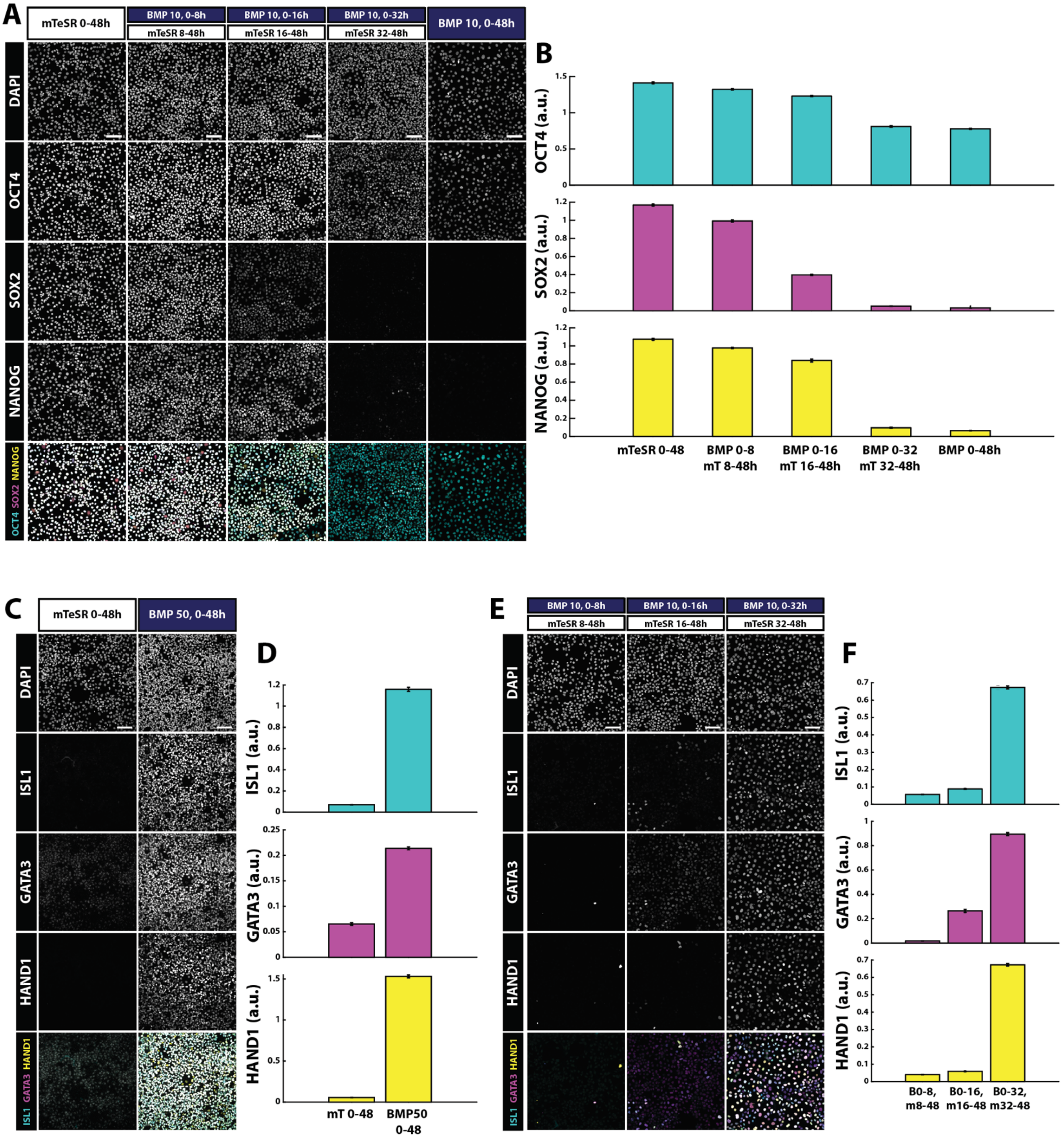

A
mTeSR 0-48h
BMP 10, 0-8h
mTeSR 8-48h
BMP 10, 0-16h
mTeSR 16-48h
BMP 10, 0-32h
mTeSR 32-48h
BMP 10, 0-48h
DAPI
OCT4
SOX2
NANOG
OCT4 SOX2 NANOG
B
OCT4 (a.u.)
SOX2 (a.u.)
NANOG (a.u.)
mTeSR 0-48
BMP 0-8 mT 8-48h
BMP 0-16 mT 16-48h
BMP 0-32 mT 32-48h
BMP 0-48h
C
mTeSR 0-48h
BMP 50, 0-48h
DAPI
ISL1
GATA3
HAND1
ISL1 GATA3 HAND1
D
ISL1 (a.u.)
GATA3 (a.u.)
HAND1 (a.u.)
mT 0-48
BMP50 0-48
E
BMP 10, 0-8h
mTeSR 8-48h
BMP 10, 0-16h
mTeSR 16-48h
BMP 10, 0-32h
mTeSR 32-48h
DAPI
ISL1
GATA3
HAND1
ISL1 GATA3 HAND1
F
ISL1 (a.u.)
GATA3 (a.u.)
HAND1 (a.u.)
B0-8, m8-48
B0-16, m16-48
B0-32, m32-48

**Figure S1. Expression of pluripotency and extraembryonic markers under different BMP treatments. (A)** Example images of immunofluorescence for OCT4, SOX2, and NANOG in cyan, magenta, and yellow, respectively, after the indicated pulses of 10ng/mL BMP4 treatments. **(B)** Quantifications of OCT4, SOX2, and NANOG intensities in (A). Error bars indicate standard error of the mean over n=6 images. **(C)** Example images of immunofluorescence for ISL1, GATA3, and HAND1 in cyan, magenta, and yellow, respectively, after the indicated BMP4 treatments. **(D)** Quantifications of the ISL1, GATA3, and HAND1 intensities in (C). Error bars indicate standard error of the mean over n=6 images. **(E)** Example images of immunofluorescence for ISL1, GATA3, and HAND1 in cyan, magenta, and yellow, respectively, after the indicated pulses of 10ng/mL BMP4 treatments. **(F)** Quantifications of the ISL1, GATA3, and HAND1 intensities in (E). Error bars indicate standard error of the mean over n=6 images. Scale bars: 100um.

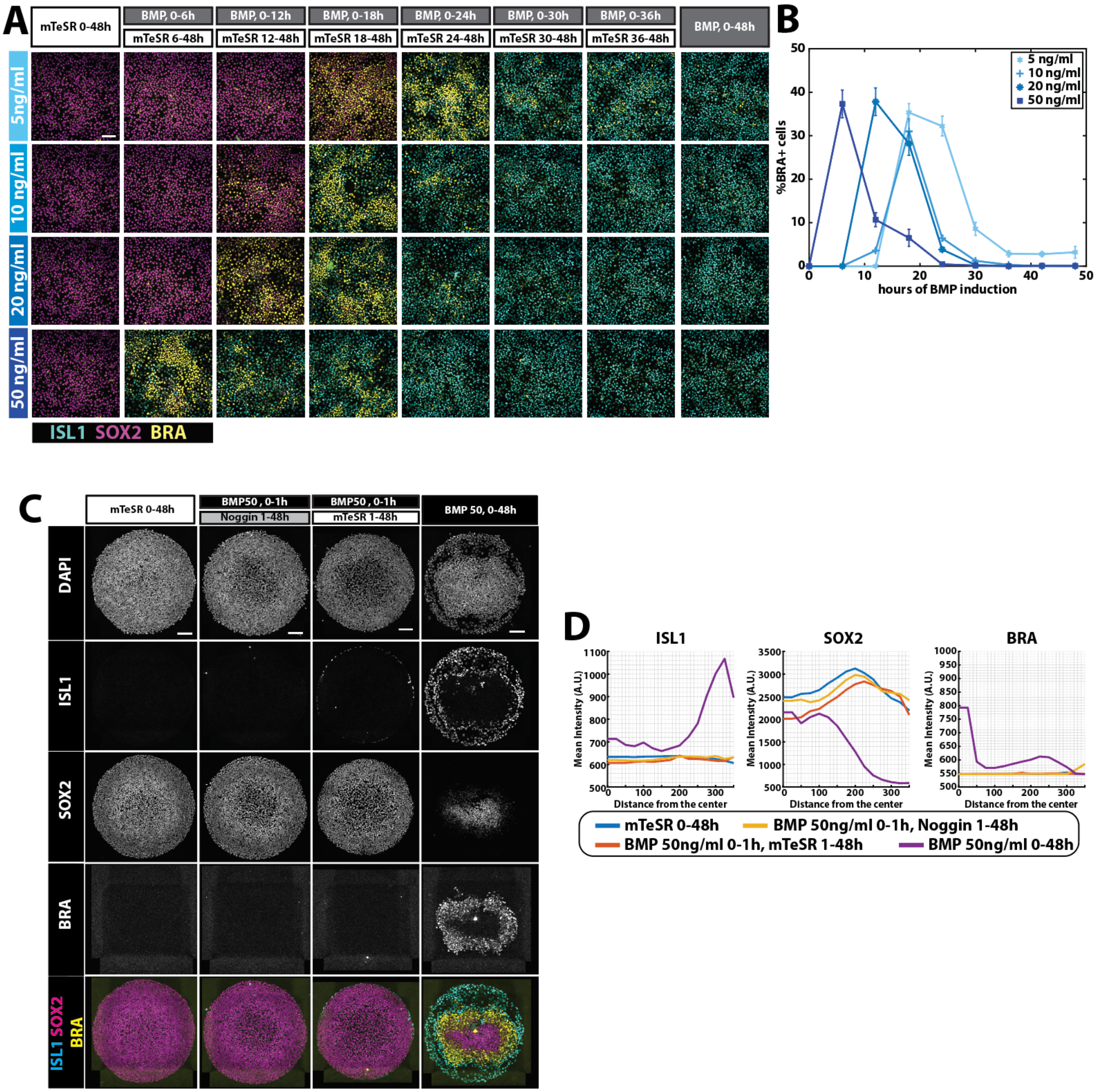
A
mTeSR 0-48h
BMP, 0-6h
mTeSR 6-48h
BMP, 0-12h
mTeSR 12-48h
BMP, 0-18h
mTeSR 18-48h
BMP, 0-24h
mTeSR 24-48h
BMP, 0-30h
mTeSR 30-48h
BMP, 0-36h
mTeSR 36-48h
BMP, 0-48h
5ng/ml
10 ng/ml
20 ng/ml
50 ng/ml
ISL1 SOX2 BRA
B
%BRA+ cells
hours of BMP induction
5 ng/ml
10 ng/ml
20 ng/ml
50 ng/ml
C
mTeSR 0-48h
BMP50 , 0-1h
Noggin 1-48h
BMP50 , 0-1h
mTeSR 1-48h
BMP 50, 0-48h
DAPI
ISL1
SOX2
BRA
ISL1 SOX2 BRA
D
ISL1
SOX2
BRA
Mean Intensity (A.U.)
Distance from the center
mTeSR 0-48h
BMP 50ng/ml 0-1h, Noggin 1-48h
BMP 50ng/ml 0-1h, mTeSR 1-48h
BMP 50ng/ml 0-48h

**Figure S2. Duration vs concentration of BMP treatment under mTeSR withdrawal. (A)** Example images of immunofluorescence for ISL1, SOX2, and BRA in cyan, magenta, and yellow, respectively, after the indicated pulses (columns) of the indicated concentrations (rows) of BMP4. **(B)** Quantifications of the proportions of cells classified as BRA+ in the treatments shown in (A). Error bars indicate SEM over n=6 images. **(C)** Example images of immunofluorescence for ISL1, SOX2, and BRA in cyan, magenta, and yellow, respectively, of micropatterned hPSCs after the indicated pulses of 50ng/mL BMP4. **(D)** Quantification of experiment shown in C (n=5 colonies per condition). Scale bars: 100um.

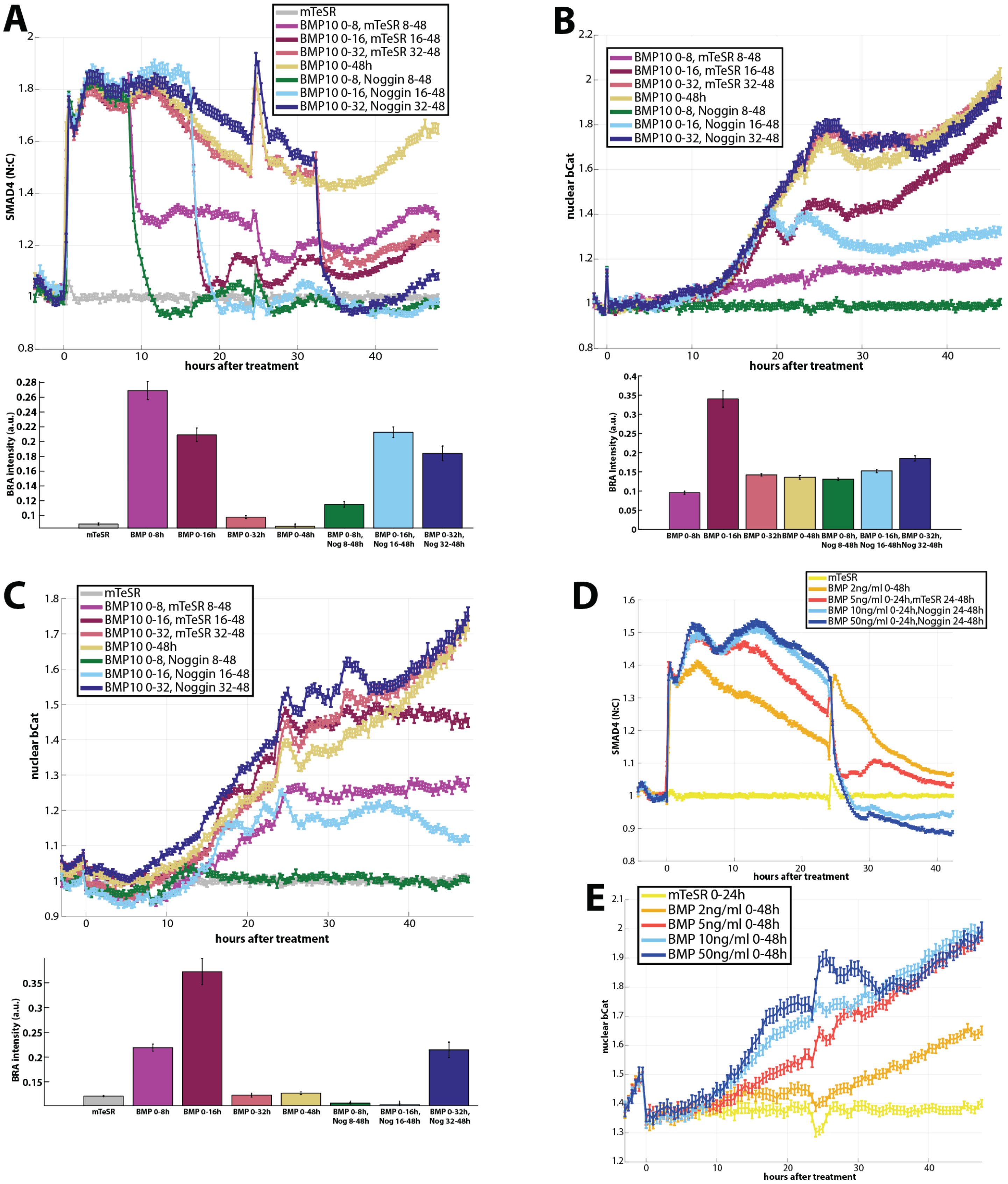
A
mTeSR
BMP10 0-8, mTeSR 8-48
BMP10 0-16, mTeSR 16-48
BMP10 0-32, mTeSR 32-48
BMP10 0-48h
BMP10 0-8, Noggin 8-48
BMP10 0-16, Noggin 16-48
BMP10 0-32, Noggin 32-48
SMAD4 (N:C)
hours after treatment
BRA intensity (a.u.)
mTeSR
BMP 0-8h
BMP 0-16h
BMP 0-32h
BMP 0-48h
BMP 0-8h, Nog 8-48h
BMP 0-16h, Nog 16-48h
BMP 0-32h, Nog 32-48h
B
BMP10 0-8, mTeSR 8-48
BMP10 0-16, mTeSR 16-48
BMP10 0-32, mTeSR 32-48
BMP10 0-48h
BMP10 0-8, Noggin 8-48
BMP10 0-16, Noggin 16-48
BMP10 0-32, Noggin 32-48
nuclear bCat
hours after treatment
BRA Intensity (a.u.)
BMP 0-8h
BMP 0-16h
BMP 0-32h
BMP 0-48h
BMP 0-8h, Nog 8-48h
BMP 0-16h, Nog 16-48h
BMP 0-32h, Nog 32-48h
C
mTeSR
BMP10 0-8, mTeSR 8-48
BMP10 0-16, mTeSR 16-48
BMP10 0-32, mTeSR 32-48
BMP10 0-48h
BMP10 0-8, Noggin 8-48
BMP10 0-16, Noggin 16-48
BMP10 0-32, Noggin 32-48
nuclear bCat
hours after treatment
BRA intensity (a.u.)
mTeSR
BMP 0-8h
BMP 0-16h
BMP 0-32h
BMP 0-48h
BMP 0-8h, Nog 8-48h
BMP 0-16h, Nog 16-48h
BMP 0-32h, Nog 32-48h
D
mTeSR
BMP 2ng/ml 0-48h
BMP 5ng/ml 0-24h,mTeSR 24-48h
BMP 10ng/ml 0-24h,Noggin 24-48h
BMP 50ng/ml 0-24h,Noggin 24-48h
SMAD4 (N:C)
hours after treatment
E
mTeSR 0-24h
BMP 2ng/ml 0-48h
BMP 5ng/ml 0-48h
BMP 10ng/ml 0-48h
BMP 50ng/ml 0-48h
nuclear bCat
hours after treatment

**Figure S3. BMP and WNT dynamics under different BMP4 durations and concentrations. (A)** (Top) GFP::SMAD4 average nuclear:cytoplasmic intensity ratio after the indicated pulses of 10ng/ml BMP4. This figure shows additional experimental conditions from the same data set shown in Fig. 3D. Intensities are normalized by the mTeSR control condition. (Bottom) Quantification of BRA intensity at 48h after each treatment. Error bars indicate SEM over n=8 images. **(B)** (Top) GFP::β-catenin average nuclear intensity after the indicated treatments. Intensities are normalized by the BMP 0-8h,Noggin 8-48h due to lack of mTeSR control, and that this condition was very similar to mTeSR condition in the repeat of the experiment in (C). (Bottom) Quantification of BRA intensity at 48h after each treatment. Error bars indicate SEM over n=8 images. **(C)** Repeat of the experiment shown in (B). (Top) GFP::β-catenin average nuclear intensity after the indicated treatments. This figure shows additional experimental conditions from the same data set shown in Fig. 3H. Intensities are normalized by the mTeSR control condition. (Bottom) Quantification of BRA intensity at 48h after each treatment. Error bars indicate SEM over n=8 images. **(D)** GFP::SMAD4 average nuclear:cytoplasmic intensity ratio after the indicated treatments. **(E)** GFP::β-catenin average nuclear intensity after the indicated treatments.

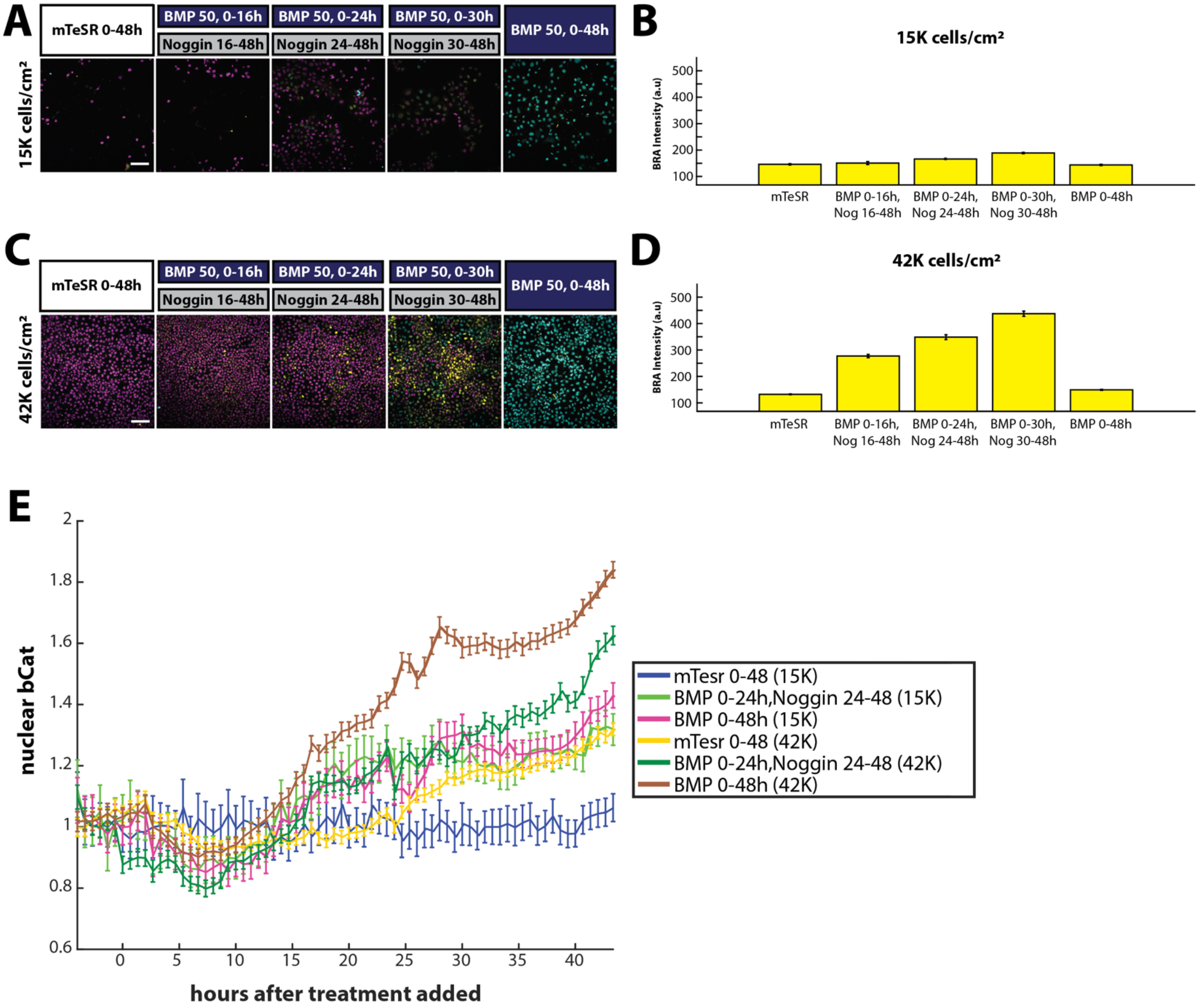
A
mTeSR 0-48h
BMP 50, 0-16h
Noggin 16-48h
BMP 50, 0-24h
Noggin 24-48h
BMP 50, 0-30h
Noggin 30-48h
BMP 50, 0-48h
15K cells/cm²
B
15K cells/cm²
BRA Intensity (a.u)
mTeSR
BMP 0-16h, Nog 16-48h
BMP 0-24h, Nog 24-48h
BMP 0-30h, Nog 30-48h
BMP 0-48h
C
mTeSR 0-48h
BMP 50, 0-16h
Noggin 16-48h
BMP 50, 0-24h
Noggin 24-48h
BMP 50, 0-30h
Noggin 30-48h
BMP 50, 0-48h
42K cells/cm²
D
42K cells/cm²
BRA Intensity (a.u)
mTeSR
BMP 0-16h, Nog 16-48h
BMP 0-24h, Nog 24-48h
BMP 0-30h, Nog 30-48h
BMP 0-48h
E
nuclear bCat
hours after treatment added
mTesr 0-48 (15K)
BMP 0-24h,Noggin 24-48 (15K)
BMP 0-48h (15K)
mTesr 0-48 (42K)
BMP 0-24h,Noggin 24-48 (42K)
BMP 0-48h (42K)

**Figure S4. Mesoderm induction by BMP4 requires a minimum cell density. (A)** Example images of immunofluorescence for ISL1, SOX2, and BRA in cyan, magenta, and yellow, respectively, after the indicated pulses of BMP4 followed by Noggin. Seeding density was 15K cells/$cm^2$ (less than 40% of the usual seeding density). This experiment was done in 18-well ibidi plates. **(B)** Quantification of BRA intensity at 48h after each treatment. Error bars indicate SEM over n=9 images. **(C)** Example images of immunofluorescence for ISL1, SOX2, and BRA in cyan, magenta, and yellow, respectively, after the indicated pulses of BMP4 followed by Noggin. Seeding density was 42K cells/$cm^2$ (similar to the usual seeding density). This experiment was done in 18-well ibidi plates. **(D)** Quantification of BRA intensity at 48h after each treatment. Error bars indicate SEM over n=9 images. **(E)** GFP::β-catenin average nuclear intensity after the indicated treatments under the indicated seeding densities (in brackets). Intensities are normalized by the mTeSR control condition under a seeding density of 15K cells/$cm^2$.

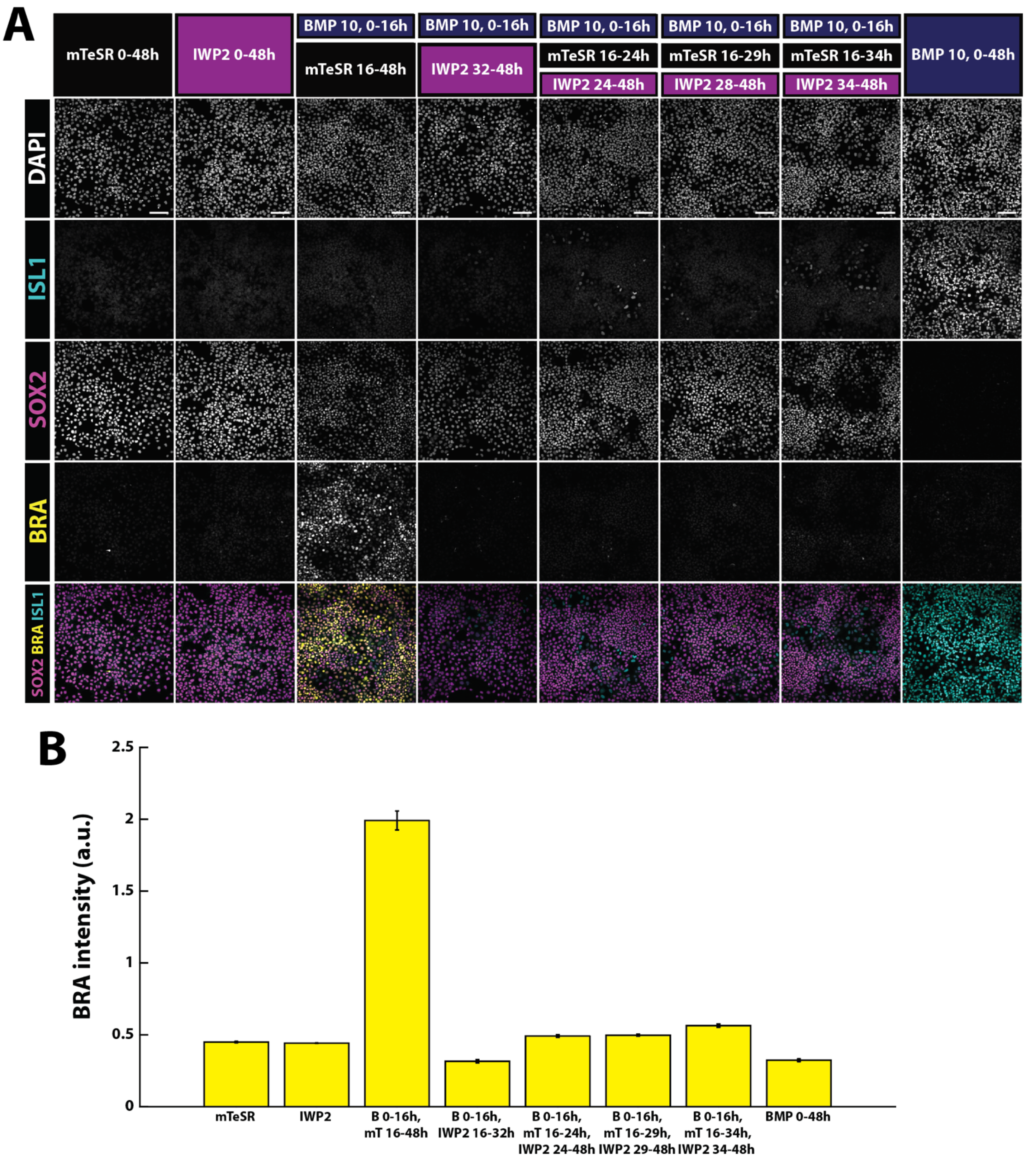
A
mTeSR 0-48h
IWP2 0-48h
BMP 10, 0-16h
mTeSR 16-48h
BMP 10, 0-16h
IWP2 32-48h
BMP 10, 0-16h
mTeSR 16-24h
IWP2 24-48h
BMP 10, 0-16h
mTeSR 16-29h
IWP2 28-48h
BMP 10, 0-16h
mTeSR 16-34h
IWP2 34-48h
BMP 10, 0-48h
DAPI
ISL1
SOX2
BRA
SOX2 BRA ISL1
B
BRA intensity (a.u.)
2.5
2
1.5
1
0.5
0
mTeSR
IWP2
B 0-16h, mT 16-48h
B 0-16h, IWP2 16-32h
B 0-16h, mT 16-24h, IWP2 24-48h
B 0-16h, mT 16-29h, IWP2 29-48h
B 0-16h, mT 16-34h, IWP2 34-48h
BMP 0-48h

**Figure S5. WNT signal is necessary for BRA induction. (A)** Example images of immunofluorescence for ISL1, SOX2, and BRA in cyan, red, and yellow, respectively, after the indicated treatments. **(B)** Quantification of BRA intensity at 48h after each treatment. Error bars indicate SEM over n=8 images.

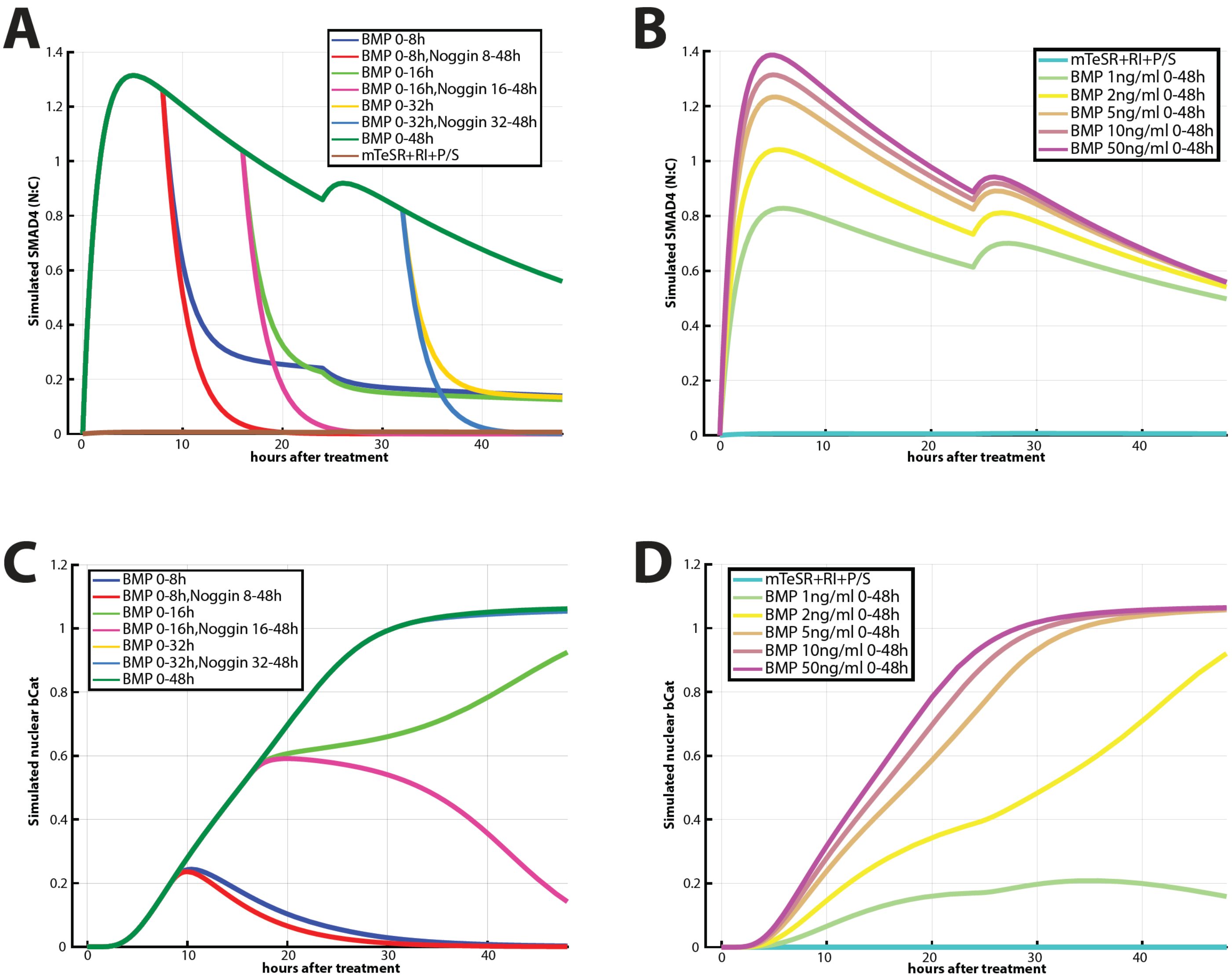
A
B
C
D
Simulated SMAD4 (N:C)
hours after treatment
BMP 0-8h
BMP 0-8h,Noggin 8-48h
BMP 0-16h
BMP 0-16h,Noggin 16-48h
BMP 0-32h
BMP 0-32h,Noggin 32-48h
BMP 0-48h
mTeSR+RI+P/S
BMP 1ng/ml 0-48h
BMP 2ng/ml 0-48h
BMP 5ng/ml 0-48h
BMP 10ng/ml 0-48h
BMP 50ng/ml 0-48h
Simulated nuclear bCat

**Figure S6. Simulated BMP and WNT dynamics under different BMP4 durations and concentrations. (A)** Simulated SMAD4 intensity after the indicated pulses of 10ng/ml BMP4. **(B)** Simulated SMAD4 intensity after the indicated concentrations of BMP4. **(C)** Simulated nuclear β-catenin intensity after the indicated pulses of 10ng/ml BMP4. **(D)** Simulated nuclear β-catenin intensity after the indicated concentrations of BMP4.

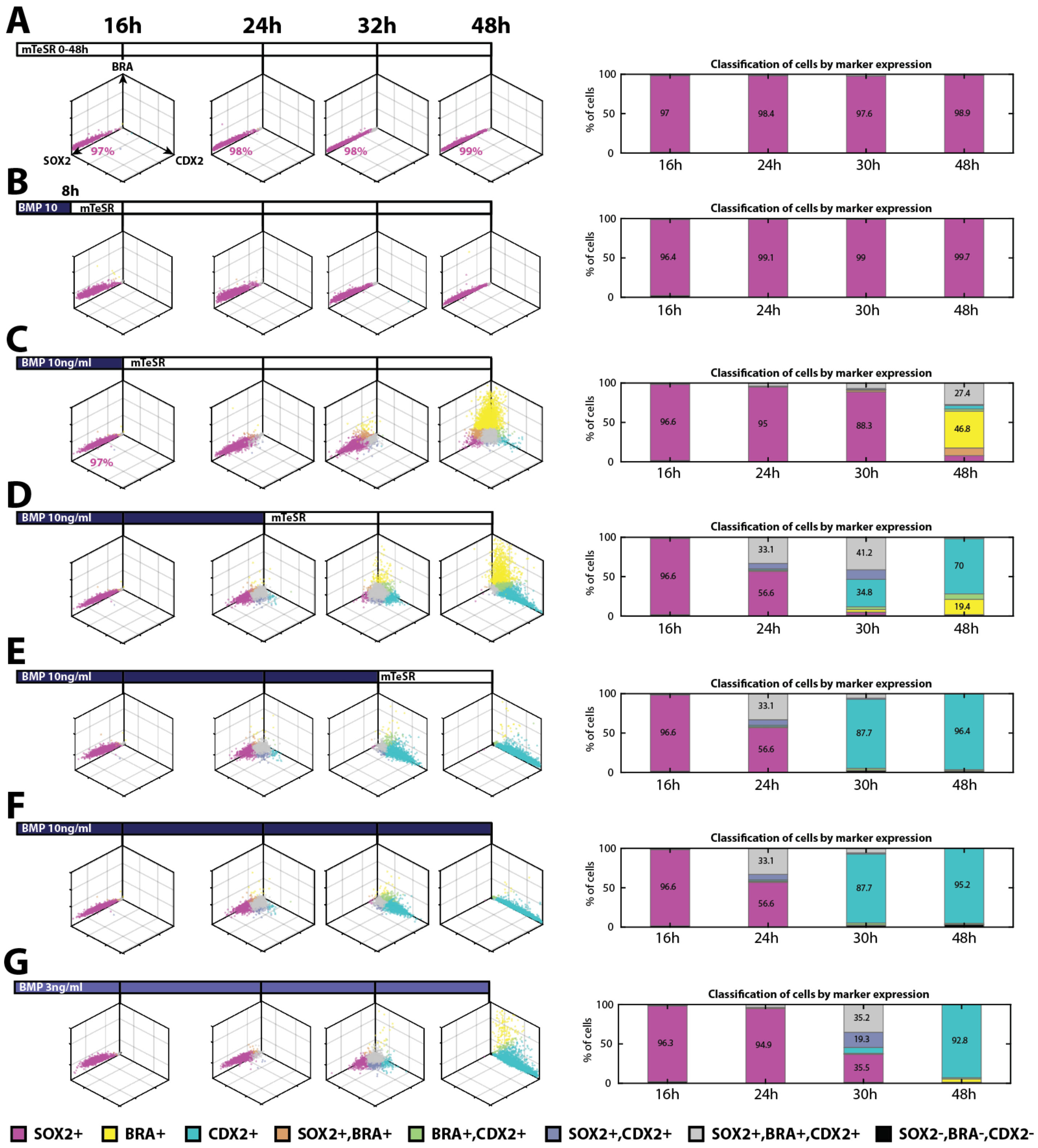

A
16h
24h
32h
48h
mTeSR 0-48h
BRA
SOX2
CDX2
97%
98%
98%
99%
Classification of cells by marker expression
% of cells
97
98.4
97.6
98.9
16h
24h
30h
48h
B
8h
BMP 10
mTeSR
96.4
99.1
99
99.7
C
BMP 10ng/ml
mTeSR
97%
96.6
95
88.3
27.4
46.8
D
BMP 10ng/ml
mTeSR
96.6
33.1
56.6
41.2
34.8
70
19.4
E
BMP 10ng/ml
mTeSR
96.6
33.1
56.6
87.7
96.4
F
BMP 10ng/ml
96.6
33.1
56.6
87.7
95.2
G
BMP 3ng/ml
96.3
94.9
35.2
19.3
35.5
92.8
SOX2+
BRA+
CDX2+
SOX2+,BRA+
BRA+,CDX2+
SOX2+,CDX2+
SOX2+,BRA+,CDX2+
SOX2-,BRA-,CDX2-

**Figure S7. Markers dynamics under different BMP4 treatments.** This figure shows additional experimental conditions from the same data set shown in Fig. 5. Scatter plots of the quantifications of SOX2, BRA and CDX2 (left) and quantifications of cell fate proportions (right) under the treatments mTeSR 0-48h (A), BMP 10ng/mL 0-8h followed by mTeSR 8-48h (B), BMP 10ng/mL 0-16h followed by mTeSR 16-48h (C), BMP 10ng/mL 0-24h followed by mTeSR 24-48h (D), BMP 10ng/mL 0-32h followed by mTeSR 32-48h (E), BMP 10ng/mL 0-48h (F), BMP 3ng/mL 0-48h at 16h, 24h, 32h, and 48h after treatment. Each dot corresponds to a single cell, and its color marks the cell fate assigned to the cell, as shown in the legend at the bottom.

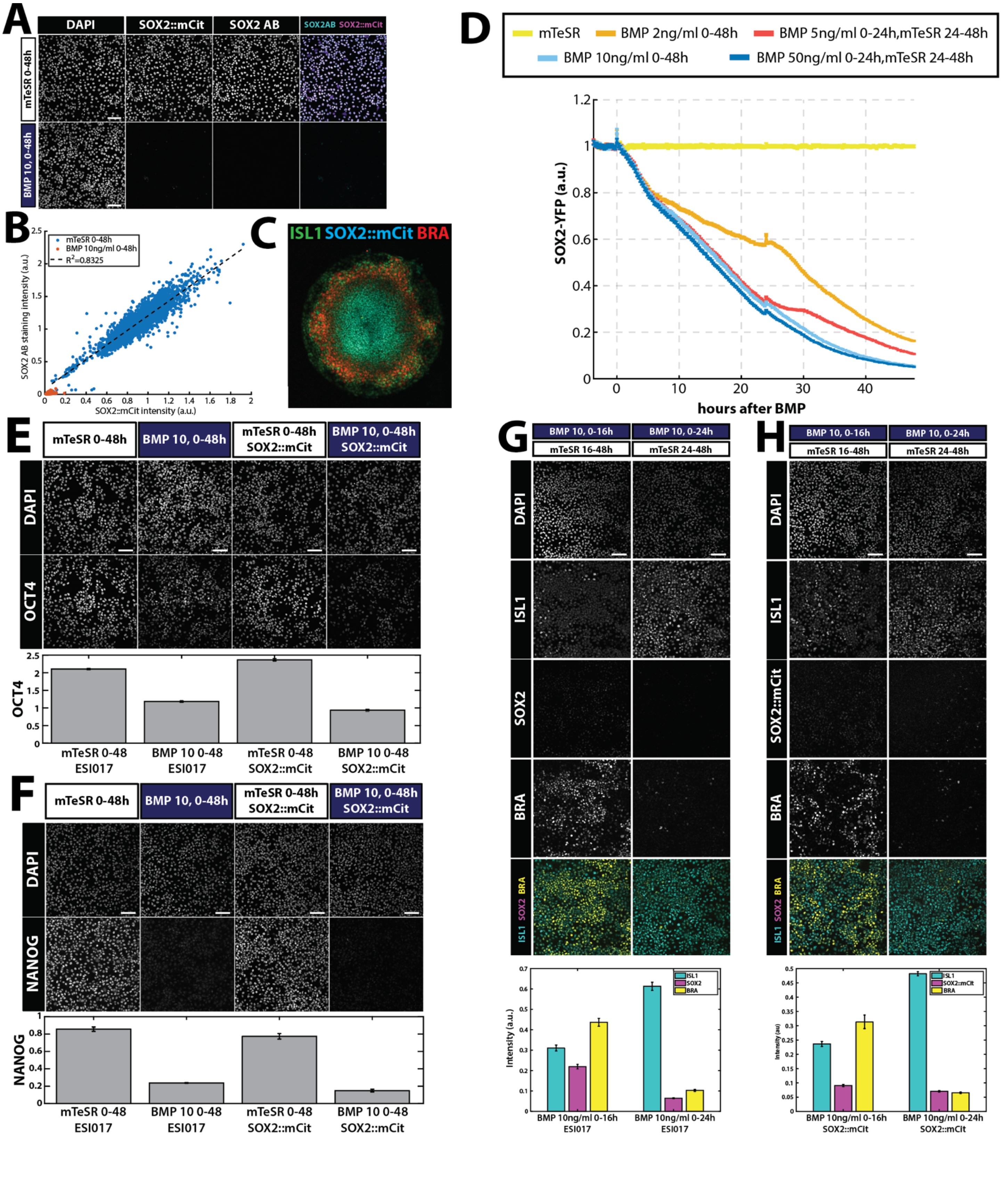

A
DAPI
SOX2::mCit
SOX2 AB
SOX2AB SOX2::mCit
mTeSR 0-48h
BMP 10, 0-48h
B
mTeSR 0-48h
BMP 10ng/ml 0-48h
R²=0.8325
SOX2 AB staining intensity (a.u.)
SOX2::mCit intensity (a.u.)
C
ISL1 SOX2::mCit BRA
D
mTeSR
BMP 2ng/ml 0-48h
BMP 5ng/ml 0-24h,mTeSR 24-48h
BMP 10ng/ml 0-48h
BMP 50ng/ml 0-24h,mTeSR 24-48h
SOX2-YFP (a.u.)
hours after BMP
E
mTeSR 0-48h
BMP 10, 0-48h
mTeSR 0-48h SOX2::mCit
BMP 10, 0-48h SOX2::mCit
DAPI
OCT4
mTeSR 0-48 ESI017
BMP 10 0-48 ESI017
mTeSR 0-48 SOX2::mCit
BMP 10 0-48 SOX2::mCit
F
mTeSR 0-48h
BMP 10, 0-48h
mTeSR 0-48h SOX2::mCit
BMP 10, 0-48h SOX2::mCit
DAPI
NANOG
mTeSR 0-48 ESI017
BMP 10 0-48 ESI017
mTeSR 0-48 SOX2::mCit
BMP 10 0-48 SOX2::mCit
G
BMP 10, 0-16h
mTeSR 16-48h
BMP 10, 0-24h
mTeSR 24-48h
DAPI
ISL1
SOX2
BRA
ISL1 SOX2 BRA
Intensity (a.u.)
BMP 10ng/ml 0-16h ESI017
BMP 10ng/ml 0-24h ESI017
H
BMP 10, 0-16h
mTeSR 16-48h
BMP 10, 0-24h
mTeSR 24-48h
DAPI
ISL1
SOX2::mCit
BRA
ISL1 SOX2 BRA
Intensity (au)
SOX2::mCit
BMP 10ng/ml 0-16h SOX2::mCit
BMP 10ng/ml 0-24h SOX2::mCit

**Figure S8. SOX2::mCitrine fusion cell line accurately marks SOX2 expression. (A)** Example images of immunofluorescence for SOX2 (SOX2 AB) of the S SOX2::mCitrine fusion cell line after the indicated treatments. **(B)** Scatter plots of the quantifications of SOX2 antibody staining vs SOX2::mCitrine in (A). Each dot corresponds to a single cell. $R^2$=0.8325. **(C)** Example image of immunofluorescence for ISL1 and BRA of micropatterned SOX2::mCitrine cells treated with 50ng/ml BMP4 for 48h. **(D)** SOX2::mCitrine average nuclear intensity under the indicated treatments. **(E-H)** Comparison of pluripotency and cell fate markers expression between hPSCs (E,G) and SOX2::mCitrine tagged hPSCs (F,H). Error bars indicate SEM over n=8 images (E&F), n=5 (G&H). Scale bars: 100um.

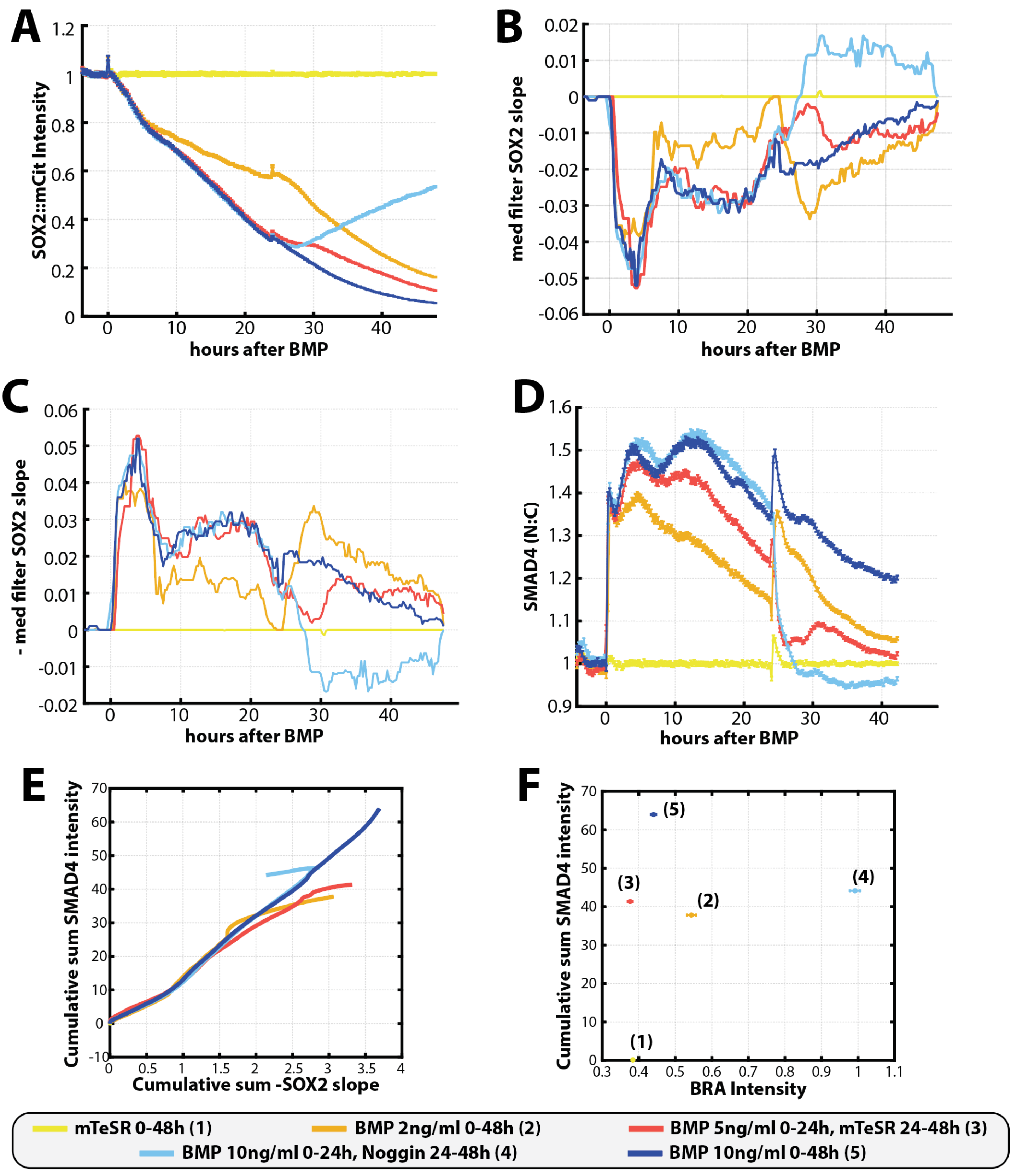
A
SOX2::mCit Intensity
hours after BMP
B
med filter SOX2 slope
hours after BMP
C
- med filter SOX2 slope
hours after BMP
D
SMAD4 (N:C)
hours after BMP
E
Cumulative sum SMAD4 intensity
Cumulative sum -SOX2 slope
F
Cumulative sum SMAD4 intensity
BRA Intensity
(1)
(2)
(3)
(4)
(5)
mTeSR 0-48h (1)
BMP 2ng/ml 0-48h (2)
BMP 5ng/ml 0-24h, mTeSR 24-48h (3)
BMP 10ng/ml 0-24h, Noggin 24-48h (4)
BMP 10ng/ml 0-48h (5)

**Figure S9. Downregulation of SOX2 by SMAD4. (A)** SOX2::mCitrine average nuclear intensity under the indicated treatments. **(B)** Median filter of the numerically computed slope of the curves in (A). **(C)** Median filter of the numerically computed slope of the curves in (A) multiplied by -1. **(D)** GFP::SMAD4 average nuclear:cytoplasmic intensity ratio for the same treatments as in (A-C). This figure shows the same data set shown in Fig. S3D. **(E)** Cumulative sum of SMAD4 intensity in time obtained from (D) versus the cumulative sum of minus the slope of SOX2 in time from (C). **(F)** Mean and SEM of BRA intensity obtained from immunostaining the experiment shown in (D) after 48 hours the corresponding treatments versus the cumulative sum of SMAD4 intensity in time obtained from (D). SEM over n=8 images.

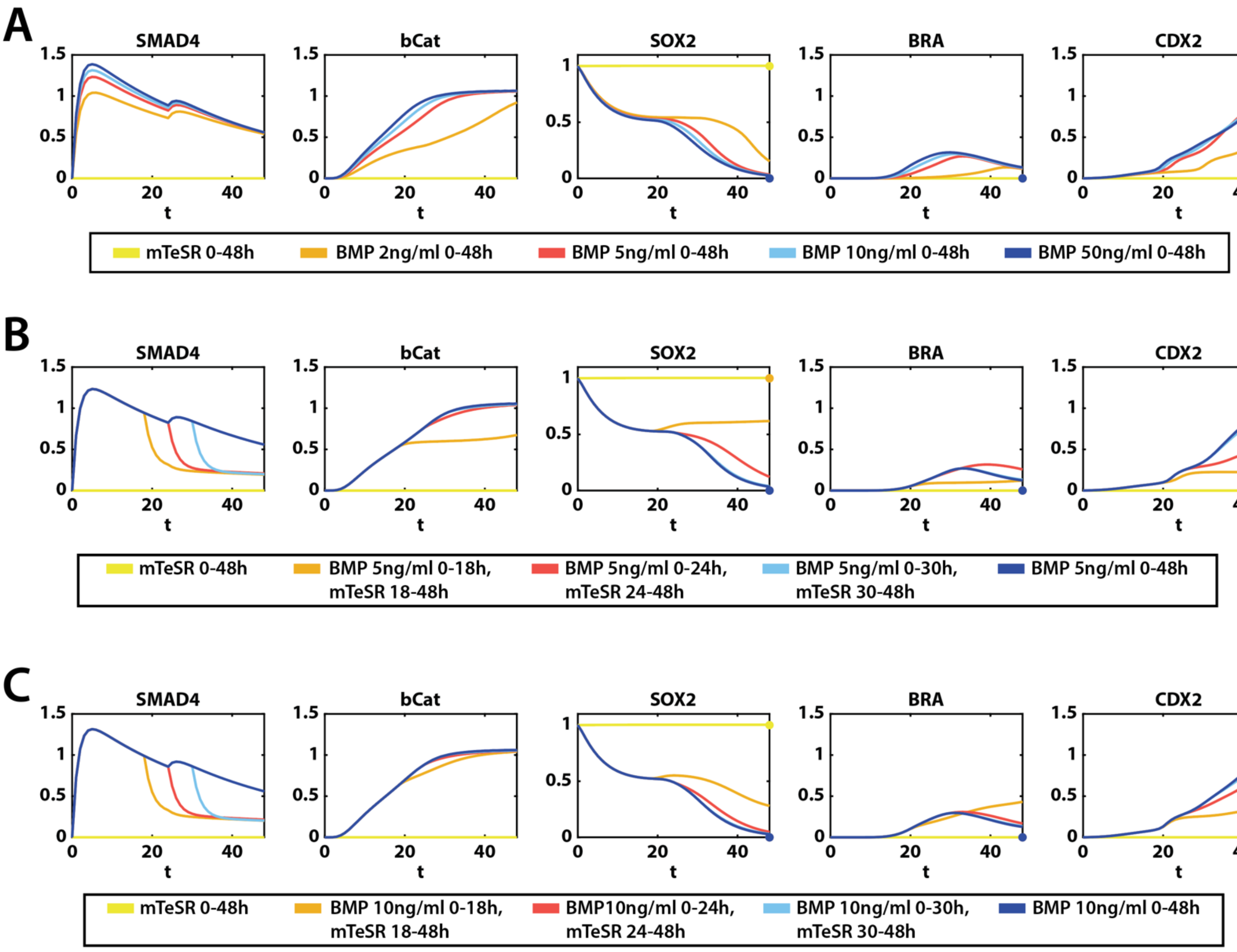
A
SMAD4
bCat
SOX2
BRA
CDX2
t
mTeSR 0-48h
BMP 2ng/ml 0-48h
BMP 5ng/ml 0-48h
BMP 10ng/ml 0-48h
BMP 50ng/ml 0-48h
B
SMAD4
bCat
SOX2
BRA
CDX2
t
mTeSR 0-48h
BMP 5ng/ml 0-18h, mTeSR 18-48h
BMP 5ng/ml 0-24h, mTeSR 24-48h
BMP 5ng/ml 0-30h, mTeSR 30-48h
BMP 5ng/ml 0-48h
C
SMAD4
bCat
SOX2
BRA
CDX2
t
mTeSR 0-48h
BMP 10ng/ml 0-18h, mTeSR 18-48h
BMP10ng/ml 0-24h, mTeSR 24-48h
BMP 10ng/ml 0-30h, mTeSR 30-48h
BMP 10ng/ml 0-48h

**Figure S10. CFN model validations. (A)** Simulated SMAD4, bCat, SOX2, BRA and CDX2 dynamics from the deterministic CFN model for the indicated experimental conditions. Dots represent the data summaries obtained as explained in the Supplementary Information. **(B)** Simulated SMAD4, bCat, SOX2, BRA and CDX2 dynamics from the deterministic CFN model for the indicated experimental conditions. Dots represent the data summaries obtained as explained in the Supplementary Information. **(C)** Simulated SMAD4, bCat, SOX2, BRA and CDX2 dynamics from the deterministic CFN model for the indicated experimental conditions. Dots represent the data summaries obtained as explained in the Supplementary Information

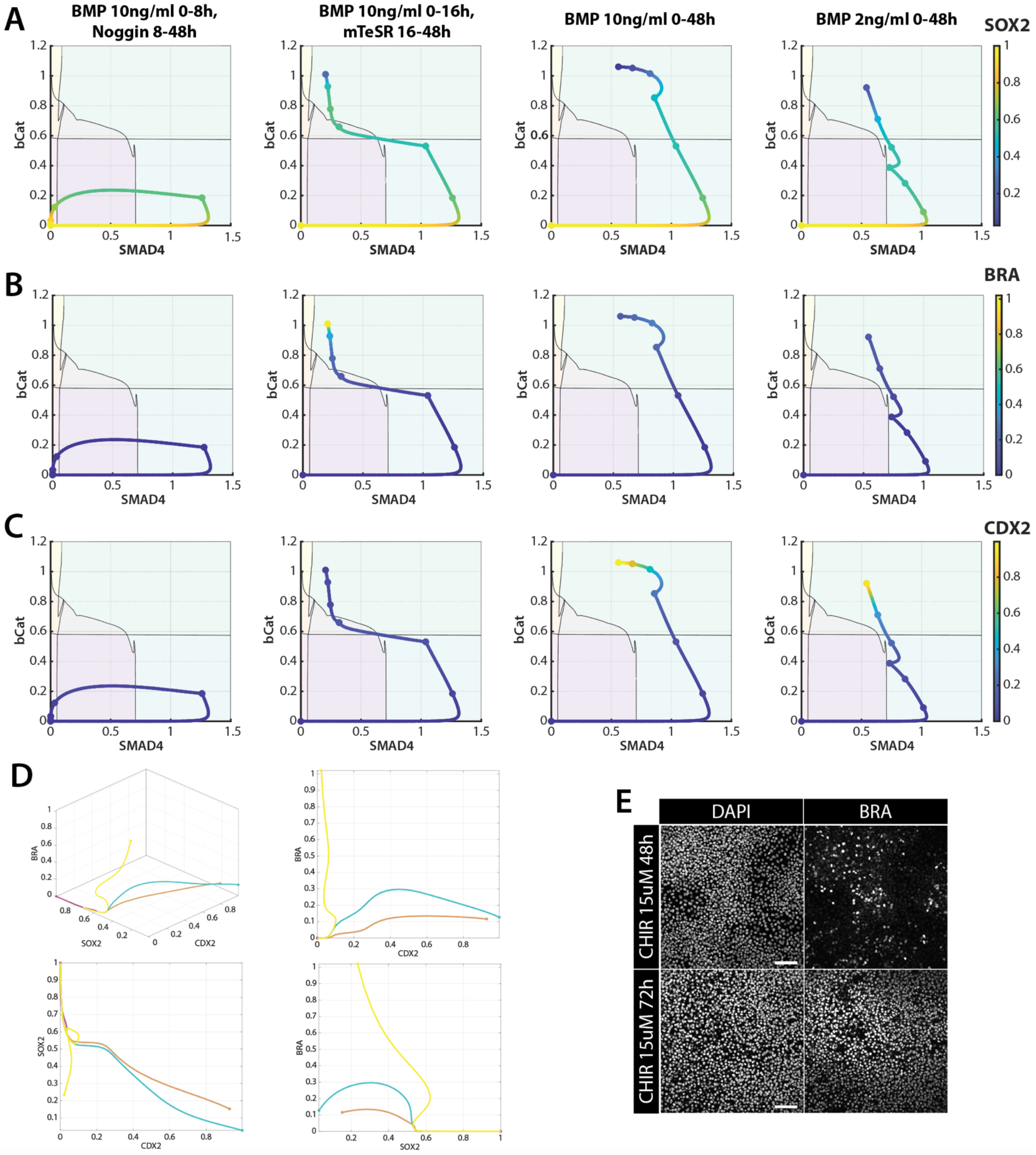
A
BMP 10ng/ml 0-8h, Noggin 8-48h
BMP 10ng/ml 0-16h, mTeSR 16-48h
BMP 10ng/ml 0-48h
BMP 2ng/ml 0-48h
SOX2
B
BRA
C
CDX2
bCat
SMAD4
D
BRA
SOX2
CDX2
E
DAPI
BRA
CHIR 15uM 48h
CHIR 15uM 72h

**Figure S11. Cell fate is a combinatorial response to BMP and WNT. (A)** Simulated signaling trajectories on the fate map for the indicated BMP treatment. Each trajectory at time t is colored by the simulated SOX2 expression value at time t. 0, 8, 16, 24, 32, 40, and 48 hour timepoints are highlighted with bigger sized dots. **(B)** Simulated signaling trajectories on the fate map for the indicated BMP treatment. Each trajectory at time t is colored by the simulated BRA expression value at time t. 0, 8, 16, 24, 32, 40, and 48 hour timepoints are highlighted with bigger sized dots. **(C)** Simulated signaling trajectories on the fate map for the indicated BMP treatment. Each trajectory at time t is colored by the simulated CDX2 expression value at time t. 0, 8, 16, 24, 32, 40, and 48 hour timepoints are highlighted with bigger sized dots. **(D)** Simulated expression dynamics for the conditions: mTeSR (pink), BMP 10ng/ml 0-48h (cyan), BMP 10ng/ml 0-16, mTeSR 16-48h (yellow), BMP 2ng/ml 0-48h (orange). End points are highlighted with a dot. **(E)** Example images of immunofluorescence for DAPI and BRA after 48h (top) and 72h (bottom) of the indicated CHIR treatment.

### Supplemental Information

Combinatorial interpretation of BMP and WNT allows BMP to act as a morphogen in time but not in concentration.

Elena Camacho-Aguilar[1], Sumin Yoon[1], Miguel A. Ortiz-Salazar[1], and Aryeh Warmflash[1,2]

[1]Department of Biosciences, Rice University, Houston, Texas, United States of America, [2]Department of Bioengineering, Rice University, Houston, Texas, United States of America

November 1, 2022

### 1 Cell classification

Given an experiment, which comprises a set of experimental conditions or treatments, individual cells were classified as All low, SOX2+, BRA+, CDX2+, or a mix of two or three markers depending on the expression levels of three reference markers SOX2, BRA, and CDX2 (or ISL1), which in our experiments mark the pluripotent, mesoderm and extraembryonic fates, respectively, and are measured by immunofluorescence staining.

Due to batch effects, each experiment was classified separately. However, all experiments contained negative and positive controls for each of the three reference markers. These conditions are shown in Table SI1.

Given an experiment $e$, comprising a set of $N$ experimental conditions or treatments, immunofluorescence images were analyzed as discussed in the Methods section of the article. Following this image analysis process, we obtained a set of data $D^e = \{(S^e_{t,c}, B^e_{t,c}, C^e_{t,c}) : t \in \{1, \dots, N\}, c \in \{1, \dots, M_t\}\}$, where $(S^e_{t,c}, B^e_{t,c}, C^e_{t,c})$ is the DAPI-normalised SOX2, BRA and CDX2 intensity for a cell $c$ grown under treatment $t$, and $M_t$ is the total number of measured cells grown under treatment $t$.

The mean protein expressions of each of the three reference markers were then normalized by dividing their values by the 99.5th percentile point of the pooled data for all the experimental conditions, obtaining $Z^e = \{(ZS^e_{t,c}, ZB^e_{t,c}, ZC^e_{t,c}) : t \in \{1, \dots, N\}, c \in \{1, \dots, M_t\}\}$, where

| | Negative control | Positive control |
|---|---|---|
| SOX2 | BMP $\geq$ 10ng/ml 0-48h | mTeSR 0-48 |
| BRA | mTeSR 0-48 | BMP 10ng/ml 0-16h, mTeSR 16-48 OR<br>BMP 10ng/ml 0-30h, Noggin 30-48 |
| CDX2 | mTeSR 0-48 | BMP $\geq$ 10ng/ml 0-48h |

Table SI1: Table of negative and positive controls for the expression of each reference marker.

$$ZS^e_{t,c} = \frac{S^e_{t,c}}{P995(\{S^e_{t,c} : t \in \{1, \ldots, N\}, c \in \{1, \ldots, M_t\}\})}, \quad (1)$$

and $P995(\Omega)$ is the function returning the 99.5th percentile of the set $\Omega$. $ZB^e_{t,c}$ and $ZC^e_{t,c}$ are similarly defined.

We also defined a background intensity for each marker as the 95th percentile point of the normalized values coming from the negative control population for each marker. For example,

$$ZS0^e = P95(\{ZS^e_{t,c} : t = \text{Negative control for SOX2}, c \in \{1, \ldots, M_t\}\}), \quad (2)$$

where and $P95(\Omega)$ is the function returning the 95th percentile of the set $\Omega$. $ZB^e_{t,c}$ and $ZC^e_{t,c}$ are similarly defined. $ZB0^e$ and $ZC0^e$ are similarly defined.

Finally, each cell $c$ under treatment $t$ is classified depending on its normalized value with respect to the background intensity:

- If $ZS^e_{t,c} < ZS0^e$, $ZB^e_{t,c} < ZB0^e$, and $ZC^e_{t,c} < ZC0^e$, i.e. both its SOX2, BRA and CDX2 intensities fall bellow the limits set by the negative controls, the cell will be classified as "All Low".
- Otherwise, we look at how much each marker contributes to the *total* immunoflorescence intensity of the cell. I.e., we define $\tilde{S}^e_{t,c} = ZS^e_{t,c}/(ZS^e_{t,c} + ZB^e_{t,c} + ZC^e_{t,c})$, and similarly $\tilde{B}^e_{t,c}$ and $\tilde{C}e_{t,c}$. It is satisfied that $\tilde{S}^e_{t,c} + \tilde{B}^e_{t,c} + \tilde{C}^e_{t,c} = 1$. If one gene defines more than 50% of the immunofluorescence intensity of the cell, the cell is classified as the corresponding type of cell. For example, if the new normalized immunofluorescence of a cell is given by $(\tilde{S}^e_{t,c}, \tilde{B}^e_{t,c}, \tilde{C}^e_{t,c}) = (0.9, 0.05, 0.05)$, the cell will be classified as "SOX2+" or pluripotent, as SOX2 contributes to the 90% of its expression.
- Otherwise, we looked for the two genes that had the biggest proportions of normalized expression. If the sum of the two values was bigger than 80%, the cell was classified as the mix of the two fates. For example, if the normalized immunofluorescence of a cell is given by $(\tilde{S}^e_{t,c}, \tilde{B}^e_{t,c}, \tilde{C}^e_{t,c}) = (0.5, 0.15, 0.35)$, the cell will be classified as "SOX2/CDX2+," i.e. a mix of SOX2 and CDX2 expression. These cells are probably in transition between two fates.
- In the case that a cell is not classified in the previous steps, it will classified as a cell with a mix of all markers. These cells are probably also in transition between fates.

### 2 Mathematical models

#### 2.1 Signaling model

We first model the response of hPSCs to the signals: the exogenously applied BMP4, and the endogenously secreted WNT. As described in the main text, this signaling model is comprised of two sub-systems of differential equations that model the BMP and Wnt responses, respectively, and are connected via BMP4 activation of Wnt, which reflects the transcriptional activation of *WNT3* by BMP signaling. Here we describe the derivation of the two sub-systems.

| Parameter | Value | Parameter | Value |
|---|---|---|---|
| $b_{aff}$ | 0.1 | $1/\tau_2$ | 0.05 |
| $\gamma$ | 10 | $1/\tau_3$ | 0.025 |
| $\delta$ | 2 | $\alpha_1$ | 30 |
| $\beta$ | 0.1 | $\alpha_2$ | 1 |
| $\delta_I$ | 0.01 | $\alpha_3$ | 1 |
| $1/\tau_1$ | 0.075 | $r_b$ | 0.1824 |

Table SI2: Table of parameter values used when simulating model in Eq 3

###### 2.1.1 Dynamics of BMP response

When exogenously stimulated, BMP dynamics show a rapid rise followed by a slow decay likely to to the degradation of activated receptors. This BMP response (encoded in the variable SMAD4 in Eq. 3) was simulated using the model proposed in [1].

$$\begin{cases} \dfrac{d\text{BMP}}{dt} = \dfrac{1}{\tau_1}(-(1-\text{Nog})\, b_{aff}\text{BMP} \times \text{BMPR} + \gamma\text{SMAD4}) \\ \\ \dfrac{d\text{BMPR}}{dt} = \dfrac{1}{\tau_2}(\beta - (\delta_I + (1-\text{Nog})\, b_{aff}\text{BMP}) \times \text{BMPR} + \gamma\text{SMAD4}) \\ \\ \dfrac{d\text{SMAD4}}{dt} = \dfrac{1}{\tau_3}((1-\text{Nog})\, b_{aff}\text{BMP} \times \text{BMPR} - (\gamma+\delta)\text{SMAD4}) \end{cases} \tag{3}$$

In this model of three variables, BMP ligands (modeled through variable BMP in Eq. 3) bind to the receptor complex (modeled through variable BMPR in Eq. 3) with rate $b_{aff}$ , which in turn activates SMAD4 but also enhances the degradation of the BMP receptor. For simplicity, in the model, the binding of BMP and BMPR directly produces activated SMAD4, and we do explicitly model the activated receptor complex or the R-SMADs. The parameter $b_{aff}$ represents the rate of binding of the BMP ligands and unoccupied receptors, $\beta$ represents the production of receptors, $\gamma$ represents the return of receptors to the unoccupied state, $\delta_I$ represents the degradation rate of unoccupied receptors, $\delta$ the degradation rate of SMAD4 due to degradation of occupied receptors, and $\tau_i$ are time constants. The presence of the inhibitor Noggin is modeled through a parameter Nog which can take value 1 if Noggin is present in the media, therefore preventing the BMP ligand from binding to the available receptors.

The system was simulated with initial condition BMP= $F_{BMP}(bmp)$, BMPR=$\beta/\delta_I$, and SMAD4=0, where $F_{BMP}(bmp) = \alpha_1/(1 + (\alpha_2/bmp)^{\alpha_3})$ is a function of the experimental BMP4 ligand concentration $bmp$ used, and the parameter values given in Table SI2. If $bmp = 0$, $F_{BMP}(bmp)$ is defined to be 0.

An experimental media change at time $t' > 0$ is simulated by changing the value of the variable BMP at time $t'$ to be equal to $F_{BMP}(bmp)$ if the BMP4 ligand concentration introduced $bmp$ is greater than 0, or $r_b F_{BMP}(bmp)$ if the media change is a removal of the original BMP by either an mTeSR or Noggin media change, and $r_b$ is a parameter that reflects the proportion of BMP that was removed by the media change. If the experimental media change also introduces Noggin to the media at time $t'$, the variable BMPR at time $t'$ is changed to 0. To mimic the experiments, a media change was always simulated at 24 hours.

Simulations of this model for different BMP4 durations are shown in Fig. SI1 and Fig. S6A, and simulations for different concentrations are shown in Fig. S6B.

###### 2.1.2 Dynamics of the WNT response

The model for the WNT response is comprised of two equations that model WNT ligand (variable WNT in Eq. 4) and nuclear $\beta$-catenin (variable $\beta$Cat in Eq. 4). WNT ligand expression is activated by

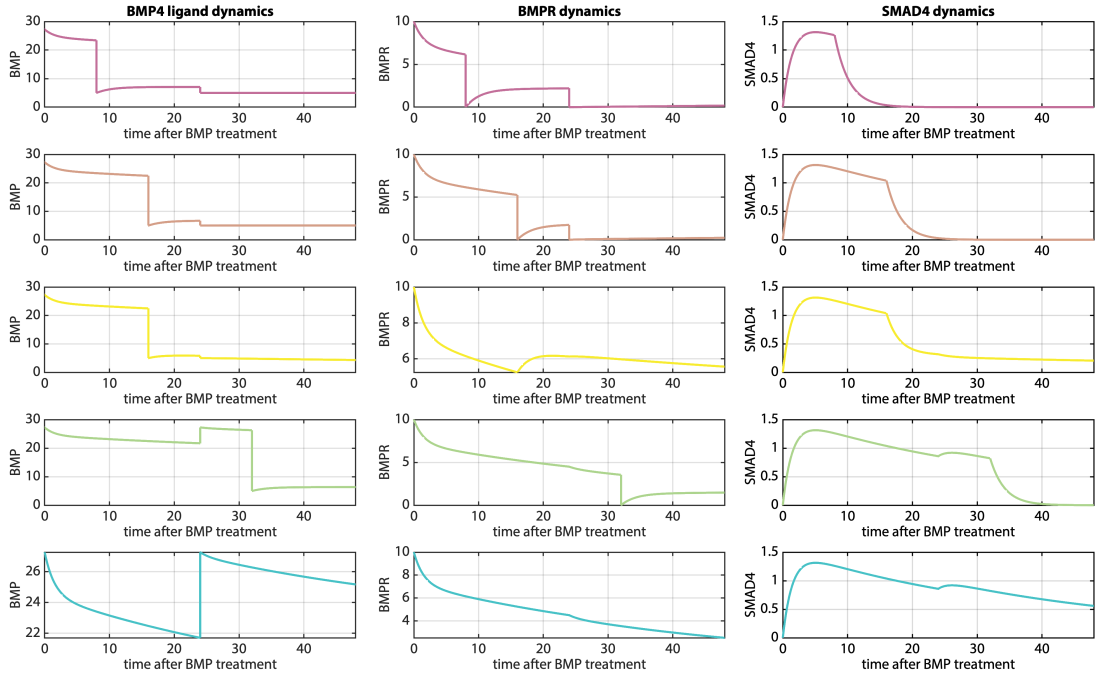

Figure SI1: From top to bottom, simulations of BMP dynamics for the conditions: BMP 10ng/ml 0-8h, Noggin 8-48h; BMP 10ng/ml 0-16h, Noggin 16-48h; BMP 10ng/ml 0-16h, mTeSR 16-48h; BMP 10ng/ml 0-32h, Noggin 32-48h; BMP 10ng/ml 0-48h.

SMAD4, which activates nuclear $\beta$-catenin which, in turn, further activates WNT ligand expression.

$$\begin{cases} \dfrac{d\mathrm{WNT}}{dt} = (1-\mathrm{IWP2})\left(a_{SW}\dfrac{\mathrm{SMAD4}^{n_{SW}}}{K_{SW}^{n_{SW}}+\mathrm{SMAD4}^{n_{SW}}} + a_{WW}\dfrac{\beta\mathrm{Cat}^{n_{WW}}}{K_{WW}^{n_{WW}}+\beta\mathrm{Cat}^{n_{WW}}}\right) - \delta_W \mathrm{WNT} \\ \\ \dfrac{d\beta\mathrm{Cat}}{dt} = a_{W\beta}\dfrac{(\mathrm{WNT}+a_{C\beta}\mathrm{CHIR})^{n_{W\beta}}}{K_{W\beta}^{n_{W\beta}}+(\mathrm{WNT}+a_{C\beta}\mathrm{CHIR})^{n_{W\beta}}} - \delta_\beta \beta\mathrm{Cat}, \end{cases} \quad (4)$$

where the parameters $a_i$, $n_i$ and $K_i$ (with $i \in \{SW, WW, W\beta\}$) are the production rates, Hill coefficients and dissociation constants respectively, $\delta_i$ with (with $i \in \{W, \beta\}$) are the degradation rates, IWP2 models the effect of introducing the small-molecule inhibitor IWP2 in the media, and CHIR models the effect of using the small-molecule GSK3$\beta$ inhibitor CHIR99021 as a WNT signaling stimulator.

The data shown in Fig. 3H and S3B-C suggests that the system should have at least two stable equilibria when BMP signal is low since, under a short pulse of BMP signal, $\beta$Cat dynamics converge to a low level, while under long pulses of BMP (such as one of 32 hours), $\beta$Cat dynamics converge to a high level.

In order to explore which parameters values can reproduce the observed WNT bistability, we first study the simpler equation in Eq. 5, which results from assuming (1) that $\beta$Cat dynamics are sufficiently fast that the $\beta$Cat can be assumed to be at equilibrium, (2) that the level of BMP signaling reflected in SMAD4 is a fixed control parameter, and that IWP2 and CHIR are not present in the media and therefore are equal to 0.

$$\frac{d\mathrm{WNT}}{dt} = \left(a_{SW}\frac{\mathrm{SMAD4}^{n_{SW}}}{K_{SW}^{n_{SW}}+\mathrm{SMAD4}^{n_{SW}}} + a_{WW}\frac{WNT^{n_{WW}}}{K_{WW}^{n_{WW}}+WNT^{n_{WW}}}\right) - \delta_W \mathrm{WNT} \quad (5)$$

| Parameter | Value |
|---|---|
| $a_{SW}$ | 0.15 |
| $n_{SW}$ | 2 |
| $K_{SW}$ | 1.55 |
| $a_{WW}$ | 0.6496 |
| $n_{WW}$ | 3 |
| $K_{WW}$ | 1.45 |
| $\delta_W$ | 0.07 |

| Parameter | Value |
|---|---|
| $a_{W\beta}$ | 2.52 |
| $n_{W\beta}$ | 2.5 |
| $K_{W\beta}$ | 0.59 |
| $\delta_\beta$ | 2.31 |

Table SI3: Table of parameter values used when simulating model in Eqs. (5, 4) to produce the bifurcation plots in Fig. SI2 and Fig. SI3

We find that, for the parameters in Table SI3, Eq. 5 can have 1, 2 or 3 positive critical points, and undergoes a saddle-node bifurcation depending on the value of SMAD4 which can be viewed as a control parameter (Fig. SI2). When SMAD4 is high (SMAD4 = 1), the system has just one stable equilibrium point at high WNT levels. After a pulse of BMP, when SMAD4 returns to low values, the value of WNT that the system will tend to will depend on whether the WNT value at the time of BMP withdrawal is higher or lower than the positive unstable critical point.

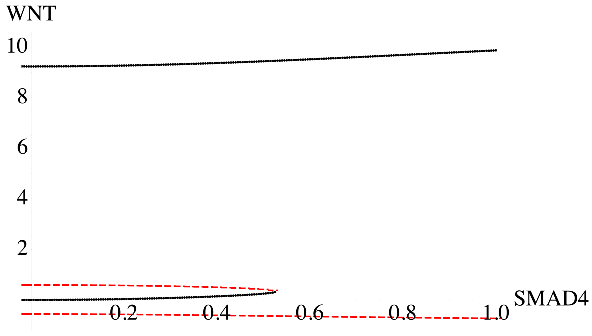

Figure SI2: Bifurcation plot showing the stable (black dots) and unstable (red dashes) critical points of the system in Eq. (5) with parameter values in Table SI3 for different values of SMAD4.

Now, fixing the parameters used for simulating Eq. 5, we study the complete WNT system in Eq. (4). In order to study how the WNT stability depends upon the parameter SMAD4, we assume IWP2=CHIR=0. Our goal is to find a bistable system which reproduces that, under short BMP4/SMAD4 pulses, WNT and $\beta$Cat converge to a low equilibrium point, while under long BMP4/SMAD4 pulses, WNT and $\beta$Cat converge to a high equilibrium point. An interesting data feature to consider is the dynamics observed under a 16 hour pulse of BMP 10ng/ml followed by mTeSR with or without Noggin (Fig. S6B-C). The data shows that if a 16 hour pulse of BMP4 is followed by mTeSR with Noggin, $\beta$Cat converges to a low equilibrium point, while if no Noggin is added after the 16 hour pulse, $\beta$Cat converges to a high equilibrium point. Translating this observation into the model, this means that, the WNT and $\beta$Cat trajectory at t=16 should be on the side of the saddle for SMAD4=0 (i.e. Noggin media change) for which the trajectory converges to the low WNT/$\beta$Cat equilibrium. While it should have crossed this saddle for SMAD4=$r_b F_{BMP}(10)$ (i.e. the amount of BMP remaining in the media for a media change without Noggin), so that it converges to the high WNT/$\beta$Cat equilibrium. Taking this into account, and joining the BMP and WNT sub-systems in Eqs. (3, 4) into a system of five ordinary differential equations, we find parameters that reproduce these features in our model (Figs. SI3 and SI4). These parameters are given in Tables SI2 and SI3. Simulations for different pulses and concentrations are shown in Fig. S6C-D.

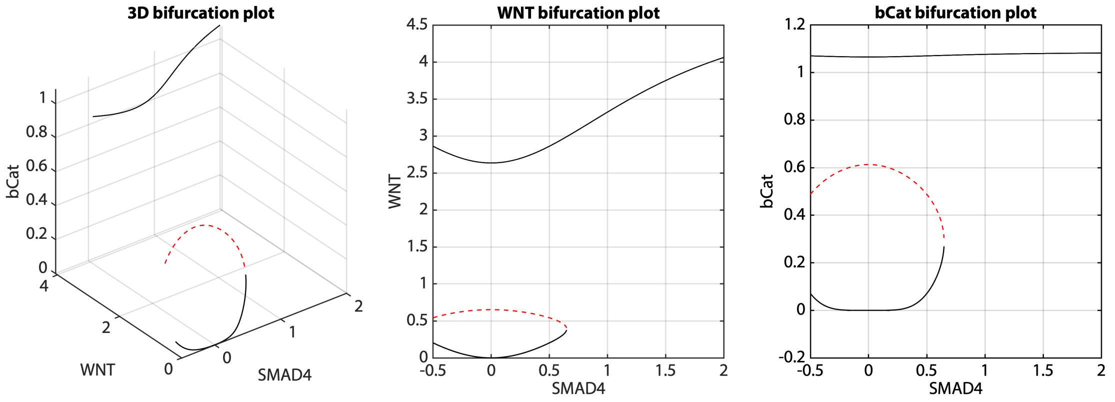

Figure SI3: (Left) Bifurcation plot showing the stable (black dots) and unstable (red dashes) critical points of the system in Eq. 4 with parameter values in Table SI3 for different values of SMAD4. (Center) Projection of the bifurcation set on the (WNT,SMAD4) plane. (Right) Projection of the bifurcation set on the ($\beta$Cat,SMAD4) plane.

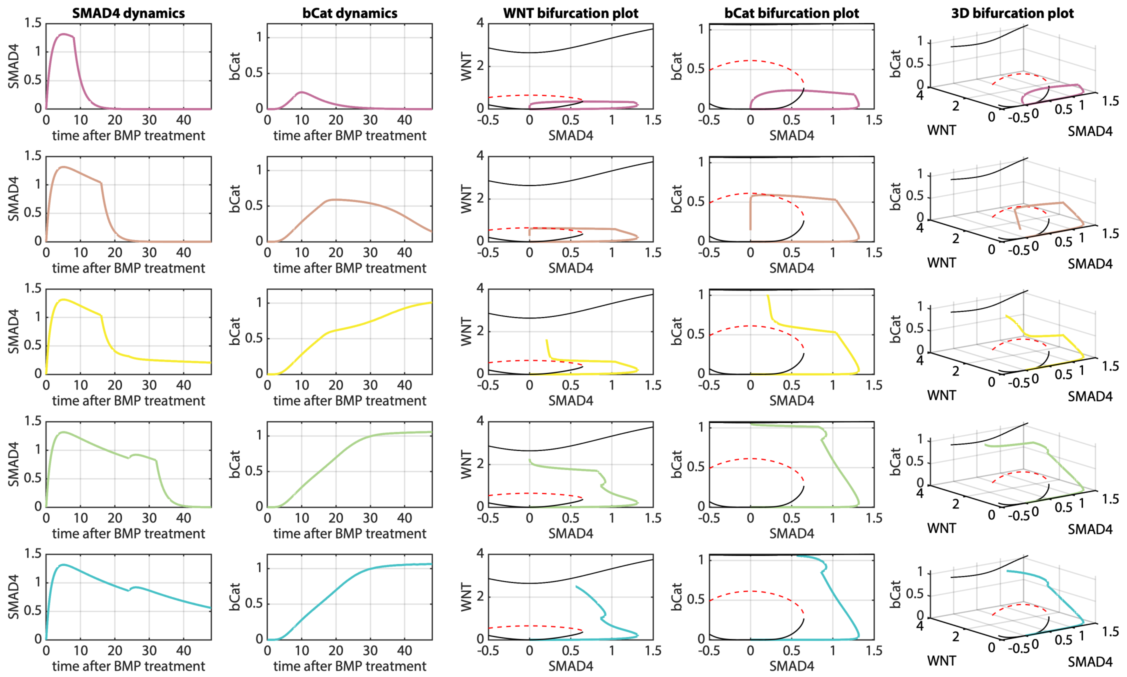

Figure SI4: From top to bottom, simulations of SMAD4 and $\beta$Cat dynamics for the conditions: BMP 10ng/ml 0-8h, Noggin 8-48h; BMP 10ng/ml 0-16h, Noggin 16-48h; BMP 10ng/ml 0-16h, mTeSR 16-48h; BMP 10ng/ml 0-32h, Noggin 32-48h; BMP 10ng/ml 0-48h.

#### 2.2 Cell Fate Network model

To understand how signal dynamics controlled the observed cell fate transitions, we created a minimal cell fate network (CFN) model, where the nodes of the network correspond to the cell fates observed, i.e. pluripotent, mesodermal, and extra-embryonic, characterized by high SOX2, BRA, or CDX2 expression, respectively. As an input, the CFN model will take the SMAD4 and $\beta$Cat trajectories from the signaling model to simulate how cells change their fate in time as a function of the signaling received. For simplicity, we modeled SOX2, BRA and CDX2 expression in time as a proxy for the corresponding cell fates.

We propose that the cell fate network should be the one given in Fig. SI5, which is modeled by the system of three ordinary differential equations in Eq. 6. Here we explain each interaction included in the CFN model:

**Mutual inhibition between the cell fate nodes** We included mutually repressive interactions between SOX2, BRA and CDX2 to generate mutually exclusive cell states. These inhibitions are modeled through Hill functions with Hill coefficients and dissociation constants $n_{ij}$, $K_{ij}$, where $i, j \in \{S, B, C\}$.

**Pluripotency (SOX2) inhibition by by BMP signal through SMAD4** The SOX2 expression dynamics observed in Fig. 5G and S9 suggested a downregulation of SOX2 by BMP, through a SMAD intermediate, as SOX2 expression begins to decay rapidly upon BMP treatment, at which stage neither Wnt signaling as reflected in nuclear $\beta$-catenin, nor mesoderm or ExE markers are yet upregulated (Fig 3H, 5A-C). In fact, early SOX2 decay dynamics show a very high correlation with the integral of SMAD4 in time (Fig. S9E). We therefore include this inhibition in our model through a Hill function with Hill coefficient $n_{SmS}$ and dissociation constant $K_{SmS}$.

**Mesoderm (BRA) induction by WNT signaling through $\beta$Cat** As we and previous studies have shown, WNT signals are necessary for BMP-induced mesoderm formation [2, 3]. We considered that mesoderm markers are therefore upregulated through $\beta$-catenin activation by WNT, and modelled the activation of BRA expression through a Hill function of $\beta$-catenin levels with Hill coefficient $n_{\beta B}$ and dissociation constant $K_{\beta B}$.

**ExE (CDX2) induction by SMAD4** Our data and previous studies have shown that BMP activation of SMAD complexes leads to transcription of ExE markers, such as CDX2 [4]. Therefore we model this activation through a Hill function with Hill coefficient $n_{SmC}$ and dissociation constant $K_{SmC}$.

**ExE state (CDX2) auto-regulation** Our data shows that BMP induction of longer than 30 hours, if replaced by mTeSR alone, or than 40 hours, if withdrawn by mTeSR+Noggin, result in ExE differentiation. Therefore, the ExE node needs to become self-sustained. We model this ExE auto-regulation through an additive auto-activation with Hill coefficient $n_{CC}$ and dissociation constant $K_{CC}$.

Finally, we include parameters for the activation, $a_i$ and degradation $\delta_j$ of each gene, where $i \in \{S, B, C, CC\}$ and $j \in \{S, B, C\}$.

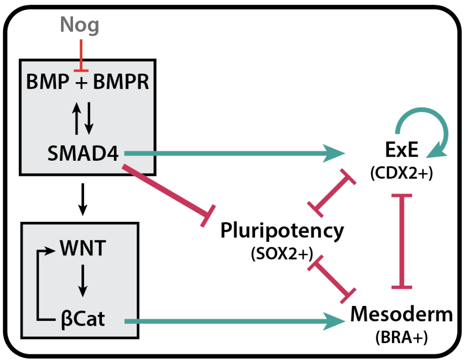

Figure SI5: From top to bottom, simulations of SMAD4 and bCat dynamics for the conditions: BMP 10ng/ml 0-8h, Noggin 8-48h; BMP 10ng/ml 0-16h, Noggin 16-48h; BMP 10ng/ml 0-16h, mTeSR 16-48h; BMP 10ng/ml 0-32h, Noggin 32-48h; BMP 10ng/ml 0-48h.

$$
\begin{cases}
\dfrac{d\mathrm{SOX2}}{dt} = \dfrac{a_S}{(K_{SmS}^{n_{SmS}} + \mathrm{SMAD4}^{n_{SmS}})(1 + (\mathrm{CDX2}/K_{CS})^{n_{CS}} + (\mathrm{BRA}/K_{BS})^{n_{BS}})} - \delta_S \mathrm{SOX2} \\[2ex]
\dfrac{d\mathrm{BRA}}{dt} = \dfrac{a_B \beta Cat^{n_{\beta B}}}{(K_{\beta B}^{n_{\beta B}} + \beta Cat^{n_{\beta B}})(1 + (\mathrm{SOX2}/K_{SB})^{n_{SB}} + (\mathrm{CDX2}/K_{CB})^{n_{CB}})} - \delta_B \mathrm{BRA} \\[2ex]
\dfrac{d\mathrm{CDX2}}{dt} = \dfrac{a_C \mathrm{SMAD4}^{n_{SmC}}}{(K_{SmC}^{n_{SmC}} + \mathrm{SMAD4}^{n_{SmC}})(1 + (\mathrm{SOX2}/K_{SC})^{n_{SC}} + (\mathrm{BRA}/K_{BC})^{n_{BC}})} + \dots \\[2ex]
\qquad \qquad \dots + \dfrac{a_{CC} \mathrm{CDX2}^{n_{CC}}}{K_{CC}^{n_{CC}} + \mathrm{CDX2}^{n_{CC}}} - \delta_C \mathrm{CDX2}
\end{cases}
\tag{6}
$$

#### 2.3 Fitting of the deterministic CFN model

Our goal was now to find parameters that could reproduce the cell fates as a function of the signaling dynamics observed. To do so, we first defined a standarized format for the data we aimed to reproduce.

##### 2.3.1 Data summaries

In order to fit the model, we summarized the data of each experimental condition into a vector (SOX2, BRA, CDX2) containing a representative SOX2, BRA and CDX2 expression for that experimental condition, that could be compared to a simulation of our deterministic CFN model.

Given an experimental condition, we first attempted averaging the normalized values obtained, as explained in Section 1, of the most prominent type of cell (pluripotent, mesoderm, ExE or mixed) present in that experimental condition. However, we could still notice some batch effects in this summary and a deterministic model may be better suited to capture the mode of the data (i.e. the most likely cell fate) rather than the mixture of fates defined by the average. For that reason, we further simplified the data corresponding to an experimental condition to (1,0,0), (0,1,0), (0,0,1) or (0.5,0.5,0.5) if the most prominent type of cell in that experimental condition was pluripotent, mesoderm, ExE or mixed, respectively. The full dataset containing fractions of each cell fate was used in fitting the stochastic model, as described below.

Data on the expression of SOX2, CDX2, and BRA captured in time at 16h, 24h, 32h and 48h after treatment (Fig. 5 and S7) required a different simplification in order to capture the dynamics of the transitions as well as the final fates. In this case, the values of the markers SOX2 and CDX2 were further normalised by the maximum value of that marker in the whole experiment if this resulting value was bigger than 0.3, otherwise it was set to 0. The resulting summaries of the data are provided in the file **DataExperiments.xlsx**.

##### 2.3.2 Data fitting

First, we found a candidate parameter set that could reproduce a subset of the experimental conditions (see Table SI4 and Fig. SI6).

From the experimental data in **DataExperiments.xlsx**, we selected a subset of conditions $\Omega = \{\Omega_S, \Omega_B, \Omega_C, \Omega_M, \Omega_0\}$ to fit the model, where $\Omega_S, \Omega_B, \Omega_C, \Omega_M, \Omega_0$ are subsets of experimental conditions where the predominant cell type is pluripotent (SOX2+), mesoderm (BRA+), ExE (CDX2+), and mixed fates, respectively, and $\Omega_0$ are control conditions where either BMP or WNT signals are selectively present. See Tables SI5, SI6, and SI7.

We defined a cost function to compare the experimental and simulated data in the following way. Let $\boldsymbol{x}_c^{exp} = (\mathrm{SOX2}_c^{exp}, \mathrm{BRA}_c^{exp}, \mathrm{CDX2}_c^{exp})$ and $\boldsymbol{x}_c^{num}(\boldsymbol{\theta}) = (\mathrm{SOX2}_c^{num}, \mathrm{BRA}_c^{num}, \mathrm{CDX2}_c^{num})$ be the simplified experimental expression and corresponding simulation for condition $c$ given parameter $\boldsymbol{\theta}$. The cost function for parameter $\boldsymbol{\theta}$ is given by:

$$
L(\boldsymbol{\theta}) = \frac{1}{\sum_{i\in\{S,C,B,M,0\}} \omega_i} \sum_{i\in\{S,C,B,M,0\}} \frac{\omega_i}{N_i} \sum_{c\in\Omega_i} \|\boldsymbol{x}_c^{exp} - \boldsymbol{x}_c^{num}(\boldsymbol{\theta})\|^2, \tag{7}
$$

| Parameter | Value |
|---|---|
| $a_S$ | 0.1364 |
| $n_{CS}$ | 2 |
| $K_{CS}$ | 0.4 |
| $n_{BS}$ | 2 |
| $K_{BS}$ | 0.375 |
| $n_{SmS}$ | 1 |
| $K_{SmS}$ | 0.8 |
| $\delta_S$ | 0.165 |

| Parameter | Value |
|---|---|
| $a_B$ | 0.375 |
| $n_{\beta B}$ | 4 |
| $K_{\beta B}$ | 0.6 |
| $n_{SB}$ | 3 |
| $K_{SB}$ | 0.4 |
| $n_{CB}$ | 3 |
| $K_{CB}$ | 0.0673 |
| $\delta_B$ | 0.18 |

| Parameter | Value |
|---|---|
| $a_C$ | 0.15 |
| $n_{SmC}$ | 2 |
| $K_{SmC}$ | 0.9 |
| $n_{SC}$ | 3 |
| $K_{SC}$ | 0.3818 |
| $n_{BC}$ | 2 |
| $K_{BC}$ | 0.5625 |
| $a_{CC}$ | 0.5 |
| $n_{CC}$ | 3 |
| $K_{CC}$ | 0.3 |
| $\delta_B$ | 0.45 |

Table SI4: Table of initial candidate parameter values used when simulating model in Eqs. (6) to produce Fig. SI6

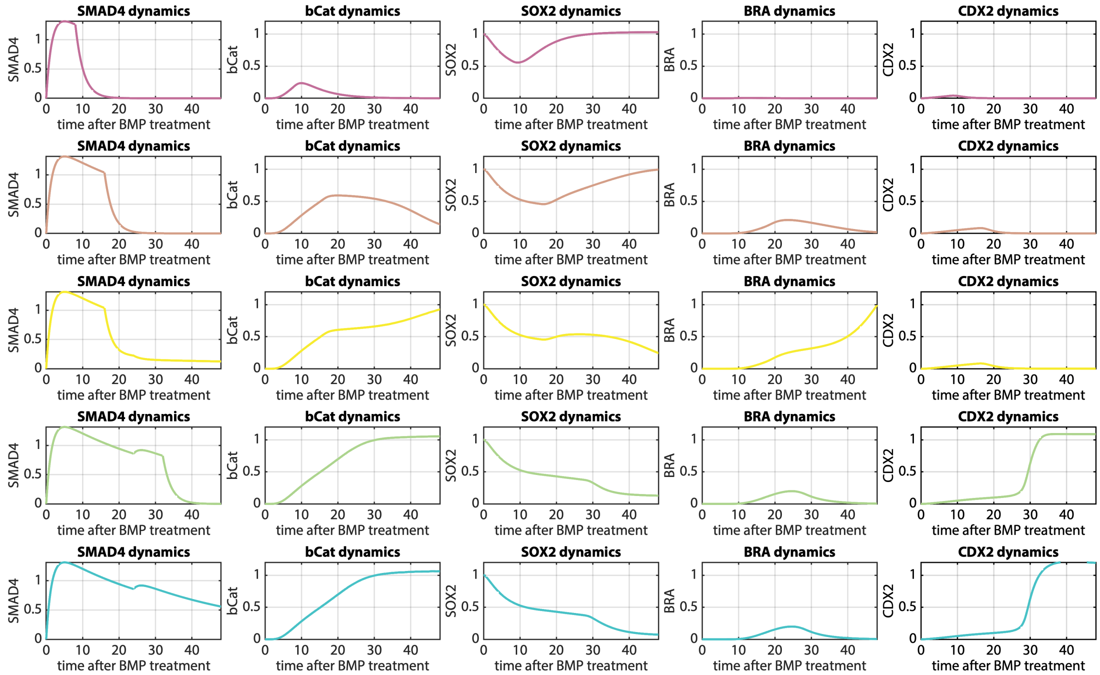

Figure SI6: From top to bottom, simulations of SMAD4, bCat, SOX2, BRA, and CDX2 dynamics for the conditions: BMP 10ng/ml 0-8h, Noggin 8-48h; BMP 10ng/ml 0-16h, Noggin 16-48h; BMP 10ng/ml 0-16h, mTeSR 16-48h; BMP 10ng/ml 0-32h, Noggin 32-48h; BMP 10ng/ml 0-48h using parameters in Tables SI2,SI3 and SI4

where $\omega_i$ are weights and are chosen as $\omega_S = 2$, $\omega_B = 3$, $\omega_C = 2$, $\omega_M = 2$ and $\omega_0 = 3$, and $N_i$ is equal to the number of experimental conditions in $\Omega_i$.

We implemented a Markov Chain Monte Carlo algorithm to explore the parameter space and find a minimum of the cost function in Eq. (7).

Choosing an initial set of parameter values, which set an initial cost function value $L(\boldsymbol{\theta}^0)$, at

| Exp. set | Exp. condition | Simplified data | | |
|---|---|---|---|---|
| | | SOX2 | BRA | CDX2 |
| $\Omega_S$ | @48h, BMP 2ng/ml 0-6, Noggin 6-48 | 1 | 0 | 0 |
| | @48h, BMP 2ng/ml 0-12, Noggin 12-48 | 1 | 0 | 0 |
| | @48h, BMP 2ng/ml 0-18, Noggin 18-48 | 1 | 0 | 0 |
| | @48h, BMP 2ng/ml 0-24, Noggin 24-48 | 1 | 0 | 0 |
| | @48h, BMP 2ng/ml 0-30, Noggin 30-48 | 1 | 0 | 0 |
| | @48h, BMP 2ng/ml 0-36, Noggin 36-48 | 1 | 0 | 0 |
| | @48h, mTeSR 0-48 | 1 | 0 | 0 |
| | @48h, BMP 10ng/ml 0-12, Noggin 12-48 | 1 | 0 | 0 |
| | @48h, BMP 10ng/ml 0-18, Noggin 18-48 | 1 | 0 | 0 |
| | @48h, BMP 10ng/ml 0-24, Noggin 24-48 | 1 | 0 | 0 |
| | @48h, BMP 50ng/ml 0-6, Noggin 6-48 | 1 | 0 | 0 |
| | @48h, BMP 50ng/ml 0-12, Noggin 12-48 | 1 | 0 | 0 |
| | @48h, BMP 50ng/ml 0-18, Noggin 18-48 | 1 | 0 | 0 |
| | @48h, BMP 50ng/ml 0-24, Noggin 24-48 | 1 | 0 | 0 |
| | @48h, BMP 1ng/ml 0-48 | 1 | 0 | 0 |
| | @48h, BMP 1.5ng/ml 0-48 | 1 | 0 | 0 |
| | @48h, BMP 10ng/ml 0-14, mTeSR 14-48 | 1 | 0 | 0 |
| | @48h, BMP 10ng/ml 0-15, mTeSR 15-48 | 1 | 0 | 0 |
| | @16h, BMP 10ng/ml 0-8, mTeSR 8-48 | 0.85 | 0 | 0 |
| | @24h, BMP 10ng/ml 0-8, mTeSR 8-48 | 0.88 | 0 | 0 |
| | @32h, BMP 10ng/ml 0-8, mTeSR 8-48 | 1 | 0 | 0 |
| | @48h, BMP 10ng/ml 0-8, mTeSR 8-48 | 1 | 0 | 0 |
| | @16h, BMP 10ng/ml 0-16, mTeSR 16-48 | 0.62 | 0 | 0 |
| | @24h, BMP 10ng/ml 0-16, mTeSR 16-48 | 0.7 | 0 | 0 |
| | @32h, BMP 10ng/ml 0-16, mTeSR 16-48 | 0.56 | 0 | 0 |
| | @16h, BMP 10ng/ml 0-24, mTeSR 24-48 | 0.62 | 0 | 0 |
| | @24h, BMP 10ng/ml 0-24, mTeSR 24-48 | 0.51 | 0 | 0 |
| | @16h, BMP 10ng/ml 0-32, mTeSR 32-48 | 0.62 | 0 | 0 |
| | @24h, BMP 10ng/ml 0-32, mTeSR 32-48 | 0.51 | 0 | 0 |
| | @16h, BMP 10ng/ml 0-48 | 0.62 | 0 | 0 |
| | @24h, BMP 10ng/ml 0-48 | 0.51 | 0 | 0 |
| | @16h, mTeSR 0-48 | 1 | 0 | 0 |
| | @24h, mTeSR 0-48 | 1 | 0 | 0 |
| | @32h, mTeSR 0-48 | 1 | 0 | 0 |

Table SI5: Table of fitted conditions in which the predominant cell type present was the pluripotent state.

| Exp. set | Exp. condition | Simplified data | | |
|---|---|---|---|---|
| | | SOX2 | BRA | CDX2 |
| $\Omega_B$ | @48h, BMP 10ng/ml 0-30, Noggin 30-48 | 0 | 1 | 0 |
| | @48h, BMP 50ng/ml 0-30, Noggin 30-48 | 0 | 1 | 0 |
| | @48h, BMP 10ng/ml 0-16, mTeSR 16-48 | 0 | 1 | 0 |
| $\Omega_M$ | @32h, BMP 10ng/ml 0-24, mTeSR 24-48 | 0.5 | 0.5 | 0.5 |
| | @48h, BMP 10ng/ml 0-17, mTeSR 17-48 | 0.5 | 0.5 | 0.5 |
| | @48h, BMP 10ng/ml 0-20, mTeSR 20-48 | 0.5 | 0.5 | 0.5 |
| $\Omega_0$ | @48h, BMP 10ng/ml+IWP2 0-48 | 0 | 0 | 1 |
| | @72h, CHIR 0-48 | 0 | 1 | 0 |

Table SI6: Table of fitted conditions in which the predominant cell types present were the mesoderm ($\Omega_B$) or mixed ($\Omega_M$) state, and control conditions ($\Omega_0$).

| Exp. set | Exp. condition | Simplified data | | |
|---|---|---|---|---|
| | | SOX2 | BRA | CDX2 |
| $\Omega_C$ | @48h, BMP 2ng/ml 0-42, Noggin 42-48 | 0 | 0 | 1 |
| | @48h, BMP 10ng/ml 0-36, Noggin 36-48 | 0 | 0 | 1 |
| | @48h, BMP 10ng/ml 0-42, Noggin 42-48 | 0 | 0 | 1 |
| | @48h, BMP 10ng/ml 0-48 | 0 | 0 | 1 |
| | @48h, BMP 50ng/ml 0-30, Noggin 30-48 | 0 | 0 | 1 |
| | @48h, BMP 50ng/ml 0-36, Noggin 36-48 | 0 | 0 | 1 |
| | @48h, BMP 50ng/ml 0-42, Noggin 42-48 | 0 | 0 | 1 |
| | @48h, BMP 50ng/ml 0-48 | 0 | 0 | 1 |
| | @48h, BMP 2ng/ml 0-48 | 0 | 0 | 1 |
| | @48h, BMP 2.5ng/ml 0-48 | 0 | 0 | 1 |
| | @48h, BMP 3ng/ml 0-48 | 0 | 0 | 1 |
| | @48h, BMP 3.5ng/ml 0-48 | 0 | 0 | 1 |
| | @48h, BMP 4ng/ml 0-48 | 0 | 0 | 1 |
| | @48h, BMP 10ng/ml 0-24, mTeSR 24-48 | 0 | 0 | 1 |
| | @32h, BMP 10ng/ml 0-32, mTeSR 32-48 | 0 | 0 | 1 |
| | @48h, BMP 10ng/ml 0-32, mTeSR 32-48 | 0 | 0 | 0.73 |
| | @32h, BMP 10ng/ml 0-48 | 0 | 0 | 0.73 |

Table SI7: Table of fitted conditions in which the predominant cell type present was the ExE state ($\Omega_C$).

each step $n \geq 1$ of the algorithm, the previous accepted cost $\boldsymbol{\theta}^{n-1}$ was perturbed to generate a new candidate parameter vector $\boldsymbol{\theta}^{n,*}$, where each component $\theta_i^{n,*} \sim \mathcal{N}(\theta_i^{n-1}, 0.001|\theta_i^0|)$, i.e. $\theta_i^{n,*}$ follows a normal distribution with mean $\theta_i^{n-1}$ and variance $0.001|\theta_i^0|$. Now, if the cost function evaluated at the candidate parameter $L(\boldsymbol{\theta}^{n,*})$ was smaller than the cost function evaluated at the previous accepted parameter $L(\boldsymbol{\theta}^{n-1})$, the candidate parameter would be accepted and $\boldsymbol{\theta}^n = \boldsymbol{\theta}^{n,*}$, otherwise a new candidate parameter would be drawn.

The positive control conditions for each cell type (see Table SI1) were included in the training data. The model was required to have an attractor representing each cell type in its corresponding positive control condition. For speed, we evaluated these conditions first and immediately discarded parameter sets that lacked the appropriate fixed point in any positive control condition as follows. For each candidate parameter value $\boldsymbol{\theta}^{n,*}$, we simulated the experimental conditions mTeSR 0-48h (which should converge to a SOX2+ state); BMP 10ng/ml 0-16, mTeSR 16-48h (which should converge to a BRA+ state); and BMP 10ng/ml 0-48h (which should converge to a CDX2+ state). We used the end points of each of the three trajectories as initial conditions to approximate the attractors under fixed signals (SMAD4 = 0, $\beta$Cat=0) (which should contain a SOX2+ attractor), (SMAD4 = 0, $\beta$Cat=1) (which should contain a BRA+ attractor), and (SMAD4 = 1, $\beta$Cat=1) (which should contain a CDX2+ attractor), respectively. If one of these attractors did not exist, then the parameter was inconsistent with our assumptions, and it was removed from consideration by setting its cost function to

$$L(\boldsymbol{\theta}) = \frac{1}{\sum_{i\in\{S,C,B,M,0\}} \omega_i} \sum_{i\in\{S,C,B,M,0\}} \frac{\omega_i}{N_i} \sum_{c\in\Omega_i} \|\boldsymbol{x}_c^{exp}(\boldsymbol{\theta})\|^2. \tag{8}$$

##### 2.3.3 Fitting results

We first fitted the CFN model assuming the WNT signaling model followed the dynamics of a slightly different ODE system to the one introduced in Eqs. 9, where CHIR affected WNT dynamics instead of the dynamics of $\beta$Cat:

$$\begin{cases} \dfrac{d\text{WNT}}{dt} = (1-\text{IWP2})\left(a_{SW}\dfrac{\text{SMAD4}^{n_{SW}}}{K_{SW}^{n_{SW}}+\text{SMAD4}^{n_{SW}}} + a_{WW}\dfrac{\beta\text{Cat}^{n_{WW}}}{K_{WW}^{n_{WW}}+\beta\text{Cat}^{n_{WW}}}\right) + a_{C\beta}\text{CHIR} - \delta_W\text{WNT} \\ \dfrac{d\beta\text{Cat}}{dt} = a_{W\beta}\dfrac{\text{WNT}^{n_{W\beta}}}{K_{W\beta}^{n_{W\beta}}+\text{WNT}^{n_{W\beta}}} - \delta_\beta\beta\text{Cat}, \end{cases} \tag{9}$$

We fixed the values of the parameters of the signaling model in Tables SI2 and SI3, except $a_{C\beta}$, which was set to $a_{C\beta} = 0.15$. We fitted the values of the parameters of the CFN model, plus the value $r_b$ which gives the proportion of original BMP that is washed out by an mTeSR media change. We chose $r_b = 0.1$ as its initial value.

The initial cost value was equal to $L(\boldsymbol{\theta}^0) = 0.9843$. After $5,668,950$ iterations of the MCMC algorithm, $7,403$ parameter values were accepted, and the evolution of the costs is shown in Fig. SI7. The last accepted parameter set, which had the lowest cost value equal to 0.0816, is given in Table SI8.

Following these optimizations, we realized that it was more biologically accurate to consider CHIR as a factor changing $\beta$Cat dynamics directly, rather than through the WNT variable, as shown in Eq. (4). With the fitted parameters for the CFN, changing the WNT model from the one shown in Eq. (9) to Eq. (4) we found a value for $a_{C\beta}$ that made the CFN dynamics practically equivalent to the ones fitted. With this new model and $a_{C\beta} = 0.8$, the cost function for the initial CFN parameters in Table SI4 was 0.9839, while for the final fitted CFN parameters in Table SI8 was 0.0857. The simulations of the fitted conditions with the final parameters are shown in Figures SI8 to SI15.

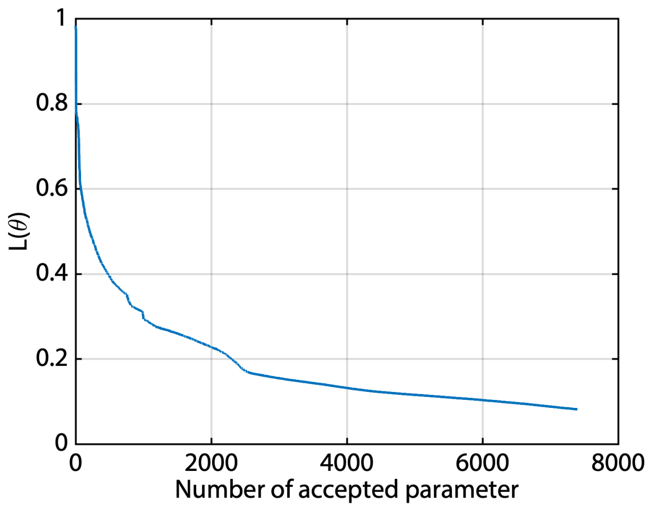

Figure SI7: Evolution of the cost function for every accepted parameter.

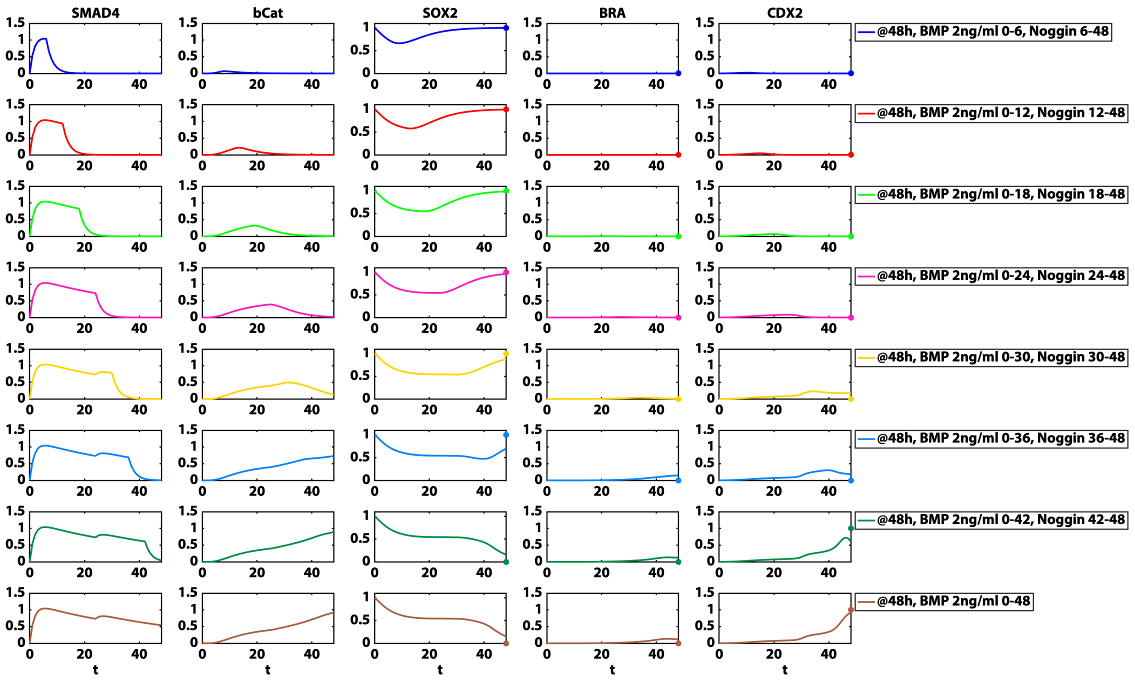

Figure SI8: From top to bottom, simulations of SMAD4, bCat, SOX2, BRA, and CDX2 dynamics for the indicated conditions using parameters in Tables SI2,SI3 and SI8. Experimental data point to be fitted is shown as a colored dot.

| Parameter | Value |
|---|---|
| $a_S$ | 0.1695 |
| $n_{CS}$ | 5.095 |
| $K_{CS}$ | 0.3426 |
| $n_{BS}$ | 3.7226 |
| $K_{BS}$ | 0.4050 |
| $n_{SmS}$ | 0.4868 |
| $K_{SmS}$ | 1.0644 |
| $\delta_S$ | 0.1639 |

| Parameter | Value |
|---|---|
| $a_B$ | 0.7833 |
| $n_{\beta B}$ | 4.8778 |
| $K_{\beta B}$ | 0.8056 |
| $n_{SB}$ | 2.1554 |
| $K_{SB}$ | 0.2726 |
| $n_{CB}$ | 1.8623 |
| $K_{CB}$ | 0.1348 |
| $\delta_B$ | 0.1705 |

| Parameter | Value |
|---|---|
| $a_C$ | 0.4239 |
| $n_{SmC}$ | 2.4569 |
| $K_{SmC}$ | 0.0680 |
| $n_{SC}$ | 5.3994 |
| $K_{SC}$ | 0.3470 |
| $n_{BC}$ | 3.9198 |
| $K_{BC}$ | 0.2856 |
| $a_{CC}$ | 0.0922 |
| $n_{CC}$ | 6.1301 |
| $K_{CC}$ | 0.1209 |
| $\delta_B$ | 0.4854 |

Table SI8: Table of parameter values which best fitted the data resulting from the MCMC algorithm.

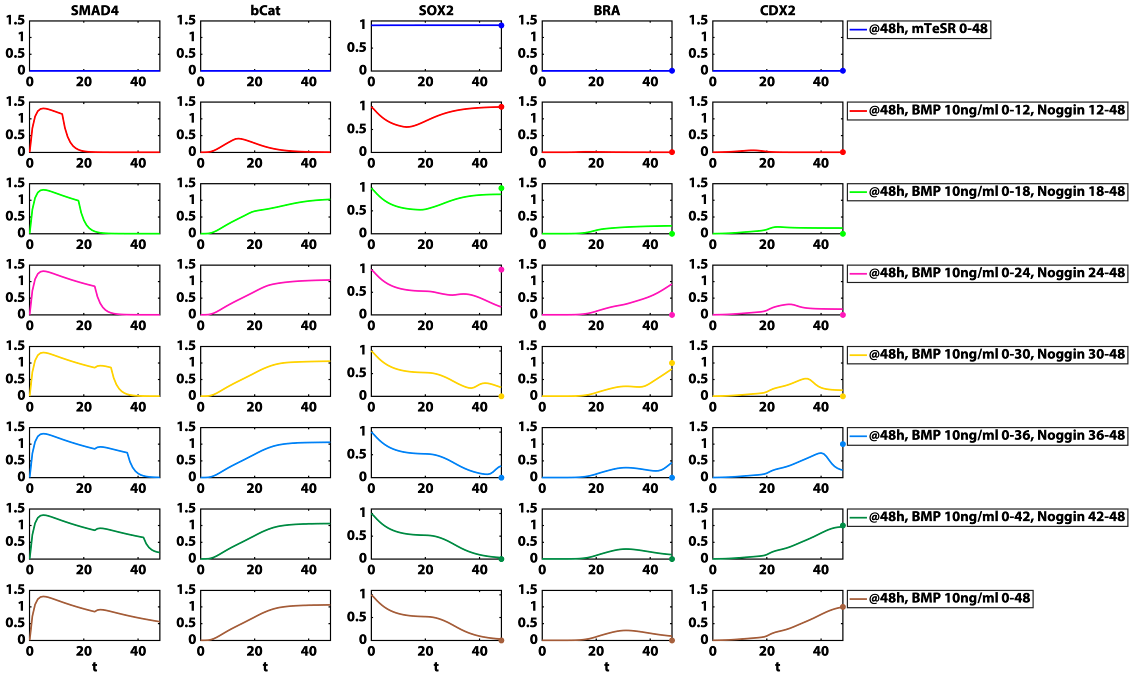

Figure SI9: From top to bottom, simulations of SMAD4, bCat, SOX2, BRA, and CDX2 dynamics for the indicated conditions using parameters in Tables SI2,SI3 and SI8. Experimental data point to be fitted is shown as a colored dot.

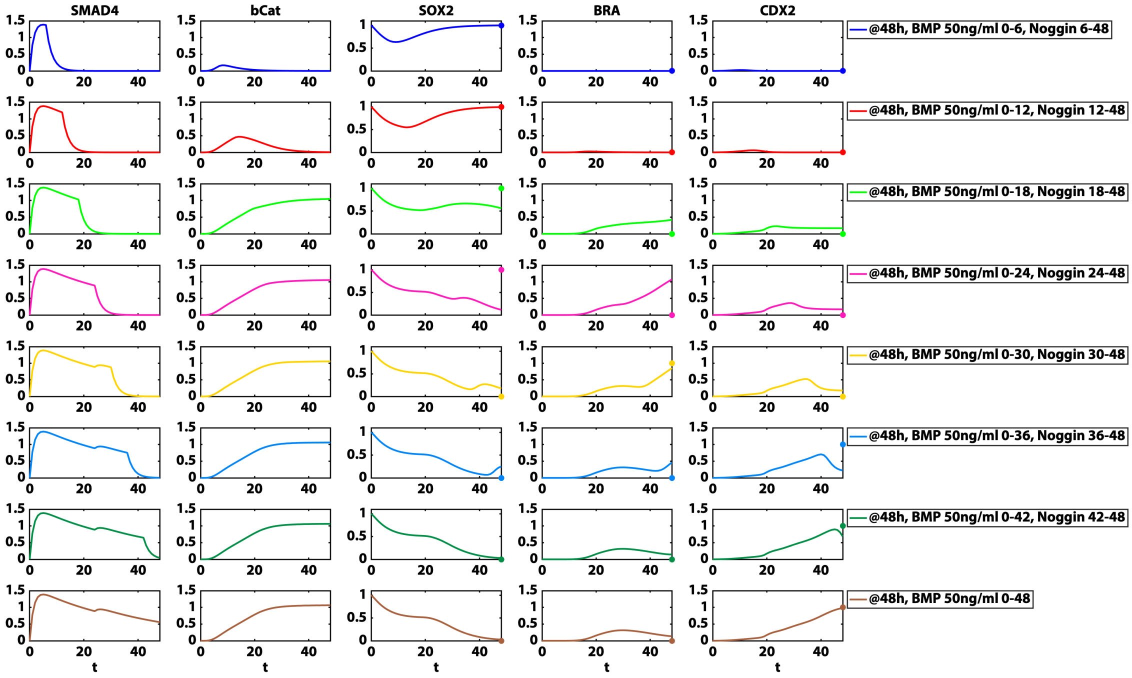

Figure SI10: From top to bottom, simulations of SMAD4, bCat, SOX2, BRA, and CDX2 dynamics for the indicated conditions using parameters in Tables SI2,SI3 and SI8. Experimental data point to be fitted is shown as a colored dot.

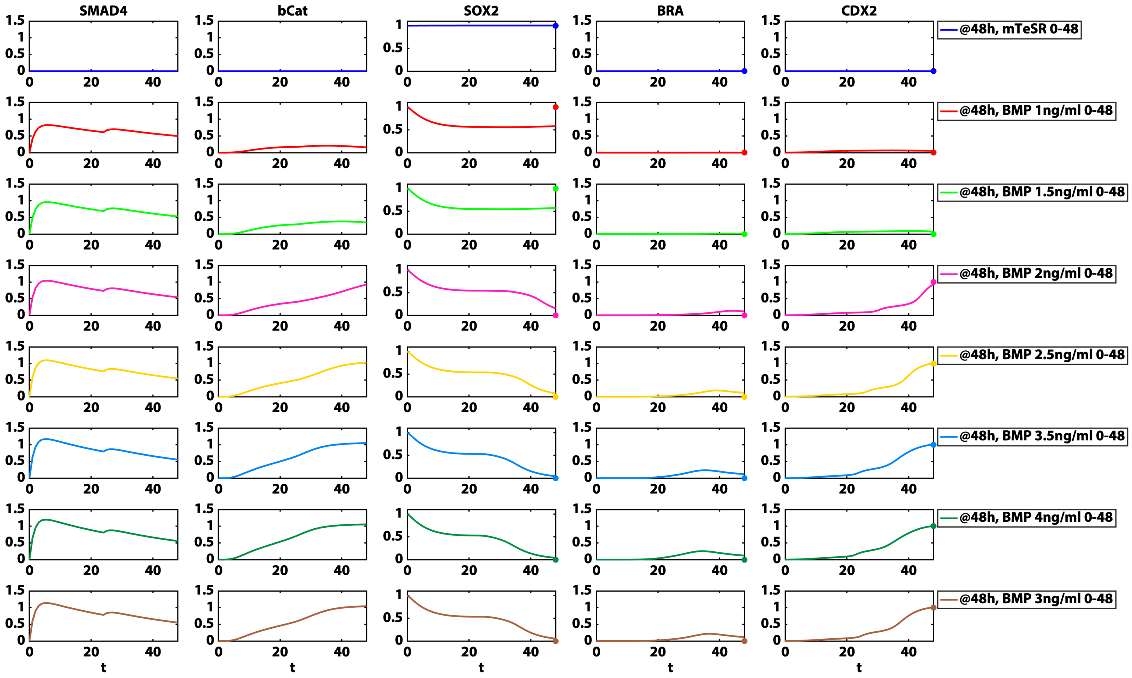

Figure SI11: From top to bottom, simulations of SMAD4, bCat, SOX2, BRA, and CDX2 dynamics for the indicated conditions using parameters in Tables SI2,SI3 and SI8. Experimental data point to be fitted is shown as a colored dot.

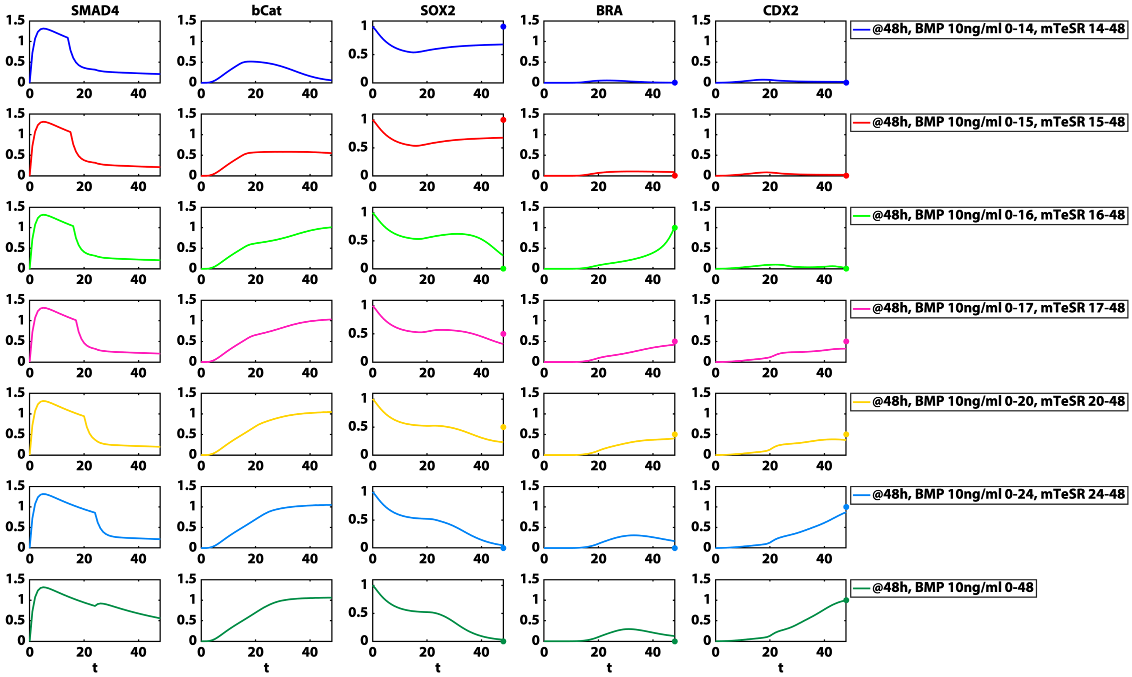

Figure SI12: From top to bottom, simulations of SMAD4, bCat, SOX2, BRA, and CDX2 dynamics for the indicated conditions using parameters in Tables SI2,SI3 and SI8. Experimental data point to be fitted is shown as a colored dot.

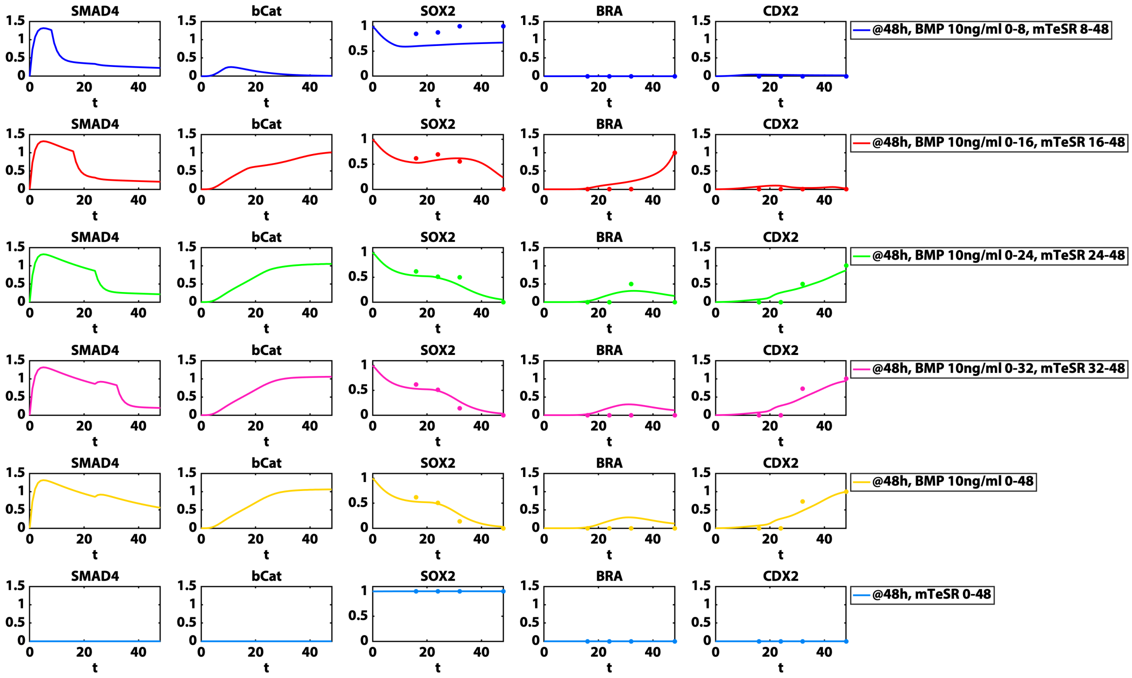

Figure SI13: From top to bottom, simulations of SMAD4, bCat, SOX2, BRA, and CDX2 dynamics for the indicated conditions using parameters in Tables SI2,SI3 and SI8. Experimental data point to be fitted is shown as a colored dot.

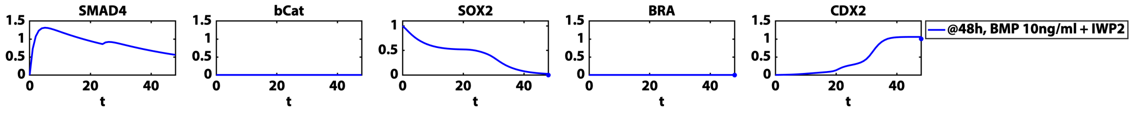

Figure SI14: Simulation of SMAD4, bCat, SOX2, BRA, and CDX2 dynamics for the indicated condition using parameters in Tables SI2,SI3 and SI8. Experimental data point to be fitted is shown as a colored dot.

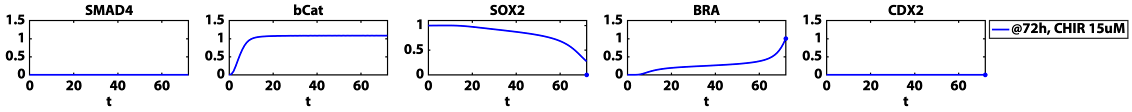

Figure SI15: Simulation of SMAD4, bCat, SOX2, BRA, and CDX2 dynamics for the indicated condition using parameters in Tables SI2,SI3 and SI8. Experimental data point to be fitted is shown as a colored dot.

##### 2.3.4 Fate map

Given the fitted parameter values, we computed a fate map that shows the attractors present for each value of SMAD4 and $\beta$Cat.

Given fixed values for SMAD4 = SMAD4$_i$ and a $\beta$Cat = $\beta$Cat$_j$, we compute the attractors that are present in the CFN for that signaling profile by exploring the different states that the CFN can evolve towards towards using a defined set of initial conditions as follows.

Using the three reference SOX2+, BRA+, CDX2+ attractors we introduced above, we create a grid in the state space. This discretization of the state space was denser on the axes, since due to the mutual inhibition interactions, the critical points tend to be near the axes. For each point in this grid (SOX2$_k$, BRA$_l$, CDX2$_m$), we study the value towards which the CFN system will tend to if this point is used as initial condition. To do so, we solve the CFN system with SMAD4 = SMAD4$_0$, $\beta$Cat = $\beta$Cat$_0$, using initial condition (SOX2$_k$, BRA$_l$, CDX2$_m$) in $t \in (0 \ldots, 1,000)$. If the final point of this trajectory at $t = 1,000$ is real and non-negative, we compute the stability of the point given by the eigenvalues of the jacobian matrix of the CFN system with SMAD4 = SMAD4$_i$, $\beta$Cat = $\beta$Cat$_j$ evaluated at the point. If the point is an attractor, it is classified as representing the pluripotent, mesoderm or ExE state depending on the distance to the reference attractors.

We repeat this procedure for 90601 pairs of (SMAD4$_i$, $\beta$Cat$_j$) values, where SMAD4 = SMAD4$_i$ $\in \{0, 0.005, 0.01, \ldots 1.5\}$ value, and a $\beta$Cat = $\beta$Cat$_j$ $\in \{0, 0.005, 0.01, \ldots 1.5\}$ to create the plot in Figure 7. The discretization of the (SOX2, BRA, CDX2) space contained 121 points: 64 points were obtained from making a grid containing all the possible combinations of equally spaced values where SOX2$_k$ $\in (0, SOX2^*)$, $k = 1, 2, 3, 4$, BRAl$\in (0, BRA^*)$, $l = 1, 2, 3, 4$, CDX2$_m$ $\in (0, CDX2^*)$, $m = 1, 2, 3, 4$, where $SOX2^*$, $BRA^*$, $CDX2^*$ are the SOX2, BRA, and CDX2 values of the three reference SOX2+, BRA+, CDX2+ attractors, respectively; 11 points had the form (SOX2$_k$, 0, 0) where SOX2$_k$ $\in \{0, 0.1, 0.2, 0.3, , SOX2^*\}$, where $SOX2^*$ is the SOX2 value of the reference SOX2+ attractor; 35 points had the form (0, BRA$_l$, 0) where BRA$_l$ $\in \{0, 0.1, 0.2, 0.3, , BRA^*\}$, where $BRA^*$ is the BRA value of the reference BRA+ attractor; 11 points had the form (0, 0, CDX2$_m$) where CDX2$_m$ $\in \{0, 0.1, 0.2, 0.3, , CDX2^*\}$, where $CDX2^*$ is the CDX2 value of the reference CDX2+ attractor.

#### 2.4 Stochastic CFN model

Having a deterministic CFN model that fitted the simplified experimental data, our goal was to obtain a model that could recapitulate the proportions of cell types observed in the data. For this purpose, we developed a stochastic version of the described CFN model in Eq. 6 by adding a white noise $\eta$ to each equation:

$$\begin{cases} \dfrac{d\text{SOX2}}{dt} = \dfrac{a_S}{(K_{SmS}^{n_{SmS}} + \text{SMAD4}^{n_{SmS}})(1 + (\text{CDX2}/K_{CS})^{n_{CS}} + (\text{BRA}/K_{BS})^{n_{BS}})} - \delta_S\text{SOX2} + \eta_1(t) \\ \dfrac{d\text{BRA}}{dt} = \dfrac{a_B\beta Cat^{n_{\beta B}}}{(K_{\beta B}^{n_{\beta B}} + \beta Cat^{n_{\beta B}})(1 + (\text{SOX2}/K_{SB})^{n_{SB}} + (\text{CDX2}/K_{CB})^{n_{CB}})} - \delta_B\text{BRA} + \eta_2(t) \\ \dfrac{d\text{CDX2}}{dt} = \dfrac{a_C\text{SMAD4}^{n_{SmC}}}{(K_{SmC}^{n_{SmC}} + \text{SMAD4}^{n_{SmC}})(1 + (\text{SOX2}/K_{SC})^{n_{SC}} + (\text{BRA}/K_{BC})^{n_{BC}})} + \dots \\ \qquad \dots + \dfrac{a_{CC}\text{CDX2}^{n_{CC}}}{K_{CC}^{n_{CC}} + \text{CDX2}^{n_{CC}}} - \delta_C\text{CDX2} + \eta_3(t). \end{cases} \tag{10}$$

where $n_i(t)$ is drawn from a Gaussian distribution with variance $\xi\Delta t$. The signaling system was kept deterministic as we only wanted to consider intrinsic noise.

To simulate the SDE system in Eq. 10 we took advantage of the Euler-Maruyama method, as in [5, 6, 7, 8]. For a given parameter set and a noise variance, we draw $N$ initial conditions from an initial distribution in (SOX2, BRA, CDX2) space and simulate $N$ random walks with the Euler Maruyama method. We classify the end points of the $N$ trajectories into the same categories as the experimental data using the unitary coordinates of these points in the coordinate basis given by the attractors of the deterministic CFN model computed as explained in Section 2.3.2, and following the same criteria as with the experimental data. This way, given a parameter set and a noise variance, we can simulate the fate proportions from the model that can be compared to the experimental data.

Using this simulation procedure, fixing the parameters fitted with the deterministic model, $N = 1,000$ and $\Delta t = 0.001$, we fit the magnitude of the noise $\xi$, and the mean and covariance matrix of the distribution of initial conditions $(\mu_0, 0, 0)$ and $\rho_0\Sigma$ using the experimental cell type proportions from a subset of conditions. We use a similar approach as with the ODE and leverage a similar MCMC algorithm but with a different cost function. In this case, the experimental and simulated cell type proportions are compared instead, as shown in Eq. 11. Let $\boldsymbol{x}_c^{exp} = (P1_c^{exp}, P2_c^{exp}, P3_c^{exp}, P4_c^{exp}, P5_c^{exp}, P6_c^{exp}, P7_c^{exp}, P8_c^{exp})$, be the vector of cell type proportions experimentally obtained under experimental condition $c$, where $Pi_c^{exp}$ is the proportion of cells that are classified as All low, SOX2+, BRA+, CDX2+, SOX2/BRA+, BRA/CDX2+, SOX2/CDX2+, and Mixed when $i = 1, \dots, 8$, respectively, and $\boldsymbol{x}_c^{num}(\boldsymbol{\theta}) = (P1_c^{num}, P2_c^{num}, P3_c^{num}, P4_c^{num}, P5_c^{num}, P6_c^{num}, P7_c^{num}, P8_c^{num})$ be the corresponding simulated proportions with parameter $\boldsymbol{\theta}$. The cost function for parameter $\boldsymbol{\theta}$ is given by:

$$L(\boldsymbol{\theta}) = \sum_{c\in\Omega_i} \|\boldsymbol{x}_c^{exp} - \boldsymbol{x}_c^{num}(\boldsymbol{\theta})\|^2. \tag{11}$$

This time, as the fitting procedure will fix the CFN parameters and only fit the magnitude of the noise $\xi$, and the mean and covariance matrix of the distribution of initial conditions $\mu_0$ and $\Sigma_0$, the 3 attractors corresponding to the SOX2+, BRA+, and CDX2+ states can always be defined. Therefore, the given definition of the cost function is sufficient.

The data set used for the fitting of the noise and distribution of the initial condition is given in Table SI9, and contained a subset of the conditions previously used for fitting the deterministic CFN.

We ran the MCMC algorithm with initial candidate noise magnitude equal to $\xi_0 = 0.0001$, and initial conditions drawn from a normal distribution with mean $\mu_0 = 1$, and variance matrix equal to $\rho_0 = 1$, $\Sigma = (0.0848, 0.0072, 0.0011; 0.0072, 0.001, 0.0002; 0.0011, 0.0002, 0.0004)$, obtained from the experimental data. At each step of the MCMC algorithm, each of this three parameters was multiplied by a number drawn from a normal distribution with mean given by the previous accepted parameter set and variances equal to 0.000001, 0.01 and 0.01, respectively. The candidate parameter set was accepted if the cost function was less than the cost defined by the previous accepted parameter value.

After $2,650$ steps in the MCMC algorithm, 10 parameter values were accepted, we considered that the trends in the experimental and simulated cell type proportions were sufficiently similar, and the

| Exp. condition | Experimental cell type proportions | | | | | | | |
|---|---|---|---|---|---|---|---|---|
| | AL | S+ | SB+ | B+ | BC+ | C+ | SC+ | M |
| @48h, mTeSR 0-48h | 0.0055 | 0.9837 | 0 | 0 | 0.0002 | 0.0027 | 0.0017 | 0.0061 |
| @48h, BMP 10ng/ml 0-4, mTeSR 4-48 | 0.0086 | 0.9805 | 0 | 0 | 0.0014 | 0.002 | 0.002 | 0.0053 |
| @48h, BMP 10ng/ml 0-8, mTeSR 8-48 | 0.0101 | 0.9717 | 0.0008 | 0.0003 | 0.0007 | 0.0081 | 0.0017 | 0.0066 |
| @48h, BMP 10ng/ml 0-16, mTeSR 16-48 | 0.0173 | 0.1937 | 0.105 | 0.2894 | 0.0124 | 0.0258 | 0.0109 | 0.3454 |
| @48h, BMP 10ng/ml 0-24, mTeSR 24-48 | 0.0319 | 0.0004 | 0.0001 | 0.0541 | 0.0352 | 0.8605 | 0.0007 | 0.0171 |
| @48h, BMP 10ng/ml 0-32, mTeSR 32-48 | 0.0366 | 0.0011 | 0 | 0.013 | 0.0113 | 0.9156 | 0.0068 | 0.0157 |
| @48h, BMP 10ng/ml 0-40, mTeSR 40-48 | 0.0483 | 0.0004 | 0 | 0.0048 | 0.0062 | 0.9394 | 0 | 0.0009 |
| @48h, BMP 10ng/ml 0-48 | 0.0182 | 0 | 0 | 0.0039 | 0.0062 | 0.9706 | 0.0001 | 0.001 |
| @48h, BMP 50ng/ml 0-6, Noggin 6-48 | 0.0059 | 0.9853 | 0.004 | 0.004 | 0 | 0 | 0 | 0.0009 |
| @48h, BMP 50ng/ml 0-12, Noggin 12-48 | 0.0032 | 0.9745 | 0.0097 | 0.0109 | 0 | 0 | 0 | 0.0016 |
| @48h, BMP 50ng/ml 0-18, Noggin 18-48 | 0.0038 | 0.9287 | 0.0276 | 0.0291 | 0.0006 | 0.0008 | 0.0008 | 0.0086 |
| @48h, BMP 50ng/ml 0-24, Noggin 24-48 | 0.007 | 0.8966 | 0.0324 | 0.0547 | 0.0003 | 0.0006 | 0 | 0.0084 |
| @48h, BMP 50ng/ml 0-30, Noggin 30-48 | 0.022 | 0.2704 | 0.0163 | 0.3068 | 0.043 | 0.0916 | 0.0486 | 0.2012 |
| @48h, BMP 50ng/ml 0-36, Noggin 36-48 | 0.0251 | 0.004 | 0 | 0.0293 | 0.0199 | 0.8717 | 0.0176 | 0.0324 |
| @48h, BMP 50ng/ml 0-42, Noggin 42-48 | 0.0122 | 0.0001 | 0 | 0.0015 | 0.0003 | 0.9808 | 0.0001 | 0.005 |
| @48h, BMP 50ng/ml 0-48 | 0.0107 | 0 | 0 | 0.0027 | 0.0006 | 0.9824 | 0 | 0.0036 |
| @48h, BMP 1ng/ml 0-48 | 0.0021 | 0.9943 | 0.0007 | 0.0004 | 0 | 0 | 0 | 0.0025 |
| @48h, BMP 1.5ng/ml 0-48 | 0.0037 | 0.7946 | 0.0169 | 0.0077 | 0 | 0.0199 | 0.0506 | 0.1066 |
| @48h, BMP 2ng/ml 0-48 | 0.0117 | 0.146 | 0.0224 | 0.0458 | 0.0119 | 0.3216 | 0.1496 | 0.2911 |
| @48h, BMP 2.5ng/ml 0-48 | 0.0108 | 0.0056 | 0.0004 | 0.0152 | 0.0227 | 0.7998 | 0.048 | 0.0975 |
| @48h, BMP 3ng/ml 0-48 | 0.0163 | 0.0002 | 0 | 0.0085 | 0.0167 | 0.9347 | 0.0037 | 0.02 |
| @48h, BMP 3.5ng/ml 0-48 | 0.0111 | 0 | 0 | 0.0042 | 0.0133 | 0.9436 | 0.0029 | 0.0249 |
| @48h, BMP 4ng/ml 0-48 | 0.0146 | 0.0005 | 0 | 0.0017 | 0.0148 | 0.9584 | 0.0012 | 0.0088 |
| @48h, BMP 10ng/ml 0-14, mTeSR 14-48 | 0.0028 | 0.8468 | 0.0748 | 0.0544 | 0 | 0.0016 | 0.0017 | 0.018 |
| @48h, BMP 10ng/ml 0-15, mTeSR 15-48 | 0.0044 | 0.4914 | 0.1798 | 0.169 | 0.0046 | 0.0141 | 0.0169 | 0.1198 |
| @48h, BMP 10ng/ml 0-17, mTeSR 17-48 | 0.0074 | 0.255 | 0.1046 | 0.2061 | 0.0138 | 0.0549 | 0.0409 | 0.3173 |
| @48h, BMP 10ng/ml 0-20, mTeSR 20-48 | 0.0106 | 0.1282 | 0.0266 | 0.0907 | 0.0305 | 0.2769 | 0.0618 | 0.3747 |
| @48h, BMP 10ng/ml 0-24, mTeSR 24-48 | 0.0095 | 0.0022 | 0.001 | 0.0467 | 0.0415 | 0.8422 | 0.0077 | 0.0491 |

Table SI9: Table experimental proportions to which the SDE model in Eq. 10 was fitted. AL, S+, SB+, B+, BC+, C+, SC+, M cell types correspond to All Low, SOX2+, SOX2/BRA+, BRA+, BRA/CDX2+, CDX2+, SOX2/CDX2+, Mixed, respectively.

comparisons are shown in Figures SI16 to SI19. Final parameter values were equal to $\xi = 0.0001003$, $\mu = 0.9775$, and $\rho_0 = 0.9437$.

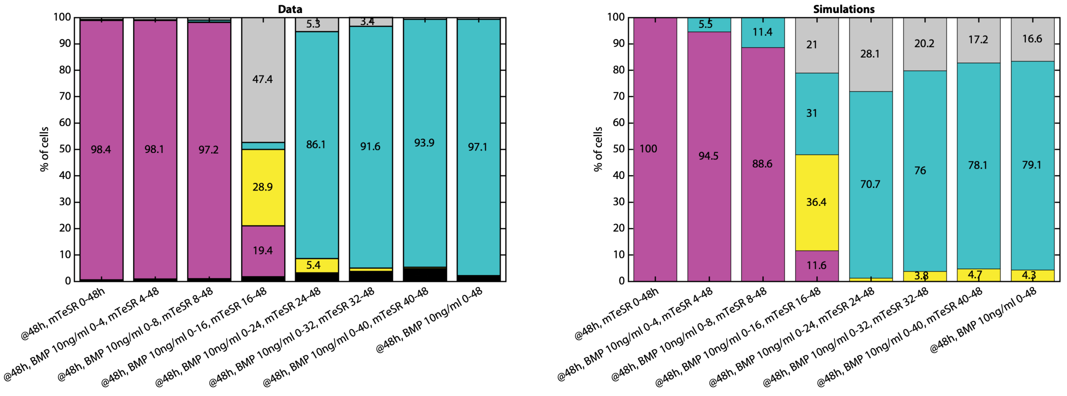

Figure SI16: Experimental (left) and simulated (right) cell fate proportions for the indicated experimental conditions. SOX2+ = magenta, BRA+ = yellow, CDX2+ = cyan, Mixed data (pooled SOX2/BRA+, BRA/CDX2+, SOX2/CDX2+, Mixed) = gray, All low = black.

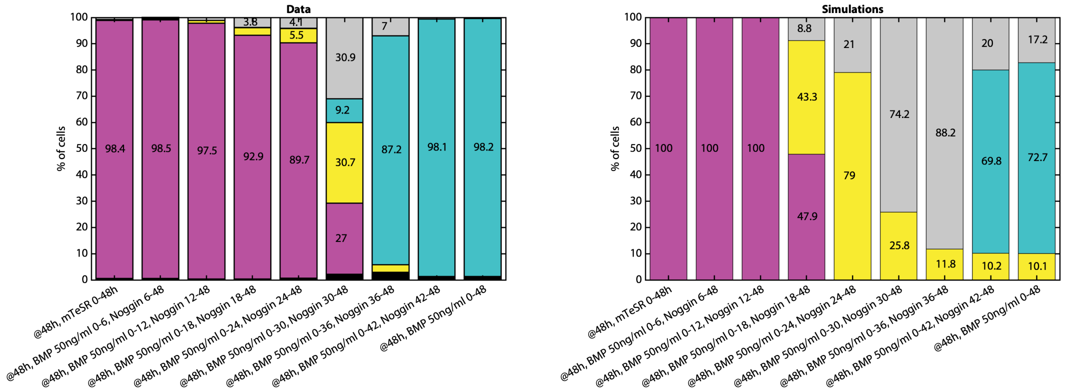

Figure SI17: Experimental (left) and simulated (right) cell fate proportions for the indicated experimental conditions. SOX2+ = magenta, BRA+ = yellow, CDX2+ = cyan, Mixed data (pooled SOX2/BRA+, BRA/CDX2+, SOX2/CDX2+, Mixed) = gray, All low = black.

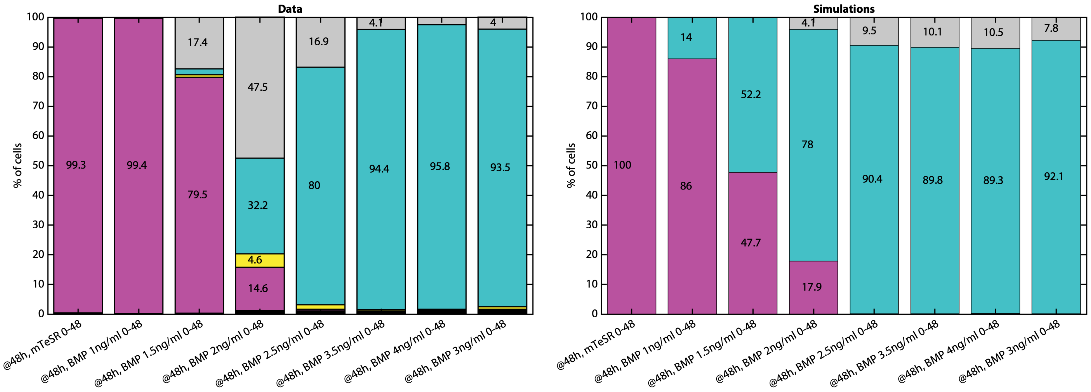

Figure SI18: Experimental (left) and simulated (right) cell fate proportions for the indicated experimental conditions. SOX2+ = magenta, BRA+ = yellow, CDX2+ = cyan, Mixed data (pooled SOX2/BRA+, BRA/CDX2+, SOX2/CDX2+, Mixed) = gray, All low = black.

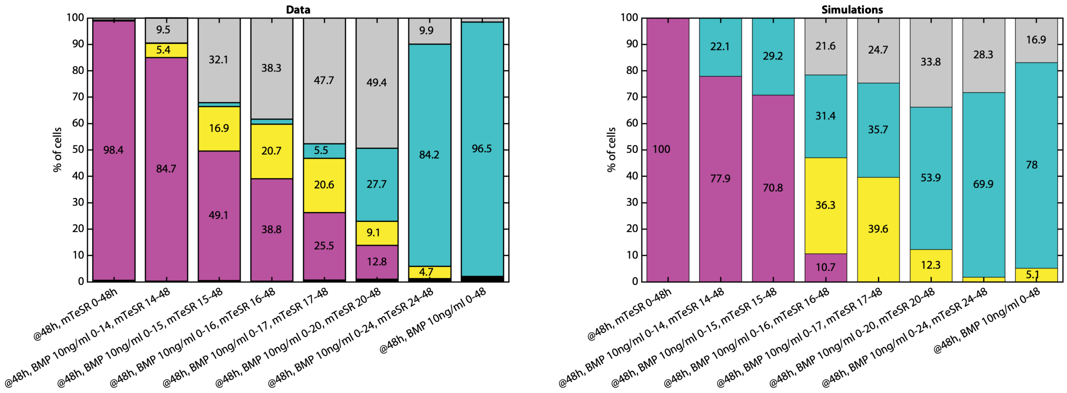

Figure SI19: Experimental (left) and simulated (right) cell fate proportions for the indicated experimental conditions. SOX2+ = magenta, BRA+ = yellow, CDX2+ = cyan, Mixed data (pooled SOX2/BRA+, BRA/CDX2+, SOX2/CDX2+, Mixed) = gray, All low = black.
